# Bright and photostable chemigenetic indicators for extended *in vivo* voltage imaging

**DOI:** 10.1101/436840

**Authors:** Ahmed S. Abdelfattah, Takashi Kawashima, Amrita Singh, Ondrej Novak, Hui Liu, Yichun Shuai, Yi-Chieh Huang, Jonathan B. Grimm, Ronak Patel, Johannes Friedrich, Brett D. Mensh, Liam Paninski, John J. Macklin, Kaspar Podgorski, Bei-Jung Lin, Tsai-Wen Chen, Glenn C. Turner, Zhe Liu, Minoru Koyama, Karel Svoboda, Misha B. Ahrens, Luke D. Lavis, Eric R. Schreiter

**Affiliations:** Janelia Research Campus, Howard Hughes Medical Institute, 19700 Helix Drive, Ashburn, Virginia, USA 20147; The Solomon H. Snyder Department of Neuroscience, The Johns Hopkins University, 725 N. Wolfe Street, Baltimore, MD 21205; Department of Auditory Neuroscience, Institute of Experimental Medicine, Academy of Sciences of the Czech Republic, Prague, Czech Republic; Institute of Neuroscience, National Yang-Ming University, Taipei, 112 Taiwan; Department of Statistics and Center for Theoretical Neuroscience, Columbia University, 3227 Broadway, New York, NY 10027; Department of Neuroscience and Grossman Center for the Statistics of Mind, Columbia University, 3227 Broadway, New York, NY 10027; Center for Computational Biology, Flatiron Institute, 162 5th Ave, New York, NY 10010

## Abstract

Imaging changes in membrane potential using genetically encoded fluorescent voltage indicators (GEVIs) has great potential for monitoring neuronal activity with high spatial and temporal resolution. Brightness and photostability of fluorescent proteins and rhodopsins have limited the utility of existing GEVIs. We engineered a novel GEVI, ‘Voltron’, that utilizes bright and photostable synthetic dyes instead of protein-based fluorophores, extending the combined duration of imaging and number of neurons imaged simultaneously by more than tenfold relative to existing GEVIs. We used Voltron for *in vivo* voltage imaging in mice, zebrafish, and fruit flies. In mouse cortex, Voltron allowed single-trial recording of spikes and subthreshold voltage signals from dozens of neurons simultaneously, over 15 minutes of continuous imaging. In larval zebrafish, Voltron enabled the precise correlation of spike timing with behavior.

---

Animal behavior is produced by patterns of neuronal activity that span a wide range of spatial and temporal scales. To understand how neural circuits mediate behavior thus requires high-speed recording from ensembles of neurons for long periods of time. Although the activity of large numbers of neurons can now be routinely recorded using genetically encoded calcium indicators (GECIs) (*1*), the slow kinetics of calcium signals complicate the measurement of action potentials, and sub-threshold voltage signals are missed entirely (*1–3*). Voltage imaging using genetically encoded voltage indicators (GEVIs) can overcome these challenges, enabling imaging of fast spikes and subthreshold dynamics in genetically defined neurons (*4,5*). The high imaging speed and excitation intensity required for voltage imaging, combined with the smaller volume of the cellular membrane, place increased demands on voltage indicators relative to GECIs. Extant GEVIs rely on fluorescence from either microbial rhodopsins (*6–8*) or fluorescent proteins (FPs) (*9–13*). These fluorophores lack the brightness and photostability to allow *in vivo* voltage imaging from large fields of view over timescales of many behavioral events, precluding the millisecond-timescale interrogation of neural circuits. Recent development of improved rhodamine dyes such as the Janelia Fluor^®^ (JF) dyes enable their use in complex biological experiments due to their high brightness and photostability (*14*), compatibility with self-labeling protein tags (*15,16*), and the ability to traverse the blood-brain barrier for *in vivo* delivery (*17*). Here we describe a ‘chemigenetic’ GEVI scaffold—termed ‘Voltron’—which incorporates these synthetic fluorophore dyes. Voltron provides an increased photon yield that enables *in vivo* imaging of neuronal spiking and sub-threshold voltage signals in model organisms with order-of-magnitude improvement in the number of neurons imaged simultaneously over substantially longer durations.

Our design for a chemigenetic voltage indicator combines a voltage-sensitive microbial rhodopsin domain (*6, 7, 11*) with a self-labeling protein tag domain (**Fig. 1A**) that covalently binds a synthetic fluorophore dye ligand (*14,15*) (**Fig. 1B**), analogous to previously reported voltage indicators using fluorescent proteins (*10, 11, 18*). Transmembrane-voltage-dependent changes in the absorption spectrum of the rhodopsin (*6,19*) reversibly modulate the degree of fluorescence quenching of the nearby bound dye through Förster resonance energy transfer (FRET). We investigated the modularity of this approach, finding that three different rhodopsin domains, QuasAr1 (*7*), QuasAr2 (*7*), and Ace2N (*11,20*), were all able to modulate the fluorescence of JF_549_ bound to either HaloTag (*15*) or SNAP-tag (*21*) self-labeling tag domains (**Fig. S1** **to S8**). Removing a modest number of amino acid residues at the junction of the rhodopsin and self-labeling tag domains increased the amplitude of fluorescent voltage signals (**Fig. S1**), presumably by decreasing average distance and thus increasing FRET efficiency between the dye and rhodopsin retinal cofactor. The configuration providing the best signal-to-noise ratio for spikes was Ace2N fused to HaloTag with five amino acids removed at their junction (**Figs. 1A,B** and **S2**), hereafter referred to as Voltron.

**Fig. 1.**
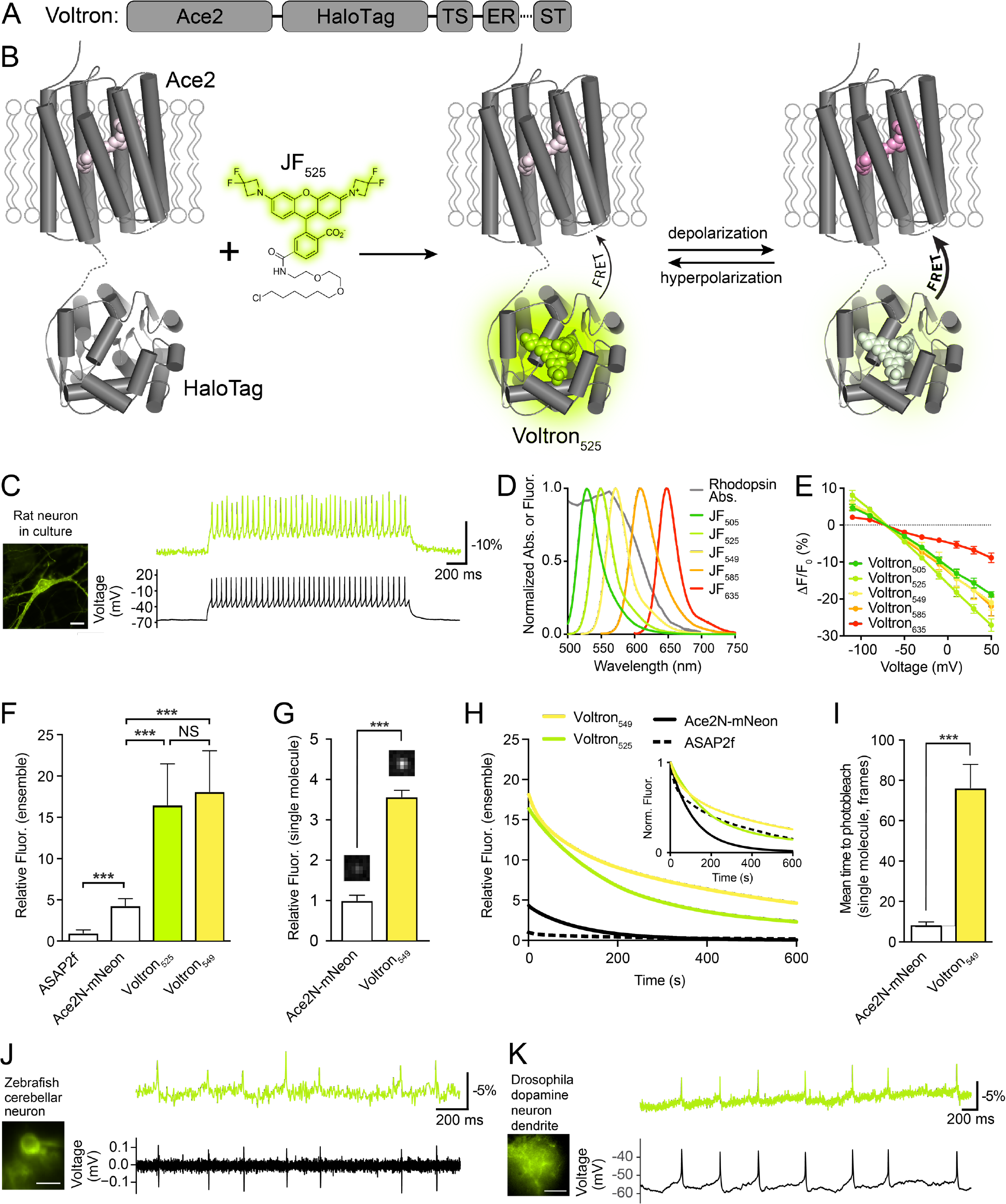
Development of the chemigenetic voltage indicator Voltron. (**A**) Schematic of Voltron sequence: A rhodopsin (Ace2) is fused to a self-labeling tag domain (HaloTag) with additional sequences added to improve or localize membrane targeting: endoplasmic reticulum export sequence (ER), Golgi export trafficking sequence (TS), and somatic targeting sequence (ST). (**B**) Model of Voltron mechanism. The HaloTag domain of the transmembrane Voltron protein (grey cylinders) covalently binds a small molecule fluorophore such as JF_525_ (green glow) with an appended HaloTag ligand. Membrane depolarization reversibly decreases JF_525_ fluorescence via increased FRET to the rhodopsin domain. (**C**) Left panel: cultured rat hippocampal neuron expressing Voltron and labeled with JF_525_. Scale bar: 20 μm. Right panel: single-trial recording of action potentials and subthreshold voltage signals from current injections in primary neuron culture using imaging (top, fluorescence) or electrophysiology (bottom, membrane potential). (**D**) Fluorescence emission spectra of different JF dyes overlaid with the absorbance spectrum of Ace2N. (**E**) Fluorescence change as a function of membrane voltage with different JF dye-Voltron conjugates. (**F**) Relative fluorescence of ASAP2f, Ace2N-mNeon, Voltron_525_ and Voltron_549_ in cultured neurons (n=70, 68, 48 and 62 measurements from five independent transfections for each construct). Illumination intensity ~10mW/mm^2^ at imaging plane. ***P<0.001, one-way analysis of variance (ANOVA) followed by Bonferroni’s test on each pair. Fluorescence was normalized to ASAP2f mean intensity. (**G**) Relative single molecule brightness of Ace2N-mNeon and Voltron_549_. ***P<0.001, two-tailed Student’s *t*-test. (**H**) Bleaching curves for ASAP2f, Ace2N-mNeon, Voltron_525_, and Voltron_549_ in primary neuron culture. Illumination intensity ~23 mW/mm^2^ at imaging plane. Bleaching curves were normalized to mean cellular fluorescence from **F** or normalized to the zero-time value (inset). (**I**) Mean time to bleach of Ace2N-mNeon and Voltron_549_ during single-molecule imaging, 100 ms frames. ***P<0.001, two-tailed Student’s *t*-test. (**J,K**) Simultaneous *in vivo* Voltron imaging and electrophysiology in larval zebrafish (extracellular) and adult *Drosophila* (whole cell), respectively.

We tested Voltron in neuron cultures using high-speed imaging with simultaneous wholecell patch clamp electrophysiology (**Fig. 1C**), first investigating different dye-Voltron combinations. Voltron could detect neuronal action potentials and sub-threshold potential changes using a range of JF dye ligands with emission maxima between 520 nm and 660 nm (**Fig. 1C-E** **and S6**). Voltron bound to JF_525_ (*i.e.*, Voltron_525_) exhibited the highest sensitivity, giving a fluorescence change of −23 ± 1% for a voltage step from −70 mV to +30 mV (Fig. 1E **and** Fig. S9); Voltron_549_ showed similar sensitivity. Voltron_525_ responded to voltage steps with sub-millisecond on and off time constants (Table S1 **and** Fig. S10). We compared the brightness and photostability of Voltron in neuronal cultures with two other recently described fluorescent protein-based GEVIs: Ace2N-mNeon (*11*) and ASAP-2f (*13*). Both Voltron_525_ and Voltron_549_ were brighter than Ace2N-mNeon (3-4×) and ASAP-2f (16-18ϗ) (**Fig. 1F**) in cell culture. This difference was not due to differences in expression levels; we compared the brightness of Voltron_549_ and Ace2N-mNeon at the single-molecule level, observing a similar 3-4× brightness difference (**Fig. 1G**). Voltron_525_ and Voltron_549_ were also more photostable in ensemble measurements (Fig. 1H, Tables S2,**S3** and Figs. S11,**S12**) as well as in single-molecule assays, where photobleaching times were 8-fold longer for Voltron_549_ than Ace2N-mNeon (Fig. 1I). Overall, the improved brightness and photostability of Voltron increase the photon yield by at least 10-fold relative to existing GEVIs in neurons.

We next deployed the chemigenetic Voltron indicator *in vivo*, observing that the protein could be reliably expressed and labeled with dye in mice, larval zebrafish, and adult fruit flies (Figs. 1**-4**, Fig. S13**-S34**). Simultaneous *in vivo* electrophysiology and imaging in both zebrafish and flies confirmed the detection of individual action potentials in singletrial imaging (Fig. 1J,K **and** Figs. S14**-S15**). For imaging in the mouse brain, we used a variant of Voltron with a soma-targeting sequence from Kv2.1 (*22,23*) (Voltron-ST, **Fig. S16**). The rapid kinetics of Voltron_525_-ST allowed clear observation of action potentials in fast-spiking parvalbumin positive interneurons in the CA1 region of mouse hippocampus (Fig. 2A-G **and** Fig. S17). We measured orientation tuning based on both spiking and subthreshold voltage signals in layer 2/3 pyramidal neurons in mouse primary visual cortex in response to drifting grating stimuli in the contralateral visual field, a benchmark for new indicators (*1,11*) (Fig. 2H-L and Fig. S18**-19**), and confirmed that spiking activity shows sharper orientation selectivity than subthreshold voltage signals (*24*). We extended the imaging period over several consecutive weeks by injection of additional JF_525_ HaloTag ligand prior to each imaging session (**Fig. 2J-L**).

**Fig. 2.**
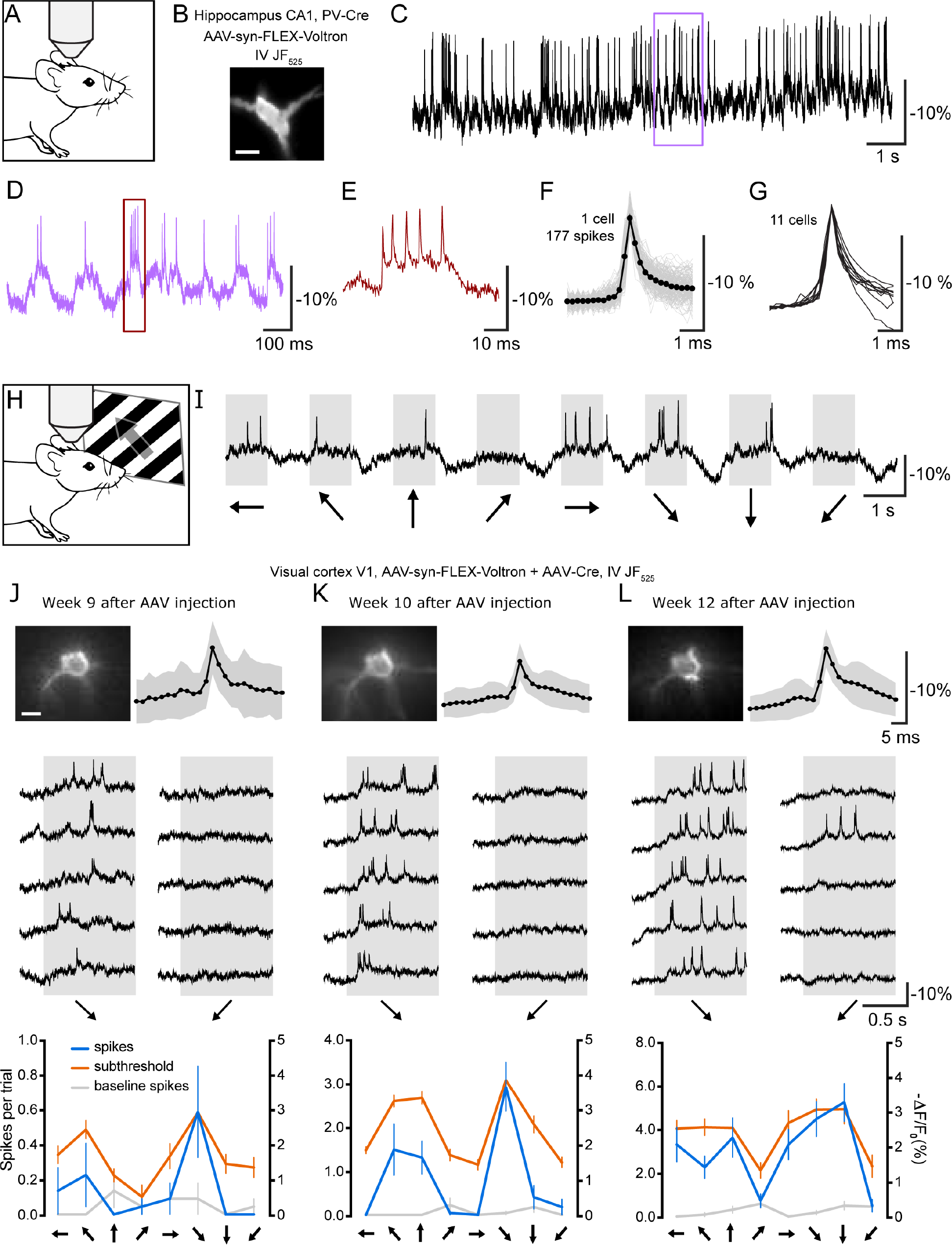
Membrane voltage dynamics in hippocampal parvalbumin (PV) neurons (A-G) and visual cortex pyramidal neurons (H-L) of mice using Voltron. (**C-E**) Voltron_525_ raw ΔF/F_0_ traces showing spontaneous spikes of a PV neuron (**B**) located at a depth of 60 μm in hippocampal CA1 region imaged at 3858 frames per second. Boxes indicate intervals shown at expanded time scales. Scalebar: 20 μm. (**F**) Overlay of 177 spikes detected during a 15 s period (gray) and their average (black). (**G**) Spike shape of 11 PV neurons. (**H**) Schematic of imaging of mouse primary visual cortex during display of drifting grating visual stimuli. (**I**) Example trace showing Voltron fluorescence during one trial of a sequence of visual stimuli. Arrows below represent the direction of movement of the drifting grating. (**J-L**) Top left, images of a pyramidal cell at a depth of 148 μm, imaged three times over a period of four weeks on the indicated weeks after virus injection. Scalebar: 10 μm. Top right, average of all spikes in session (black) and standard deviation (grey). Middle, raw ΔF/F_0_ trace for five repetitions in each session, showing two orthogonal orientations (indicated with arrows below) from the neuron pictured on the top left. Bottom, orientation tuning to full-frame drifting gratings of the neuron pictured on the top left, displayed from number of spikes during trials (blue), number of spikes during preceding inter-trial intervals (grey), and subthreshold ΔF/F_0_ (right y-axis) after low-pass filtering traces using a 10-point median filter. For each orientation, response is calculated by averaging the low-pass filtered trace between 100 - 400 ms after trial onset, and baseline is calculated by averaging the low pass filtered trace from 80 ms preceding trial onset to 20 ms after trial onset. Displayed as response minus baseline. Error bars represent standard error of the mean (s.e.m.) (20 - 22 repetitions per session).

To further assess the advantages garnered from Voltron’s improved photostability and brightness, we investigated the illumination intensity, imaging duration, and field of view used for *in vivo* imaging (**Fig. 3**). Using widefield microscopy and illumination intensities between 3 and 20 mW/mm^2^, we could clearly identify and distinguish action potentials from nearby neurons throughout 15 minutes of continuous imaging (SNR = 4.4 during final minute); (**Fig. 3B-E**). We expanded the field-of-view to include dozens of cortical interneurons labeled with Voltron_525_-ST via an NDNF-Cre mouse line (*25*), while imaging at 400 Hz (**Figs. 3F,G**, **Fig. S20**-**S33**). Even with this large field of view, we observed clear signals for spikes and subthreshold voltage signals in ~90% of neurons in focus within the imaging field. Overall, we imaged a total of 449 neurons (12 fields of view in 3 mice), demonstrating routine voltage imaging of populations of neurons in superficial mouse cortex (**Figs. 3G**, **Fig. S20**-**S33**). This unprecedented scale of *in vivo* voltage imaging enabled analysis of membrane potential correlations between many neuron pairs (**Fig. S21**).

**Fig. 3.**
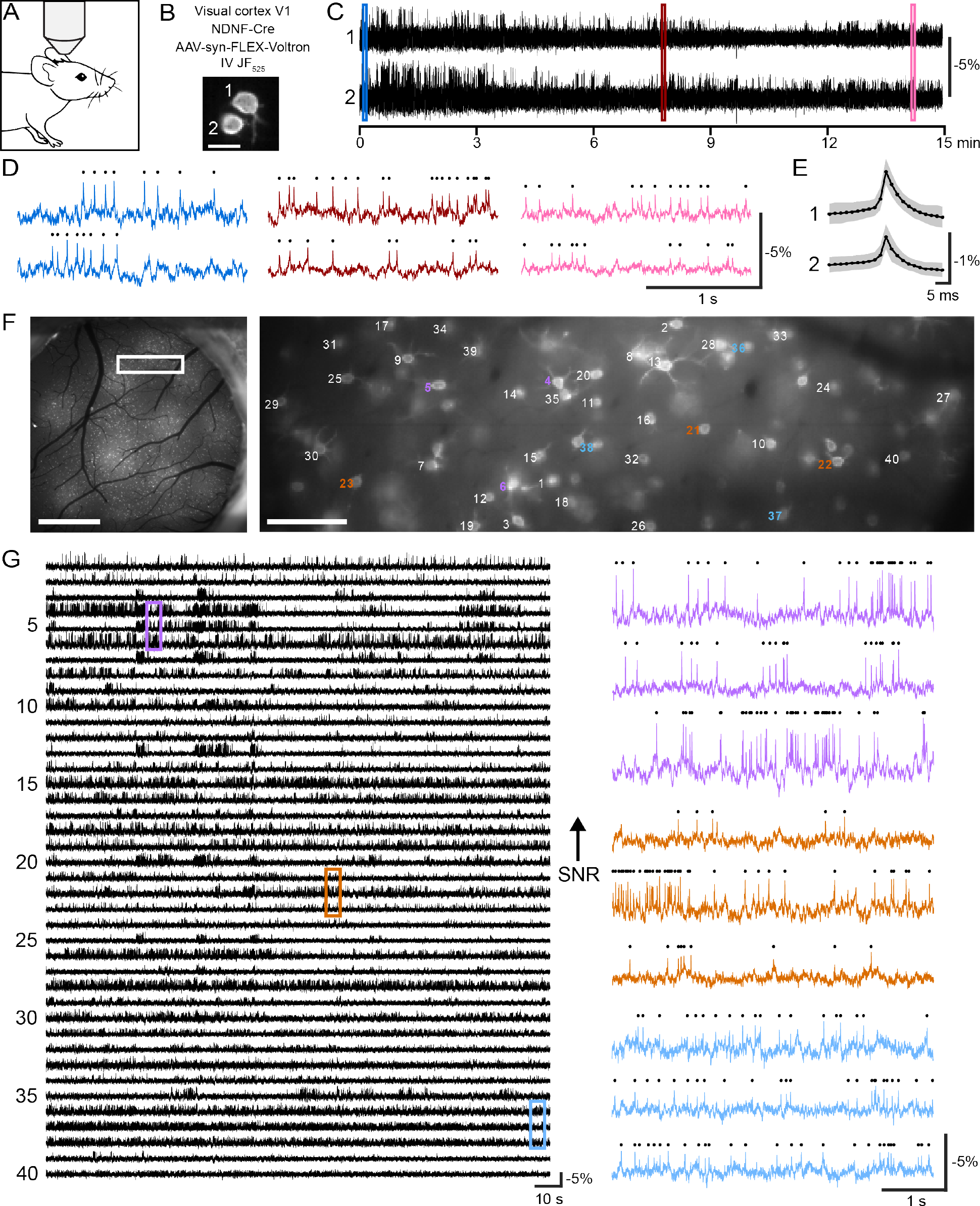
Long duration and large FOV imaging of voltage activity in GABA-ergic neurons in mouse neocortex. (**A**) Schematic of the imaging setup. (**B**) Image of two neurons expressing ST-Voltron_525_ in layer 1 of visual cortex of an NDNF-Cre mouse. Scalebar: 10 μm. (**C**) ΔF/F_0_ traces from neurons in **B**, recorded over 15 minutes. (**D**) Color-coded zooms of indicated regions of the traces in **C** with detected action potentials indicated by black dots above the fluorescence traces. (**E**) Average of all spikes in session (black) and standard deviation (grey). (**F**) Left panel: Fluorescence image of a cranial window over primary visual cortex (V1) in an NDNF-Cre mouse showing Cre-dependent expression of ST-Voltron_525_ (bright spots). Scalebar: 1 mm. Right panel: zoomed image of **F** in the area indicated by the white rectangle, with neurons labels corresponding to fluorescence traces in **G**. Scalebar: 100 μm. (**G**) Left panel: ΔF/F_0_ traces during 3 min. recording from neurons pictured in (**F**), in decreasing order of signal-to-noise ratio. Right panel: zooms of ΔF/F_0_ traces from color-coded regions of **G** with detected action potentials represented as black dots above, illustrating representative traces with high (top), medium (middle), and low (bottom) SNR. Traces have been background-subtracted, which removes shared subthreshold membrane potential fluctuations (Supplementary Methods; Compare vs. Fig. S20 without subtraction).

We then used Voltron to image behaving zebrafish larvae, which reliably respond to visual input with fast, directed swim bouts that are tailored to the details of the stimulus (*26*). We sought to uncover how this sensory-to-motor transformation unfolds in neuronal populations at fine timescales that are inaccessible with calcium imaging. We first verified that Voltron could detect action potentials and subthreshold voltage signals in live zebrafish using several different colors of dye ligands (Figs. S14 **and S34**). We then used Voltron_525_ to monitor neural spiking patterns during visual-motion-induced swims (**Fig. 4A**). We recorded activity patterns from 179 neurons across 43 fish in a motor-sensory nucleus in the tegmental area of the midbrain (**Figs. 4B**, **S35A**), yielding data on subthreshold membrane voltage modulation as well as automatically-detected spike times (**Figs. 4C**, **S35B-C**). We found neuron populations with different temporal activity patterns, including neurons whose firing rate increased ~1 second before the fish started swimming (**Fig. S35D-E**, ‘Ramp’), neurons whose firing rate was suppressed each time the fish swam (**Fig. 4D**, ‘Off’), and neurons that fired each time the fish swam (**Fig. 4D**, ‘Onset’ and ‘Late’). Of the latter types, some fired just before swimming (~20 ms before swim onset, ‘Onset’) and others fired just after swimming (~10 ms after swim onset, ‘Late’). There was a change in subthreshold voltage that preceded these firing-rate changes by tens of milliseconds (**Fig. 4D**). The neuron types were spatially intermingled within this midbrain nucleus (**Fig. 4E-F**). The existence of neurons that fired before swimming and neurons that fired after swimming suggests that this nucleus both partakes in the generation of swim bouts and receives an efference copy of motor output (**Fig. 4G**). Thus, Voltron allows for the dissection of population motor coding and sensorimotor integration circuits in ways that neither single-cell electrophysiology nor population calcium imaging can.

**Fig. 4.**
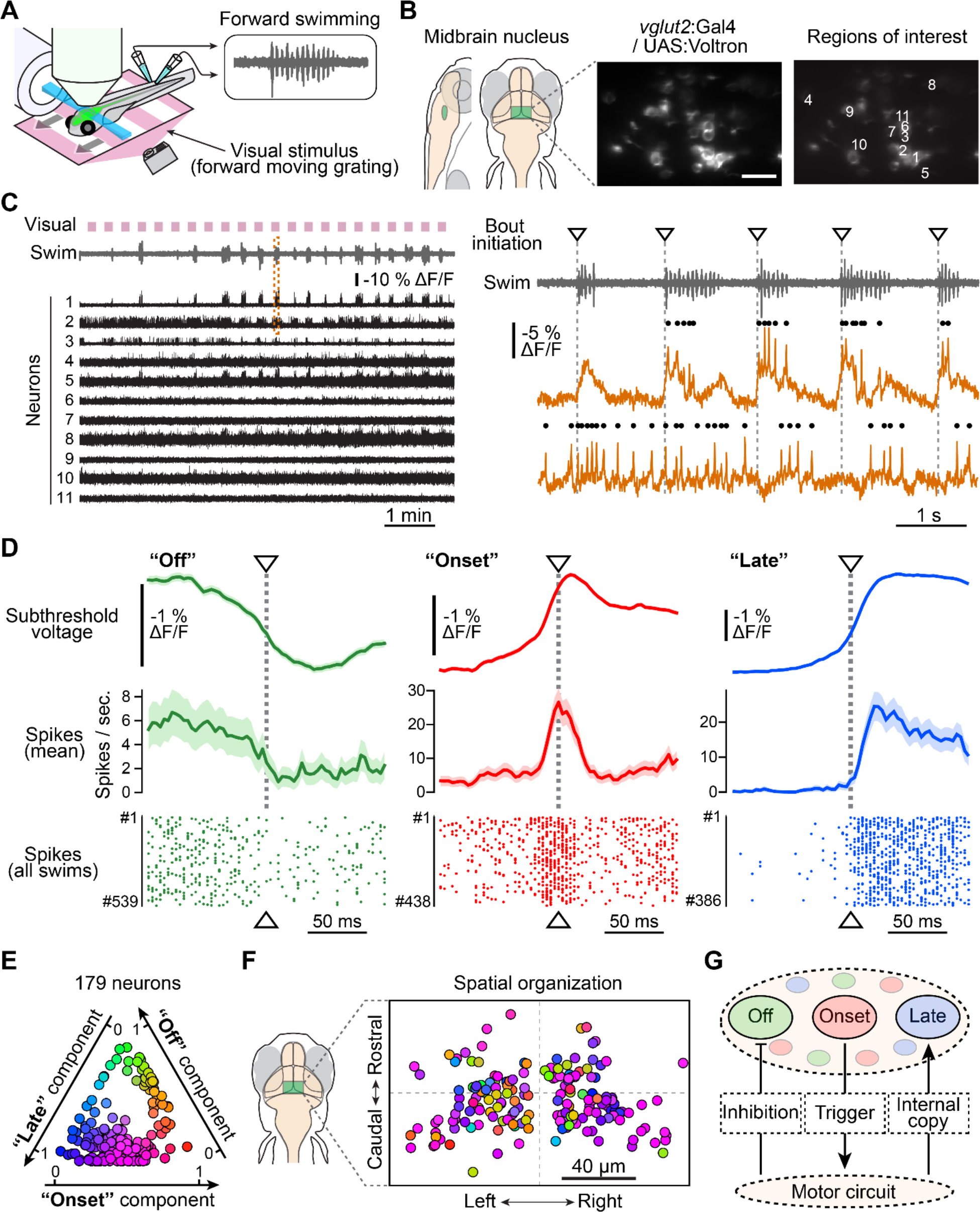
Voltron reveals millisecond-timescale neural dynamics during swimming behavior in zebrafish. (**A**) Schematic illustration of the setup. An immobilized zebrafish is placed under the light-sheet microscope and the motor signals (inset) from its tail are recorded during the imaging session using a pair of electrodes. Visual stimuli (forward drifting gratings) for triggering swimming responses are presented below the fish. (**B**) Left panel: anatomical location of the imaged brain region (midbrain nucleus; see Fig. S35A). Center, a representative field of view of the imaged region expressing Voltron. Scale bar, 20 μm. Right, the position of neurons analyzed in **C**. (**C**) Left panel: periods of visual motion (pink) and swim signals (grey) are plotted above Voltron fluorescence traces (black) simultaneously recorded from 11 neurons shown in **B**. Right panel: zoom of swimming signals (top) and Voltron fluorescence traces from two representative neurons (bottom) are expanded from the dashed box in the left panel. Dots on the top of each trace represent spikes recognized by the algorithm described in Fig. S35B-C. Downward triangles and dotted gray lines indicate initiation of each swim bout. (**D**) Mean subthreshold signal (top), mean spiking frequency (middle) and spike raster plots (bottom) near the initiation of swim bouts from three representative neurons: “Off” (green), “Onset” (red) and “Late” (blue) neuron. Shadows in the top and middle panels represent s.e.m. across swim events. (**E**) Classification of recorded neurons by their mean subthreshold signals near the initiation of swim bouts. 179 neurons recorded from 43 fish were classified using non-negative matrix factorization and colored according to the weights for three factors: “onset” (red), “off” (green) and “late” (blue). The details of this classification are described in the Methods. (**F**) Spatial organization of the same population of neurons as in **E**. Neurons from multiple fish are superimposed to a single map based on the distance from the center of this midbrain nucleus. (**G**) Hypothetical model of neural activity modulation in this midbrain nucleus. “Onset” neurons send motor commands to downstream motor circuits to trigger swim bouts, while activity of “off” neurons is inhibited. “Late” neurons receive internal copy signals of ongoing swim bouts from the motor circuit.

Finally, we tested Voltron in adult *Drosophila in vivo* by expressing the protein in a pair of dopaminergic neurons, one in each brain hemisphere, which innervate a single compartment in the mushroom body. We detected strong spiking signals from axons and dendrites of these neurons using Voltron_549_ (**Figs. 1K**, **S15**), which matched spikes detected using electrophysiology. In some neuronal cell types in *Drosophila*, calcium indicators located in the cell body have failed to exhibit fluorescence changes even under conditions where high spike rates are expected (*27*). However, spikes were clearly detectable when imaging from the soma of dopamine neurons with Voltron (**Fig. S15E**). Remarkably, we could clearly distinguish spikes from the two neurons based on the amplitude of the spiking signals even when imaging from neuropil where their axons overlap extensively, likely because each bilaterally-projecting cell contributes a denser innervation of the mushroom body in the ipsilateral hemisphere (**Fig. S15D**).

Combining the molecular specificity of genetically encoded reagents with the superior photophysics of chemical dyes is an established path to improved imaging reagents (*14*). However, previous attempts to create hybrid small-molecule:protein indicators using a variety of approaches have not been successful for *in vivo* imaging (*28*). Here, we engineered a modular sensor scaffold where the targeting and sensor domains are genetically encoded and only the fluorophore and its protein-binding anchor are synthetic. The resulting chemigenetic indicator, Voltron, exhibits substantially increased photon output, enabling *in vivo* voltage imaging of many more neurons over longer times—approximately 10^2^ more neuron-minutes than other sensors. This improvement enables imaging experiments that reveal how the precise electrical dynamics of neuronal populations orchestrate behavior over different time scales.

## Acknowledgements

We acknowledge the Janelia Vivarium, Cell Culture, Instrument Design & Fabrication, Imaging, Molecular Biology, and, Virus Production facilities for assistance. Specifically, we would like to thank Brenda Shields, Deepika Walpita, Jim Cox, Chelsea McGlynn, Damien Alcor, Aaron Taylor, Jared Rouchard, Kim Ritola, Xiaorong Zhang and Jordan Towne. We thank Ziqiang Wei for discussions on data analysis. Funding was provided by Simons Collaboration on the Global Brain Research Awards 325171 (MA, LP) and 542943SPI (MA, LP), IARPA MICRONS D16PC00003 (LP), NIH R01EB22913 (LP), Taiwan Ministry of Science and Technology MOST106-2628-B-010-004, MOST105-2628-B-010-005, MOST106-2320-B-010-012 and Taiwan National Health Research Institute NHRI-ex-107-10509NC (TWC, BJL), as well as the Howard Hughes Medical Institute.

## Supplementary Materials

### Materials and Methods

#### Reagent availability

Voltron plasmids pCAG-Voltron (plasmid#), pCAG-Voltron-ST (plasmid#), pAAV-hsyn-Voltron (plasmid#), pAAV-hsyn-flex-Voltron (plasmid#), pAAV-hsyn-flex-Voltron-ST (plasmid#), pTol2-Huc-Voltron (plasmid#), pTol2-Huc-Voltron-ST (plasmid#), p10XUAS-IVS-Syn21-Voltron-p10 (plasmid#), and p13XLexAOP2-IVS-Syn21-Voltron-p10 (plasmid#) have been deposited at Addgene (http://www.addgene.org).

AAV-hsyn-flex-Voltron-ST virus is available from Addgene (http://www.addgene.org).

Transgenic *Drosophila* stocks for UAS-Voltron and LexAop-Voltron in multiple landing sites are available from the Bloomington Drosophila Stock Center (http://flystocks.bio.indiana.edu).

##### UAS

Voltron transgenic zebrafish are available from the Ahrens Lab at Janelia Research Campus, and from ZIRC (http://sudheer.zinovyevcurie.com).

#### Cloning

Generally, cloning was done by restriction enzyme digest or PCR amplification of plasmid backbones, PCR amplification of inserted genes, and isothermal assembly to combine them, followed by Sanger sequencing to verify DNA sequences. The genes for QuasAr1 and QuasAr2 (*1*) were amplified from Addgene plasmids 51629 and 51692. The gene for Ace2N was synthesized (Integrated DNA Technologies) with mammalian codon optimization (*2*). The soma localization tag was synthesized (Integrated DNA technologies) through adding a 66 amino acid domain from the Kv2.1 potassium channel (residues 536 to 600) (*3*). This domain directs localization to clusters at the soma and proximal dendrites (*4*). Linker length variants were generated by Quikchange site-directed mutagenesis (Agilent). For expression in primary neuron cultures, sensors were cloned into a pcDNA3.1-CAG plasmid (Invitrogen) at the NheI and HindIII sites. For expression in zebrafish, Voltron and Voltron-ST were cloned into the pTol2-HuC vector (for panneuronal expression) at the AgeI restriction sites and into the pT2-Tbait-UAS vector (for Gal4-dependent expression) at the EcoRI and PspXI restriction sites. For expression in *Drosophila melanogaster*, Voltron was cloned into p10XUAS-IVS-Syn21-p10 at the XhoI and XbaI sites. For Cre-dependent expression in mouse brain, Voltron and Voltron-ST were cloned into a pAAV-hsyn-flex plasmid at the BamHI restriction sites. The DNA and amino acid sequences of Voltron and Voltron-ST are given in **Fig. S2**. Plasmids and maps are available from Addgene.

#### *In vitro* spectroscopy of fluorophores

To create JFdye-HaloTag conjugates, 5 μM JFdye HaloTag ligand and 10 μM HaloTag protein were incubated in 10 mM HEPES with 0.1 mg/ml CHAPS at pH 7.3 at 4°C overnight. Completeness of dye-binding was determined by titrating HaloTag protein (2.5 μM to 12.5 μM) with fluorogenic JF_635_ HaloTag ligand (5 μM) in overnight reactions and then measuring absorbance at 640 nm. Additionally, thin-layer chromatography was performed on a reaction of 5 μM JF_549_ with 7.5 μM HaloTag, which showed >95% of the dye was bound to HaloTag. Fluorescent proteins sfGFP (parent fluorophore of ASAP2f) and mNeonGreen (parent fluorophore of Ace2N-mNeon) were purified from *E. coli*. All photophysical measurements used either 1 μM solutions of JFdye-HaloTag conjugate in 10 mM HEPES buffer at pH 7.3, or 1 - 3 μM purified fluorescent proteins in 100 mM MOPS buffer at pH 7.2. Absorbance measurements were performed on a UV-VIS spectrometer (Lambda 35, Perkin Elmer). Fluorescence excitation and emission spectra were measured using a fluorimeter (LS55, Perkin Elmer). Quantum yield measurements were performed using an integrating-sphere spectrometer (Quantaurus, Hamamatsu). Extinction coefficients for the JFdye-HaloTag conjugates were determined from peak absorbance at known concentration of JF dye. Extinction coefficients for fluorescent proteins were determined by the alkali denaturation method, using the extinction coefficient of denatured FP equal to that of denatured GFP (ε = 44000 at 447 nm) (*5*).

#### Fluorescence microscopy for photobleaching

To investigate photobleaching of fluorophores in solution, aqueous droplets of JFdye-HaloTag conjugates or fluorescent proteins were made by aliquoting 5 μl of a fluorophore solution into 45 μl of 1-Octanol and agitating by tapping or brief vortexing. 5 μl of the emulsion mixture was sandwiched between a pre-silanized glass slide and a glass coverslip to disperse isolated microdroplets of dye-conjugates or proteins for fluorescence microscopy. To perform fluorescence microscopy, microdroplets were continuously illuminated using an inverted microscope (Eclipse Ti2, Nikon) with a 40x (N.A. = 1.3, Nikon) oil immersion objective (PLAN Flour, Nikon). Fluorescence excitation was achieved using an LED (SpectraX Light engine, Lumencor) with the following filter sets for the respective fluorophores: For sfGFP, mNeonGreen and JF_505_ (FITC5050A cube (semrock): FF02-475/50, FF506-Dio3, FF01-540/50); for JF_525_ (510/25 excitation filter, T525lprx dichroic(Chroma), 545/40 emission filter); for JF_549_ (Cy34040C cube (semrock): FF01-531/40, FF562-Dio3, FF01-593/40); for JF_585_ ( 49912 cube (Chroma): ZET594/10x, ZT594rdc, ET610lp) and for JF_635_ ( 89000 cube (Chroma), ET645/30x, 89100bs, ET705/72m). Power at the imaging plane for each filter set was set to 12 mW determined with a microscope slide power sensor (S170C, Thorlabs). From measurement of the sample area illuminated, the irradiance was determined to be 40 mW/mm^2^. In order to calculate the excitation rate *W* (photons absorbed/sec), the LED excitation spectrum was measured after the objective for each filter set using a fiber spectrometer (QE65000, Ocean Optics). Fluorescence images were collected using a scientific CMOS camera (ORCA-Flash 4.0, Hamamatsu) and image acquisition was performed using HCImage Live (Hamamtsu). Each sample was bleached continuously for 10 min. and images were acquired at 1Hz. Fluorescence intensity from each droplet was obtained after background subtraction using ImageJ software.

To investigate photobleaching of GEVIs in cells, Voltron, Ace2N-mNeon, and ASAP2f were transfected in hippocampal neurons extracted from P0 to P1 Sprague-Dawley rat pups. After transfection, hippocampal neurons were plated onto 35 mm glass-bottom dish (MatTek) coated with poly-D-lysine (Sigma) and cultured for 8-10 days in NbActiv4 medium (BrainBits). For labeling Voltron-expressing neurons, cells were incubated with a 100 nM JFdye-HaloTag ligand for 30 minutes. The same setup and procedure used with droplets above was used to measure photobleaching of GEVIs in cells.

#### Photobleaching analysis

The bleaching profile of individual cells or droplets was fit to either a single or double exponential function of the form 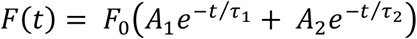 to obtain time constants τ_1_, τ_2_ and weighting *A*_1_, *A*_2_. Data fitting was performed in MATLAB (MathWorks) and Origin (OriginLab), and goodness of fit assessed by minimal residual sum of errors or minimal *χ*^2^. To quantify photobleaching across fluorophores requires knowledge of the excitation rate ***W*** and the fluorescence quantum yield *ϕ*_*f*_. The excitation rate ***W*** was computed (*6*) from integration over the wavelength dependence of the product of measured extinction coefficient and irradiance spectral profile. The fluorescence quantum yield for the GEVIs is not directly measured, and assumed to be the same as that measured for the parent fluorophores. Three quantities characterizing bleaching were calculated for each fluorophore or GEVI. These are (i) the calculated time *t*_1/2_ for the fluorescence rate to drop to 1/2 its initial value, scaled by the excitation rate to achieve an initial fluorescence rate of 10^3^ photons/sec (*6*), (ii) the total number of photons emitted before photobleaching *N*_*p*_ (the photon budget), and (iii) the photobleaching probability *P*_b_. These are per-molecule quantities averaged over the ensemble of molecules in each droplet or cell. The characteristic time t_1/2_ was found by determining from the raw data, the time *t*_*raw*_ for 50% reduction in fluorescence *F*(*t*_*raw*_)/*F*_0_ = 0.5, from which

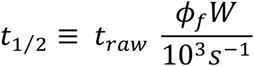

where *ϕ*_*f*_ is the fluorescence quantum yield and *W* is the excitation rate. To determine *N*_*p*_, the fit function *F*(*t*) was integrated over time, where initially *F*_0_ ≡ *ϕ*_*f*_*W*,

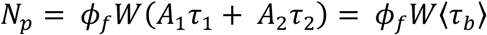

where 〈τ_*b*_〉 is the amplitude-weighted lifetime (*7,8*) 〈τ_*b*_〉 = *A*_1_*τ*_1_ + *A*_2_τ_2_. The photobleaching probability *P*_*b*_, based on rate equation models where bleaching proceeds from singlet or triplet states, is inversely related to the total number of fluorescent photons emitted, *N*_*p*_ = *ϕ*_*f*_/*P*_*b*_ (*9,10*), or

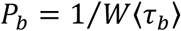

Of the three photobleaching quantities above, the photobleaching probability is most rigorous as it is independent of the fluorescence quantum yield.

#### Single-Molecule Imaging and Analysis

Hippocampal neurons extracted from P0 to 1 Sprague-Dawley rat pups were transfected with Ace2N-mNeon and Voltron plasmids by electroporation (Lonza, P3 Primary Cell 4D-Nucleofector X kit) according to the manufacturer’s instruction. After transfection, hippocampal neurons were plated onto 25 mm ultra-clean cover glasses coated with poly-D-lysine (Sigma) and cultured for 9 days in NbActiv4 medium (BrainBits). To label Voltron-expressing neurons, cultures were incubated with 2 nM JF_549_ HaloTag ligand for 15 mins, then transferred to the Attofluor cell chamber (Thermo Fisher Scientific) and supplemented with Tyrode’s solution (140 mM NaCl, 5 mM KCl, 3 mM CaCl_2_, 1 mM MgCl_2_, 10 mM HEPES, 10 mM glucose, pH 7.35). Single-molecule imaging was performed on a Nikon Eclipse TiE Motorized Inverted microscope equipped with a 100X oil-immersion objective lens (N.A. = 1.49, Nikon), 488/561 nm laser lines, an automatic TIRF illuminator, a perfect focusing system and a Tokai Hit environmental control (humidity, 37 °C, 5% CO_2_). Excitation light was passed through a 405/488/561/647 nm laser quad band filter set filter that allows 488 nm or 561 nm light through to the sample (Chroma set number 89902). Emission from sample was collected through the same filter set then passed through a splitter (dichroic mirror: T560lpxr (Chroma), with perpendicular emission filters: ET525/50m (Chroma) and ET605/52m (Chroma)) to split green and red fluorescence. The light was then collected onto two EMCCD cameras (iXon Ultra 897, Andor). Samples were first pre-bleached to achieve sparse single molecule detections. We calibrated the laser power output for 488 nm (to image Ace2N-mNeon) and 561 nm (to image Voltron_549_) to 17.5 mW with a Si Sensor power meter (Thorlabs, PM202). Images were acquired under TIRF imaging mode with 10 Hz frame rate. 1000 frames were recorded for each imaging area and 10 imaging areas were collected for each indicator. For image analysis, single molecules lasting for at least 3 frames were manually selected. The brightness and mean time to photobleach of these molecules were determined with ImageJ (1.51n) quantification tools to assess the single-molecule photostability of Ace2N-mNeon and Voltron_549_.

#### Fluorescence imaging in primary neuron culture

All culture imaging was performed in imaging buffer containing the following (in mM): 145 NaCl, 2.5 KCl, 10 glucose, 10 HEPES, pH 7.4, 2 CaCl_2_, 1 MgCl_2_. Wide-field imaging was performed on an inverted Nikon Eclipse Ti2 microscope equipped with a SPECTRA X light engine (Lumencore), 40X oil objective (NA = 1.3, Nikon), and imaged onto a scientific CMOS camera (Hamamatsu ORCA-Flash 4.0). A FITC filter set (475/50 nm (excitation), 540/50 nm (emission), and a 506LP dichroic mirror (FITC-5050A-000; Semrock)) was used to image mNeonGreen, ASAP1, and ASAP2f. A Cy3 filter set (531/40 nm (excitation), 593/40 nm (emission), and a 562LP dichroic mirror (Cy3-4040C-000; Semrock)) was used to image Volton_549_. A custom filter set (510/25 nm (excitation), 545/40 nm (emission), and a 525LP dichroic mirror (Semrock)) was used to image Voltron_525_. A quad bandpass filter (set number: 89000, Chroma) was used along with the appropriate color band from the SPECTRA X light source to image Voltron_505_, Voltron_585_ and Voltron_635_. For time-lapse imaging during field stimulation or simultaneous electrophysiology measurements, neurons were imaged at 200 - 3200 Hz depending on the experiment. The LED light power output at the imaging plane was measured with a Si Sensor power meter (Thorlabs, PM202) for each imaging experiment.

For quantifying brightness of voltage indicators expressed in neurons, the excitation spectrum was measured after the objective for each excitation filter used using a spectrometer (QE65000, Ocean Optics). The spectrum was then integrated to get the excitation rate *W* as described above (see section: Photobleaching analysis). As with the photobleaching experiments, when the data sets for the light spectrum and extinction coefficient are taken at incommensurate wavelengths, interpolation was used to re-cast the wavelengths of one of the data sets using MATLAB (MathWorks). The fraction of collected fluorescence using the emission filter compared to the total emission spectrum of the fluorophore was calculated. Illumination intensity of 20 mW/mm^2^ at imaging plane was used for all indicators. Fluorescence images were acquired from five independent transfections for each construct for brightness measurements. Using MATLAB (MathWorks), fluorescence intensity was then corrected using the calculated excitation rates (*W*), fraction of emission collected, and quantum efficiency of the Hamamatsu ORCA-Flash 4.0 camera over the emitted wavelengths for ASAP2f, Ace2N-mNeon, Voltron_525_ and Voltron_549_. Values were calculated relative to ASAP2f.

#### Simultaneous field stimulation and fluorescence imaging in primary neuron cultures

A stimulus isolator (A385, World Precision Instruments) with platinum wires was used to deliver field stimuli (50V, 83 Hz, 1 ms) to elicit action potentials in cultured neurons as described previously (*11*). The stimulation was controlled using an Arduino board and timing was synchronized with fluorescence acquisition using the Nikon Elements software and a national instruments PXI-6723 board.

#### Simultaneous electrophysiology and fluorescence imaging in primary neuron culture

All imaging and electrophysiology measurements were performed in imaging buffer (see “Fluorescence imaging in primary neuron culture” section) adjusted to 310 mOsm with sucrose. For voltage clamp measurements, 500 nM TTX was added to the imaging buffer to block sodium channels. Synaptic blockers (10 μM CNQX, 10 μM CPP, 10 μM GABAZINE, and 1 mM MCPG) were added to block ionotropic glutamate, GABA, and metabotropic glutamate receptors (*11*).

Filamented glass micropipettes (Sutter Instruments) were pulled to a tip resistance of 4 - 6 MW. Internal solution for current clamp recordings contained the following (in mM): 130 potassium methanesulfonate,10 HEPES, 5 NaCl, 1 MgCl_2_, 1 Mg-ATP, 0.4 Na-GTP, 14 Tris-phosphocreatine, adjusted to pH 7.3 with KOH, and adjusted to 300 mOsm with sucrose. Internal solution for voltage clamp recordings contained the following (in mM): 115 cesium methanesulfonate,10 HEPES, 5 NaF, 10 EGTA, 15 CsCl, 3.5 Mg-ATP, 3 QX-314, adjusted to pH 7.3 with CsOH, and adjusted to 300 mOsm with sucrose.

Pipettes were positioned with a MPC200 manipulator (Sutter Instruments). Whole cell voltage clamp and current clamp recordings were acquired using an EPC800 amplifier (HEKA), filtered at 10 kHz with the internal Bessel filter, and digitized using a National Instruments PCIe-6353 acquisition board at 20 kHz. Data were acquired from cells with access resistance < 25 MΩ. WaveSurfer software was used to generate the various analog and digital waveforms to control the amplifier, camera, light source, and record voltage and current traces. For fluorescence voltage curves, cells were held at a potential of −70 mV at the start of each step and then 1 second voltage steps were applied to step the potential from −110 mV to +50 mV in 20 mV increments. For current-clamp recordings to generate action potentials, current was injected (20 - 200 pA for 1-2 s) and voltage was monitored.

#### Imaging Parvalbumin (PV) neurons in mouse hippocampus

Hippocampal PV neuron imaging was performed using adult PV-Cre mice (JAX 008069). Imaging window was implanted using procedures similar to those described in Dombeck et. al. (*12*). In short, a circular craniotomy (3 mm diameter) was made centered at 2.0 mm caudal and 2.0 mm lateral to bregma. The surface of CA1 was exposed by gently removing the overlying cortex with aspiration. AAV2/1-syn-Flex-Voltron-ST virus was diluted to 1.9×10^12^ GC/ml and injected at three locations (separated by 800 μM, 30nl per location) 200 μM from CA1 surface. The imaging window (constructed by gluing a 3 mm diameter cover glass to a stainless steel cannula of 3 mm diameter and 1.5 mm height) was placed onto the hippocampus and glued to the skull using super-bond C&B (Sun Medical). A titanium head bar was glued to the skull for head fixation during imaging.

Imaging experiments started 4-5 weeks after surgery. JF_525_-HaloTag ligand (100μl, 1mM) was delivered using retro-orbital injection (*13*) 1 day before imaging. Labeled PV neurons (25 - 195 μM deep) were illuminated using a green LED (M530L3, Thorlabs) through an excitation filter (FF02-520-28, Semrock). A field aperture (diameter ~1 mm) was used to limit illumination to a circular area (~160 μM diameter at sample) around the cell of interest. The excitation intensity was ~25 mW/mm^2^ at the sample plane. JF_525_ fluorescence was collected using a 16X 0.8 NA objective (Nikon), separated from excitation light using a dichroic mirror (540lpxr, Chroma) and an emission filter (FF01-575-59, Semrock), and imaged onto a sCMOS camera (Zyla 4.2 plus, Andor). Images were collected at 3858 Hz.

Image analysis was performed in MATLAB. Brain movement was corrected using ImageJ plugin TurboReg (14). A constant camera offset (measured by taking images without illumination) was subtracted from each frame. The fluorescence of each cell was measured by averaging pixels within a region of interest covering the cell body. To detect action potentials (AP), slow baseline fluctuation (measured by moving average with 20 ms window) was first subtracted from the raw fluorescence trace. The timings of AP events were detected as local minima of the baseline subtracted trace with amplitudes larger than four times the standard deviation and peaks separated by at least 5 ms from each other. To quantify AP waveform, 5 ms segments of fluorescence signal around the detected peaks were taken from the raw fluorescence trace, peak aligned and then averaged. The AP amplitude was measured as percent change (F-F_0_)/F_0_ with F_0_ being the fluorescence baseline averaged over a time window 2.5 ms to 1.5 ms before the peak of an individual AP. The rise time, decay time, and the width of the AP waveform was measured using the averaged trace for each cell. The rise time was the time from half the amplitude to the peak. The decay time was the time from the peak to half the amplitude in the decay phase. The width (full width at half maxima, FWHM) was the sum of rise and decay time. This is shown in **Fig. S17**.

#### Imaging mouse cortex

NDNF-Cre mice (JAX 28536) were used for imaging Layer 1 neurons (2 females, 1 male; 100 -120 days old at the time of the window surgery). C57BI/6NCrl (Charles River Laboratories) mice were used for imaging Layer 2/3 neurons (2 females; 100-120 days old). NDNF-Cre mice were injected with 30 nl of AAV2/1-syn-FLEX-Voltron-ST (titer, 2*10^12^ GC/ml) at 8-12 injection sites 200 μM deep (injection rate, 1 nl/s). C57BI/6NCrl mice were injected with 30 nl mixture of AAV2/1-syn-FLEX-Voltron-ST (titer, 2*10^12^ GC/ml) and AAV9-CamKIIa-Cre (titer, 10^8^ GC/ml) 250 μM deep. AAV2/1-syn-FLEX-Voltron without soma targeting signal was injected in additional NDNF-Cre mice (titer, 2*1012 GC/ml)) and C57BI/6NCrl mice (AAV2/1-syn-FLEX-Voltron (titer, 2*10^12^ gc/ml) + AAV9-CamKIIa-Cre (titer, 10^8^ gc/ml)). This resulted in diffuse fluorescence and was not used for imaging experiments shown in this manuscript.

Cranial windows (4 mm diameter) were implanted over the injection sites in visual cortex (centered on − 2.5 mm lateral, + 0.5 mm anterior from lambda). Four to nine weeks later JF_525_ dye was injected into the retro-orbital sinus. Imaging was done 2 to 6 days after dye injection, with subsequent dye injections and imaging 1 to 6 weeks after the first imaging session. To prepare the JF dye for injection, 100 nanomoles of lyophilized JF_525_ were dissolved in 20 μl of DMSO, 20 μl Pluronic F-127 (20% w/v in DMSO), and 60-80 μl of PBS (final dye concentration 1 μM). Mice were anesthetized with 2-3% isoflurane and 100 μl of the dye solution was injected into the retro-orbital sinus of the right eye using a 27 - 30 gauge needle (*13*).

For imaging experiments of Layer 1 neurons, a wide-field fluorescence microscope equipped with a water immersion objective (20X, NA 1.0, Olympus XLUMPLFLN) was used for imaging. Illumination was delivered using a 525 nm LED (Mightex, LCS-0525-60-22); intensity at the sample, <20 mW/mm^2^. An mKO/mOrange filter set (530/30 nm (excitation), 575/40 nm (emission), and a 550LP dichroic mirror (Chroma, 49014)) was used for fluorescence imaging of Voltron_525_. Images were collected using a sCMOS camera (Hamamatsu Orca Flash 4.0 v3) at frame rates of 400-1000 Hz. A 0.55X magnification camera tube was placed between the objective and the camera for imaging large fields of view of 1064 μM X 266 μM (**Fig. 3F-G** and **Fig. S20**-**S33**). The pixel resolution was 2.08 μM/pixel. For smaller fields of view (**Fig. 3B-E** and **Fig. S18**) we used a 1X camera tube. The pixel size was 1.04 μM. Mice were awake and imaged in darkness.

To image Layer 2/3 pyramidal cells, the following changes were made from the imaging protocol for Layer 1 interneurons: Images were recorded at frame rate of 500-700 Hz. Illumination intensity at the sample was < 50 mW/mm^2^. 1X camera tube was used and the field of view imaged was typically 50 μM X 50 μM. The pixel size was 1.04 μM. A digital mirror device (Texas Instruments, LightCrafter) restricted the illumination to the cell being imaged. Mice were imaged while lightly anesthetized and passively viewing drifting gratings (described below).

#### Visual stimulation for pyramidal cell recordings

Mice were presented with drifting grating visual stimuli during imaging sessions (spatial frequency: 0.03 cycles/degree, temporal frequency: 1Hz, trial period: 1s, and inter-trial interval: 1s). Gratings were shown in blue with a black background. During the inter-trial interval, the screen was black. Eight orientations separated by 45° were presented. Mice were anesthetized during all sessions. To induce anesthesia, we injected chlorprothixene (0.2 mg/ml, 5ul/g weight mouse) into the hind paw followed by keeping the mouse in a chamber with 2-3% isoflurane for 1-2 minutes. Anesthesia was maintained at 0.4-0.8% isoflurane for the duration of the imaging session. Mice were kept on a heating blanket at a temperature of 37°.

#### Data Analysis

##### Analysis of Layer 2/3 pyramidal cell imaging

Motion was removed using a rigid registration algorithm. A constant camera offset was subtracted from each frame. A region of interest (ROI) was manually drawn around the neuron. The initial trace (X0) is the mean intensity over the ROI in time. X0 was fit with a piecewise linear curve using a Savitzky-Golay filter with a window size of 10 s to estimate the slow baseline fluctuations, F0. We calculated ΔF/F as 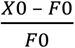. Spike times were manually selected as large amplitude local minima in the ΔF/F trace occurring in periods of depolarization and separated from other local minima by at least 2 ms.

Visual responses (**Fig. 2H-L** and **Fig. S18**) were calculated as the average number of spikes during the trial for each orientation, averaged over repetitions. To estimate the subthreshold fluctuations, we low-pass filtered the ΔF/F trace at 50 Hz using a median filter. The response for each orientation was calculated as the average of the low-pass filtered trace from 100 ms to 400 ms after the trial start. The baseline was calculated as the average of the low-pass filtered trace from 80 ms before trial start to 20 ms after the trial start. We subtracted the baseline from the response for each trial and averaged over 20 repetitions.

The orientation selectivity index (**Fig. S19**) was calculated as:

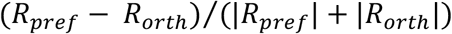

where *R*_*pref*_ is the response (mean spikes in trial or mean subthreshold membrane potential) to the preferred orientation, and *R*_orth_ is the response to the orientation 90° away from the preferred orientation.

##### Analysis of Layer 1 interneuron imaging

To identify neuronal activity and spatial structure from Voltron recordings, we designed an iterative spatial and temporal filtering approach we call: Spike Pursuit. In essence, Spike Pursuit begins with a poorly estimated voltage trace for a neuron, and uses detected spikes to iteratively estimate improved temporal and spatial filters that increase the signal to noise ratio of the spikes while controlling for overfitting. Spike pursuit relies on linear methods (the whitened matched filter for temporal filtering, and regularized linear regression for spatial filtering) (*15*).

Motion was removed using Fast Fourier transform-based rigid registration in MATLAB. Initial ROIs were manually drawn around each neuron in the field of view. Data was processed in chunks of *T* = 40,000 frames. The same initial ROIs were used for each chunk. For each neuron in each chunk, a region of 50×50 pixels centered on the neuron (the ‘context region’, C) was selected for further processing (**Fig. S22**). We high-pass filtered the data (MATLAB filtfilt) in the context region at 0.33 Hz using a 3^rd^ order Butterworth filter to correct for photobleaching. We denote the high-pass filtered movie as *D*_*N*×*T*_ where *N* = *n*(*C*) is the number of pixels in the context region. The raw data was also high-pass filtered at 60 Hz using a 3^rd^ order Butterworth filter; we denote this high-pass filtered movie as 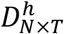. The initial temporal trace *X*_0_(*t*) was the mean of the 0.33 Hz high-pass filtered video over the pixels in the ROI (*R*):

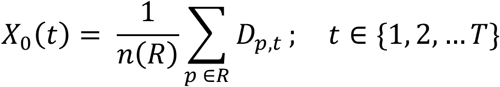

where *R* denotes the set of pixels in the ROI, *n*(*R*) is the number of pixels in the ROI.

*X*_0_ and the high-pass filtered videos *D*_*N*×*T*_ and 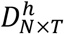 were provided as input to the Spike Pursuit algorithm, which consisted of a two-step loop for each iteration (*i*):

###### Step 1: Spike time estimation

To detect spikes in the initial trace, we first subtracted contributions from local background. This is intended to reduce the chance of optical crosstalk producing a false spike detection due to an adjacent neuron overlapping the initial ROI, and was not performed when computing the final trace with the optimized spatial filter. We defined the ‘local background’ (*B*) as the all pixels in the context region more than 12 pixels away from any pixel in the ROI, with *M* = *n*(*B*) pixels. We computed the SVD (singular value decomposition) of the background movie 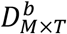:

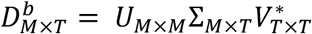

We then performed multiple linear regression of the trace *X*_*i*-1_ against the top eight background principal components:

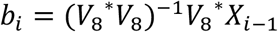

where *V*_8_ is the first eight columns of *V*; *b*_*i*_ are the regression coefficients. We denoised the trace *X*_*i*-1_(*t*) by subtracting the contribution of background pixels:

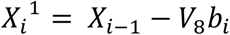

*X*_*i*_^1^(*t*) was high-pass filtered at 60Hz using a third order Butterworth filter. Local minima in the filtered trace below a threshold *s*_*i*_ were selected as an initial estimate of spike times. The threshold was chosen as follows: the distribution of local minima *p*_*min,i*_(*x*) was calculated by kernel density estimation and its median m was computed. The distribution of the noise *p*_noise,i_(*x*) was estimated by symmetrizing about the median; i.e. setting

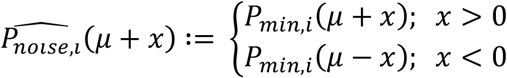

The distribution of spikes *P*_*spike*_ was estimated as:

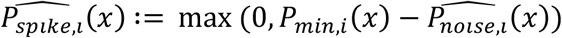

The threshold was selected as:

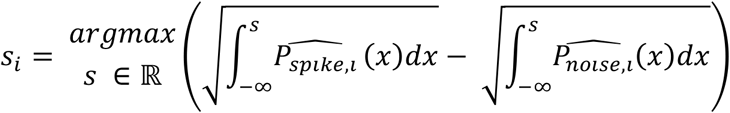

Thus, the initial estimate of spike times was *S*_*i*_ = {*t*|*X*_*i*_^1^(*t*) < *s*_*i*_, *X*_*i*_^1^(*t*) < *X*_*i*_^1^(*t* + 1), *X*_*i*_^1^(*t*) < *X*_*i*_^1^(*t* − 1)}.

This approach assumes that spikes only occur in a small proportion of time points, that *P*_*noise*_(*x*) is symmetric about *μ* in the absence of spikes, that local minima are uncorrelated to the voltage trace in the absence of spikes, and that no spikes produce local minima larger than *μ*. These assumptions are only approximately satisfied, but results of this method agree well with manual threshold selection.

Following the first round of spike detection, an action potential template *Z*_*i*_(τ) was generated as:

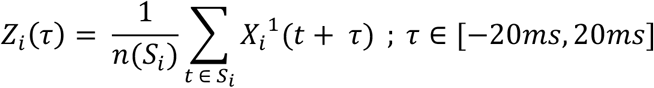

The template *Z*_*i*_(τ) was used to perform a whitened matched filter (*15*) on *X*_*i*_^1^(*t*), producing the temporally filtered trace 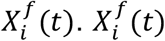 was again adaptively thresholded to obtain the estimated spike times 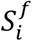 for iteration *i*, and regenerate the action potential template, 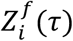.

###### Step 2: Spatial filter estimation

A target trace 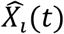 was produced by convolving the action potential template with the spike time indicator function:

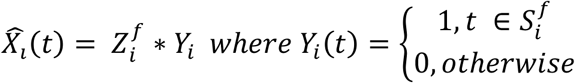

A spatial filter 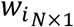 was estimated by ridge regression of the target trace 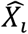 against 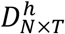 (**Fig. S22**).

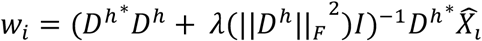

where ‖*D*^*h*^‖_*F*_ is the Frobenius norm of the high pass filtered data. The regularization parameter λ was selected by cross-validation on one dataset, and fixed for the remaining datasets. The activity trace corresponding to the spatial filter was calculated:

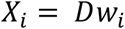

The Spike Pursuit loop was performed for five iterations. As a final step, we removed the contribution of pixels from a ‘global background’ (*G*), defined as the entire field of view excluding all pixels less than 12 pixels away from any ROI, with *L* = *n*(*G*) pixels. We computed the SVD of the global background movie high pass filtered at 0.3Hz, 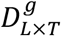:

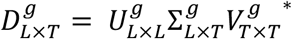

We performed multiple linear regression of the trace *X*_5_ against the top 8 principal components of the global background movie:

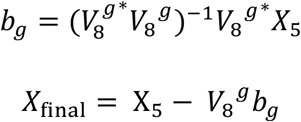

Global background subtraction removes fluorescence fluctuations that are shared across most pixels of the movie. It remains unclear to what degree these fluctuations reflect shared membrane potential transients versus other sources of shared variability. Traces without global background subtraction (*X*_5_) are shown in **Fig. S20**.

##### Calculating spike triggered averages for layer 1 interneurons

For each pair of neurons (*p*, *q*) the spike triggered average from neuron *p* to neuron *q* was calculated as:

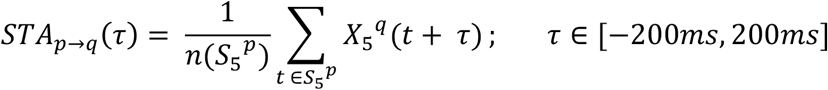

To calculate the shuffled distribution of spike triggered averages, each spike time was shifted by a random amount ranging from 2s to 4s (minimum of 2s chosen based on the typical autocorrelation function of the fluorescence traces). The spike triggered average of each pair was normalized by the standard deviation of the distribution of shuffled spike triggered averages (gray bar in **Fig. S21C**). We estimated the ‘modulation’ of one neuron by another as the L2 norm of the spike triggered average. To z-score this norm, we fit a log-normal distribution to its shuffle distribution. The background color in Fig. S21A and Fig. S21C represents this z-score.

We obtained similar results (not shown) for the spike triggered averages using raw fluorescence traces, calculated as:

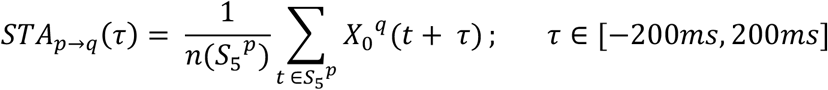

#### Transgenic zebrafish

Transgenic zebrafish which expresses Voltron under UAS promoter were generated as follows. A sequence of Voltron (Ace2-HaloTag) was cloned downstream of a 10×UAS sequence and the E1B minimal promoter (*16*). This plasmid was injected into 2-cell stage embryos of *Casper* mutant zebrafish (*17*) with mRNA of Tol2 transposase (*18*) to generate founder (F0) transgenic zebrafish.

Transgenic zebrafish which expresses Voltron under *elavl3* promoter for Figure S8 were generated as follows. A sequence of Voltron (Ace2-HaloTag) and its soma-localized variant (Ace2-HaloTag-SOM2) was cloned downstream of an *elavl3* promoter sequence. This plasmid was injected into 2-cell stage embryos of *Casper* mutant zebrafish (*17*) with mRNA of Tol2 transposase (*18*) to generate founder (F0) transgenic zebrafish. Images of the brains of their embryos (F1) were used for **Fig. S13**.

Experiments described in Fig. 4 **and S35** were performed using embryos generated by crossing the UAS.Voltron F0 founder and *vglut2a:Gal4* transgenic zebrafish (*19*) (a gift from Dr. Shin-ichi Higashijima). To label Voltron-expressing neurons with the accompanying fluorescent dye, 4-day old embryos were incubated in dye solution [3.3 μM JF_525_-HaloTag ligand (*20*) and 0.3 % DMSO] in fish rearing water at room temperature for two hours. After screening for the fluorescence of the JF dye in the brain, the fish were returned to fish rearing water with food until the time of the experiment.

#### Preparation for zebrafish imaging experiments

Imaging experiments were performed using 5- or 6-day larval zebrafish. The zebrafish was immobilized and mounted to an imaging chamber as described previously (*21*) with minor modifications. Briefly, the zebrafish were habituated in an artificial cerebrospinal fluid (ACSF) [in mM: 120 NaCl, 2.9 KCl, 2.1 CaCl_2_, 1.2 MgCl_2_, 20 NaHCO3, 1.25 NaHPO_4_, 10 Glucose] bubbled with carbogen gas (95% oxygen, 5% carbon dioxide) for 30 minutes. The muscle of the zebrafish was paralyzed by a short (up to 30 seconds) bath incubation with alpha-bungarotoxin (1 mg/ml, Thermo Fischer Scientific, B1601) dissolved in external solution. After the fish became immobile, heart movement of the zebrafish was stopped by microforceps to prevent the shadowing effect of blood cells in the brain during imaging experiments. The zebrafish showed robust optomotor behavior in ACSF (bubbled with carbogen gas for 30 minutes before the experiment) for several hours after this treatment. The zebrafish was further mounted to a custom-made chamber using 2% agarose (Sigma-Aldrich, A9414) and placed under a light-sheet microscope (*22*) with a 20× objective lens (Olympus, XLUMPLFLN).

#### Light-sheet imaging of zebrafish Voltron signals

Imaging was performed in a light-sheet microscope according to a published design (*23*) with modifications targeted at optimizing Voltron imaging. To increase the fraction of time the imaged cells were exposed to the excitation laser beam, the beam was expanded in the horizontal dimension using a pair of cylindrical lens (LJ1878L1-A (f = 10mm) and LJ1402L1-A (f = 40mm), Thorlabs). Imaging was performed using a 488 nm excitation laser (80 μW) and a 562/40 emission filter (Semrock, FF01-562/40) and with a frame rate of 300 frames/second recorded by a sCMOS camera (Hamamatsu, ORCA Flash4.0 v2). In this setup, the pixel dimension on the camera was 0.293 μm/pixel and the imaged neurons occupied an area of 150-200 pixels on the image.

#### Simultaneous cell-attached extracellular recordings and voltage imaging in zebrafish

We performed simultaneous electrophysiology and imaging of neurons expressing Voltron in *Tg(vglut2:Gal4);Tg(UAS:Voltron*) transgenic zebrafish as described previously (*21*) with a minor modification. Fire-polished borosilicate glass pipettes (Sutter, BF150-75-7.5) were pulled using a heat puller (Sutter, P1000). The tip of the pipette was further coated by quantum dots (Ocean Nanotech, QSR-600) using a previously reported method (*24*). The pipette resistance after the quantum dot coating was 10-12 MΩ.

The fish was bathed in an external solution and a small incision on the top of the head was made using a sharp glass needle. The pipette was filled with an external solution and inserted into the cerebellum of the brain using a micromanipulator (Sutter, MPC-200), and extracellular spiking signals were recorded from vGlut2-positive neurons in the dorsal part of the cerebellum using cell-attached extracellular recordings. We assume these neurons are eurydendroid neurons in the cerebellum, homologues of neurons in the deep cerebellar nuclei in mammalian brains (*25*), based on their previously described anatomical locations (*26*) and their expression of the *vglut2* gene (*26*). Signals from the pipette were amplified by an amplifier (Molecular Devices, AxoClamp 700B) and recorded by custom software written in C# (Microsoft) at 6 KHz. Optical signals from the same neurons were simultaneously imaged as described above.

#### Behavioral experiments in zebrafish

Recording of fictive swim signals and presentation of visual stimuli were performed as described previously (*21,22*). To record swim signals from the axonal bundles of spinal motoneurons in the tail, we attached a pair of large barrel electrodes to the dorsal left and right side of the tail. Signals were amplified by an amplifier (Molecular Devices, AxoClamp 700B) and recorded at 6 KHz using custom software written in C# (Microsoft). For synchronization between the swimming signals and neural activity imaging, camera trigger signals that initiate the acquisition of individual frames in the light-sheet microscope were recorded simultaneously with the swim signals. During the experiments, we projected red visual stimuli (red/black gratings with bars 2 mm thick) to the bottom of the fish chamber. The speed of the moving the visual stimulus alternated between 0 mm/s (stopped) and 2 mm/s (moving forward) every 10 seconds. Every trial (20 seconds) contained a stop period and a forward-motion period. In a subset of tested fish, the forward moving speed was changed from 2 mm/s to 0.5 mm/s every other trial. Swimming behavior was continuously recorded for a duration ranging from 6 minutes to 12 minutes (18 to 36 trials).

Signals from the electrodes were processed and individual swim events were detected according to a method described previously (*21*). Briefly, the raw signals were high-pass filtered, squared and smoothed by applying a Gaussian filter (σ = 3.3 milliseconds). The resulting traces were defined to be the swim signal, as shown in Fig. 4. Individual swim bouts were detected by finding the time points at which the swim signal crossed a threshold. This threshold was automatically set to lie just above a noise level based on a histogram of the swim signals (*22,27*).

#### Analysis of imaging dataset

The flow of the data processing is described below and in **Fig. S35B** and **C**. We provide custom Python scripts for this analysis on Github (https://git.io/vA2Ee).

##### Step 1. Image registration

Sequences of recorded images were corrected for horizontal drift during the imaging session at the subpixel level with a phase correlation algorithm (*28*) using a custom Python script and a GPU computing board.

##### Step 2. Segmentation of neurons

Individual neurons in the imaging field were segmented in a semi-automatic way. This was done using a combination of cell recognition by a pre-trained convolutional neural network build with the Python Keras library (https://keras.io/) and manual correction. This convolutional network discriminates whether a locally darkest point in a circular patch (radius = 2.67 μm) is a center of a Voltron-expressing neuron or not. Once the cells are segmented, ring-shaped masks are drawn automatically over the cells (1^st^ pixel weights in **Fig. S35B** and **C**). This is achieved by (1) selecting the brightest points on a line (at 0 degrees) from the center of a cell, (2) selecting such points for different angles around the cell (0 to 342 degrees in 18-degree steps), (3) smoothing the line connecting these brightest points by median filtering the distance from the center of the cell to the brightest points, and (4) dilating the resulting line by 1 pixel. Pixels on this dilated line are given an initial weight of 1, and all other pixels an initial weight of 0, to create the mask.

##### Step 3. Optimization of pixel weights for individual neurons

Weights on the pixels of the above masks were optimized to maximize the signal-to-noise ratio of the voltage signals in individual neurons (**Fig. S35C**). This is necessary because the light scattering through the tissue during the imaging experiment mixes to a small extent the optical signals across pixels surrounding the cell. This process optimizes the weights on the pixels over the cell to maximize the objective function *J*:

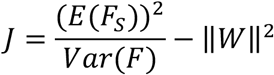

where *W* is a matrix of weights over pixels, *Var(F)* is a variance of a weighted mean fluorescence time-series of candidate pixels using *W*, *E(Fs)* is the average of the weighted mean fluorescent values at the time of detected spikes, and ‖*W*‖^2^ is the L^2^ norm of the pixel weights for regularization. This objective function measures the ratio between the mean heights of the spikes and the noise level of the estimated fluorescence time-series of a cell. Pixel weights ***W*** are optimized so that they maximize the objective function *J* using a gradient ascent method. Spiking events used for this optimization are detected using the fluorescence time series of the 1^st^ pixel weights using the same algorithm as described below. The final fluorescence time series is obtained by (1) calculating the weighted average of fluorescence values across pixels using the optimized ***W***, (2) subtracting the camera background (i.e., the pixel value when the camera records dark images), and (3) normalizing the resulting time-series by dividing by its baseline time-series, which is a rolling percentile (80% [since Voltron becomes dimmer with increasing voltage, an upper percentile was used instead of a lower percentile used for calcium imaging data], 500-ms time window) of the time-series.

##### Step 4. Spike detection

Lastly, spiking events were detected for individual neurons using an iterative method that first estimates the subthreshold potential and subtracts it from the raw voltage trace, and secondly estimates spiking events on the resulting trace. These three steps were iterated three times:

1. The subthreshold potential was obtained by subtracting the current estimate of the voltage trace attributable to spikes (i.e. the convolution of the estimated spike train ***s*** with spike shape ***k***, ***s*** * ***k***) from the raw trace followed by low pass filtering to remove the noise. Using a simple Butterworth filter of order 5 with a cutoff at 10 Hz was effective. Subtracting the subthreshold potential yielded the high-frequency component ***y*** that consists of voltage transients due to spikes corrupted by noise.
2. Spiking events were detected using a method based on adaptive template matching. First, large spiking events were detected using a high threshold (3.5 * rolling standard deviation + rolling median, window size of 3 seconds) to avoid false positives. The neuron’s spike shape was constrained to have non-zero values only in a small window around the time of a spike and was calculated using linear regression of ***y*** on ***s***.
3. The less clear spike events were detected using this mean spike shape *k* as a template instead of merely relying on a threshold. Potential spike candidates were detected using a low threshold (2.5 * rolling standard deviation + rolling median) to avoid false negatives. Template matching by regressing ***y*** on ***k*** yielded the sizes of these candidate events. The candidates that were not actual spikes but were merely due to noise had a small size and were iteratively removed with the regression being repeated. After the three outer iterations we obtained a reasonable estimate of the spike shape ***k*** and the spike times at frame-rate resolution.

##### Step 5. Validation of the authenticity of the detected spikes

To minimize the false-positive detection rate of spiking events, we measured the authenticity of the spike shapes throughout the time-series by measuring the gradient of the voltage trace just before the estimated spiking events. This is based on an assumption that spiking events are always preceded with an increase of subthreshold membrane voltage and that non-spike high-frequency noise does not have this preceding component. The gradient of voltage signals from 10 milliseconds to 3.3 milliseconds (3 time points) before the spiking events were quantified for individual spikes. Detected spiking events were binned into contiguous blocks of 50 spiking events. The gradient values for each block of spiking events were tested for its deviation from zero using a Wilcoxon signed rank test. Spiking events in blocks that had significantly positive gradients (p<0.05) are used for subsequent analysis.

#### Analysis of the relationships between neural activity and behavior

Neurons that were used for analysis in **Fig. 4** and **Fig. S35** were statistically selected based on their modulation of spiking activity by visual stimuli and behavior. We used two criteria for this. First, the difference of numbers of spikes in the two task periods (stop, forward visual motion) across multiple trials was tested using Wilcoxon’s ranked sum test. Second, the modulation of subthreshold signals at the initiation of swim bouts (−100 ms to 100 ms) was tested across all swim bouts using two-way analysis of variance. Neurons which showed significant differences (p<0.05) for both criteria were used for subsequent analyses. A total of 468 neurons from 81 fish were tested, and 179 neurons from 43 fish were used for subsequent analyses.

Mean subthreshold signals and firing rates on **Fig. 4E** and **S35E** were smoothed by gamma density causal filters (t*exp(-t/θ) for t>0) with a hyperparameter 0 set differently for each panel (3.3 ms for **Fig. 4E** and 200 ms for **Fig. S35E**).

For classifying neurons in **Fig. 4E**, subthreshold voltage signals were first smoothed by a gamma density causal filter (θ = 3.3 ms) and then averaged centered at the onset of all swim bouts (−100 to +100 milliseconds, 60 time points). The resulting averaged subthreshold signals were normalized to between 0 and 1 using their minimum and maximum values. Non-negative matrix factorization (NMF) was applied to the pool of these normalized subthreshold signals with a prior number of components set to 3. We confirmed that three components similar to the ones shown in **Fig. 4D** (‘Off’, ‘Onset’ and ‘Late’) always appear as NMF components regardless of the initial conditions. Component weights for each of the neurons are further adjusted so that the sum of weights for three components becomes 1, and these adjusted weights are allocated to red, green and blue channels to color neurons.

#### Widefield imaging of Voltron expressing neurons in zebrafish

*Mitfa*^w2/w2^ *roy*^a9/a9^ (Casper) zebrafish were maintained under standard conditions at 28 °C and a 14:10 hr light:dark cycle. Embryos (1-2 cell stage) of Tg(elavl3:Gal4-VP16) were injected with 25 ng/μl DNA plasmid encoding the Voltron-ST indicator under the control of the 10xUAS promoter, and 25 ng/μl Tol2 transposase mRNA diluted in E3 medium. Subsequently, the injected embryos at three day post-fertilization (dpf) were incubated in system water containing JF dye-HaloTag ligands (JF_525_, JF_549_, JF_585_ or JF_635_) at 3 μM for 2 hours and then washed in dye-free system water. Larvae at 5 or 6 dpf were screened for the expression of the Voltron-ST based on the fluorescence from JF dye-HaloTag ligands. They were paralyzed by 5-min bath application of 1 mg/ml a-bungarotoxin (Sigma, 203980) and mounted dorsal-side up with 1.5% low-melting point agarose. Spontaneously active forebrain and olfactory neurons were imaged using a custom widefield microscope. The objective was a 60x 1.0 NA water immersion lens (Nikon, MRD07620). Fluorophores were excited with a LED light source (Luminus, CBT-90-W for JF_525_, JF_549_ and JF_585_; Luminus, CBT-90-RX for JF_635_) with a proper filter set (Semrock, FITC-A-Basic-000 for JF_525_; Semrock, Cy3-4040C-000 for JF_549_; Semrock, LED-mCherry-A-000 for JF_585_; Semrock, LF635-C-000 for JF_635_). The images were acquired with sCMOS camera (PCO, pco.edge 4.2) at 400 Hz (2.5 ms exposure time) for 1-2 min. Data were analyzed using MATLAB (Mathworks). Regions of interest (ROIs) corresponding to identifiable cell bodies were selected manually and the mean signal from each ROI was extracted. The baseline was estimated by fitting the raw fluorescence time course with an exponential curve to account for bleaching. The estimated baseline was used to calculate the ΔF/F_0_.

#### Simultaneous whole-cell recording and Voltron imaging in zebrafish

Experiments were performed on 6-day old progeny of a cross between Tg(10xUAS:Voltron) and TgBAC(slc17ab:LOXP-mCherry-LOXP-GAL4FF;vsx2:Cre). Fish were loaded with JF525 and then paralyzed as described above. After anesthetizing fish with MS-222, they were head-fixed and prepared for whole-cell recording and imaging of V2a hindbrain neurons. They were secured to a Sylgard-coated glass-bottom dish containing extracellular solution (134 mM NaCl, 2.9 mM KCl, 1.2 mM MgCl_2_, 2.1 mM CaCl_2_, 10 mM HEPES, and 10 mM glucose, adjusted to pH 7.8 with NaOH) with etched tungsten wires through the notochord. Then the head was rotated and secured ventral side up with etched tungsten pins placed through the ears and the rostral part of the jaw. The ventral surface of the hindbrain was carefully exposed by removing the notochord using an etched tungsten pin and fine forceps. Whole-cell recordings were guided based on fluorescence image and scanned Dodt gradient contrast image acquired with a custom two-photon microscope equipped with 40× 0.8 NA objective lens (Nikon, MRD07420). Borosilicate glass pipettes (Sutter, BF150-86-15) were pulled by a micropipette puller (Sutter, P-1000) and filled with intracellular solution (125 mM potassium gluconate, 2.5 mM MgCl_2_, 10 mM EGTA, 10 mM HEPES and 4 mM ATP-Na_2_ adjusted to pH 7.3 with KOH). The resistance of the pipette was 5 to 7 MOhm. Recordings were made using the EPC 10 Quadro amplifier and PatchMaster (HEKA instruments). Voltron signal was acquired as described above but with 40x objective lens. After extracting Voltron signal from the patched cell using the procedure described above, the signal was further denoised using *wden* function in Wavelet Toolbox in MATLAB (Mathworks) to reveal Voltron signal corresponding to small subthreshold voltage changes.

#### Simultaneous Voltron imaging and whole-cell patch clamp in live adult flies

Experiments were performed on 2- to 10-day-old heterozygous progeny of a cross between UAS-IVS-syn21-Voltron-p10 and MB058B-Gal4 (*29*). The cross was kept on standard cornmeal food supplemented with all-trans-retinal (0.2 mM before eclosion and then 0.4 mM). Flies were head-fixed and prepared for imaging and electrophysiology as described previously (*30*). A small window was opened on the head cuticle, and fat cells and trachea that overlaid the target region were removed. The exposed brain was bathed in a drop (~200 μl) of dye-containing saline (1 μM for JF_549_ and 5 μM for JF_525_) for 1 hr. Saline contains (in mM): NaCl, 103; KCl, 3; CaCl_2_, 1.5; MgCl_2_, 4; NaHCO_3_, 26; N-tris(hydroxymethyl) methyl-2-aminoethane-sulfonic acid, 5; NaH_2_PO_4_, 1; trehalose, 10; glucose, 10 (pH 7.3 when bubbled with 95% O2 and 5% CO2, 275 mOsm. The brain was then washed with fresh saline several times, and maintained in the saline for 1 hr. During the dye application and washout, animals were placed in a moist chamber to avoid dehydration. After that, they were moved to the imaging rig, where superfusion continued at 1-2 mL/min with oxygenated saline. To minimize movement during imaging, the proboscis was fixed with a UV-curable glue (NOA 68T, Norland products) and the frontal pulsatile organ muscle 16 was removed. Imaging was performed on a wide-field fluorescence microscope (SOM, Sutter Instruments) equipped with a 60X, NA 1.0, water-immersion objective (LUMPlanFl/IR; Olympus) and a sCMOS camera (Orca Flash 4.0 V2+, Hamamatsu). Images were acquired at 800 Hz with 4×4 binning through the Hamamatsu imaging software (HCImage Live). Data presented used JF_549_. Illumination was provided by a 530 nm LED (SA-530, Sutter) with an excitation filter (FF01-543/22-25, Semrock); intensity at the sample plane was ~5 mW/mm^2^ for axons and dendrites, and 8-16 mW/mm^2^ for soma; emission was separated from excitation light using a dichroic mirror (FF562-Di03-25×36, Semrock) and an emission filter (LP02-568RU-25, Semrock). Experiments with JF_525_ tended to yield shorter duration imaging sessions (~2 min versus >5 min for JF549 in dopamine neurons), likely because of greater phototoxicity with the shorter wavelength light. For JF_525_, illumination was provided by a 506 nm LED (SA-506-1PLUS, Sutter) with an excitation filter (FF01-503/40-25, Semrock); intensity at the sample plane was typically 10-25 mW/mm^2^; emission was separated from excitation light using a dichroic mirror (Di02-R532-25×36, Semrock) and an emission filter (FF01-562/40-25, Semrock).

Whole-cell recordings (*31*) were guided by Voltron fluorescence from target cells. The patch pipettes were pulled for a resistance of 5-7 MΩ and filled with pipette solution containing (in mM): L-potassium aspartate, 125; HEPES, 10; EGTA, 1.1; CaCl_2_, 0.1; Mg-ATP, 4; Na-GTP, 0.5; biocytin hydrazide, 13; with pH adjusted to 7.3 with KOH (265 mOsm). Recordings were made using the Axon MultiClamp 700B amplifier (Molecular Devices). Cells were held at around −60 mV by injecting hyperpolarizing current (< 50 pA). Signals were low-pass filtered at 5 kHz and digitized at 10 kHz.

Voltron data were analyzed in MATLAB. Regions of interest (ROIs) corresponding to different neuron compartments were manually selected, and the mean intensity of the ROI was extracted. Median filtering with a 50-ms time window was performed on the raw fluorescence traces to get a filtered trace, and F_0_ was calculated as the mean over the first 1 s of imaging session. For detecting action potential spikes and quantifying SNR, the filtered trace was subtracted from the raw trace. Spikes were detected by finding local minima with peak amplitude over 3.5 times the standard deviation of the entire subtracted trace, and SNR was quantified as peak amplitude over the standard deviation of the trace excluding the time zone (50 ms) containing spikes. To analyze the axon signals, the ROIs of ipsilateral and contralateral axons were first pooled together to detect spikes. The spikes were then assigned to either the patched cell or its sister cell depending on the relative peak amplitude, i.e. if ipsilateral / contralateral > 1, spike is assigned to the patched cell, otherwise it is assigned to the sister cell.

---

**Supplementary Figure 1.**
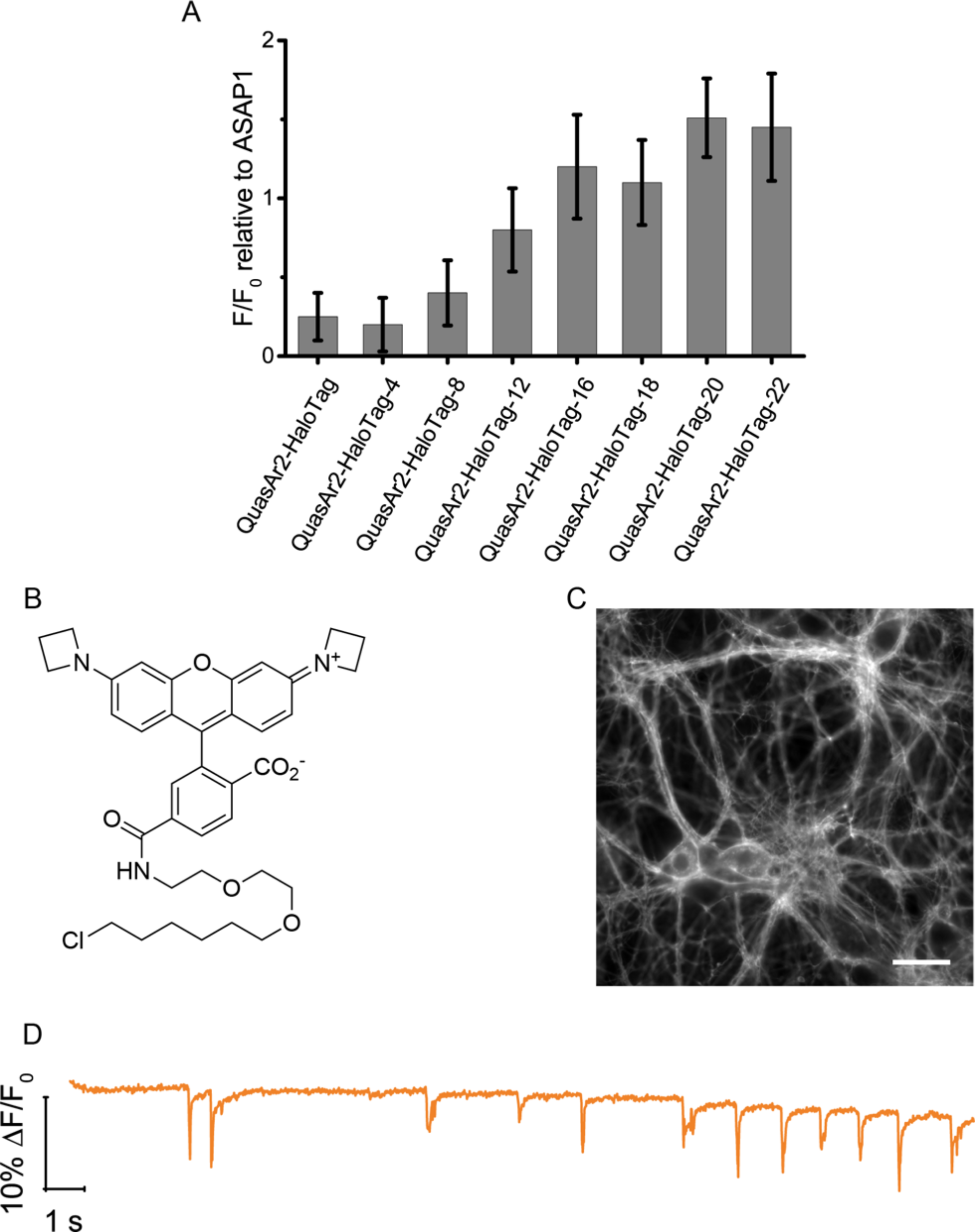
(A) To screen for the optimal linker length between the rhodopsin and self-labeling tag domains, QuasAr2-HaloTag fusions (labeled with JF_549_) and ASAP1 (*32*) were co-transfected into neurons and stimulated using a field stimulation electrode (see methods section). Truncating residues from the C-terminus of QuasAr2 and the N-terminus of HaloTag led to indicators with improved voltage sensitivity. Bar graph shows number of residues truncated from 0 - 22 amino acids. (B) Chemical structure of JF_549_. (C) Image of primary neuron cells expressing Voltron labeled with JF_549_. Scale bar: 20 μm (D) Imaging of spontaneous activity in neuron culture with QuasAr2-HaloTag-16 labeled with JF_549_.

**Supplementary Figure 2.**
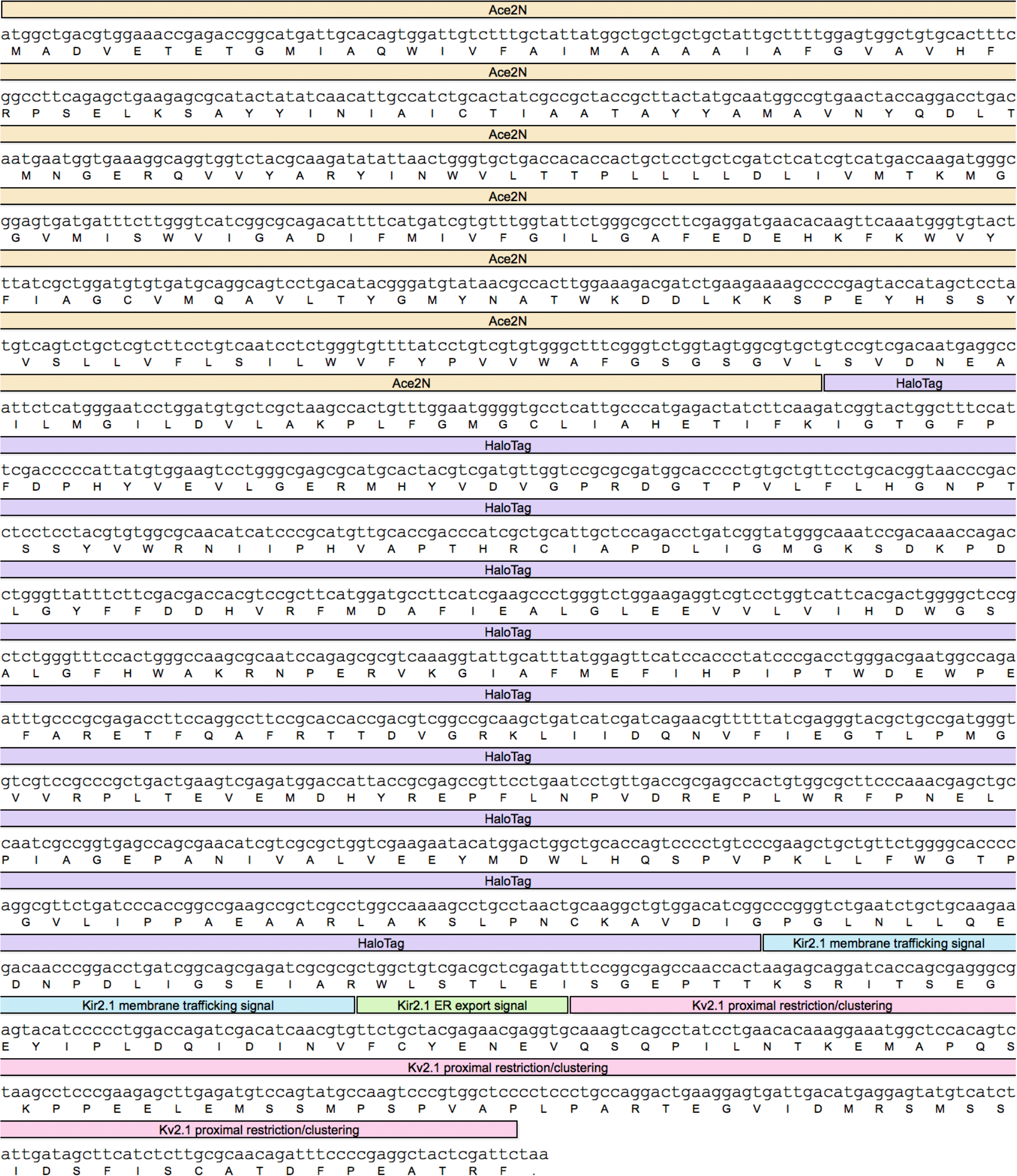
DNA and amino acid sequence of Voltron, with sequence features annotated. To localize Voltron to the neuron soma, a targeting sequence from Kv2.1 is added to the C-term of Voltron as shown above.

**Supplementary Figure 3.**
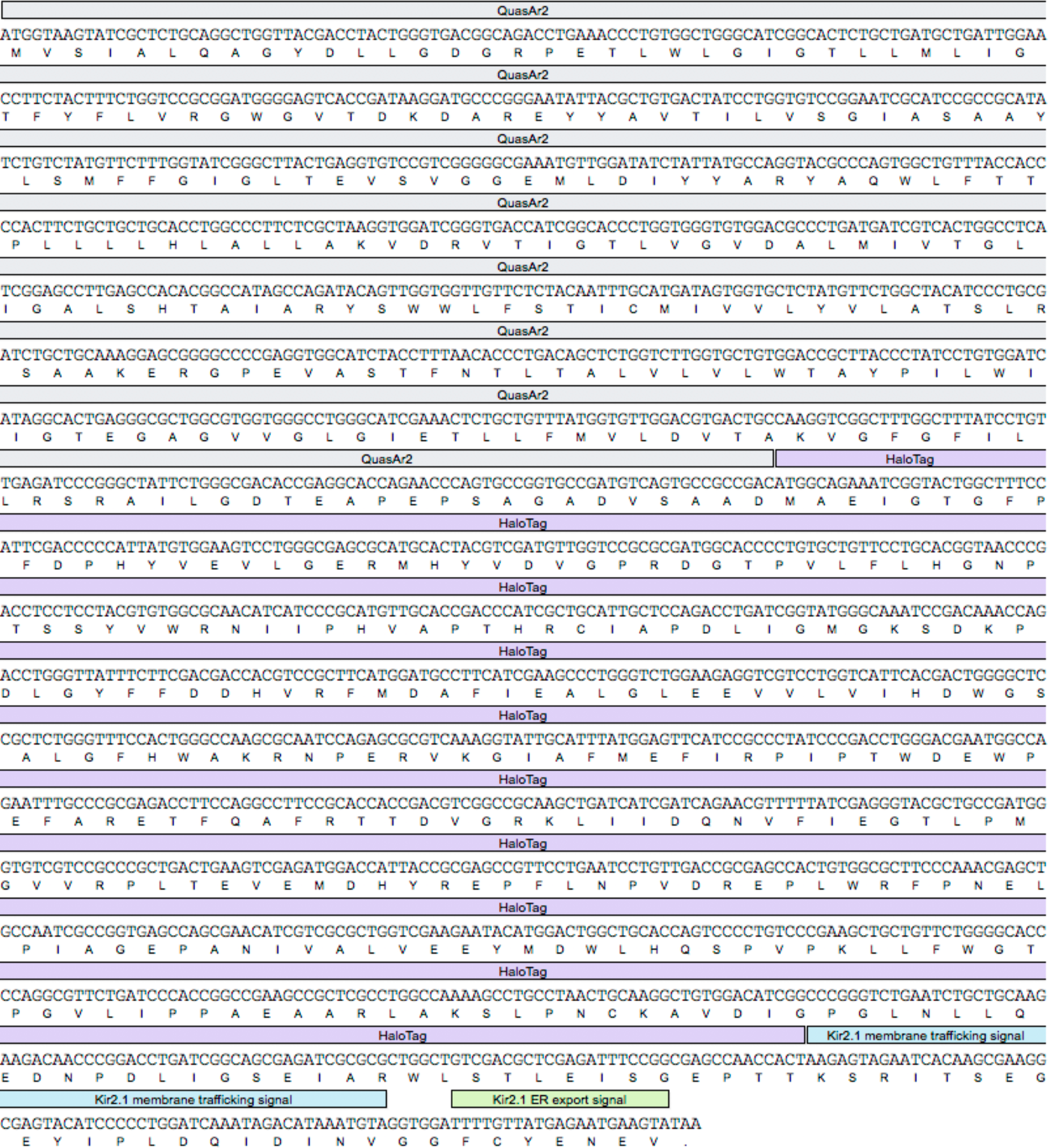
DNA and amino acid sequence of QuasAr2-HaloTag, with sequence features annotated.

**Supplementary Figure 4.**
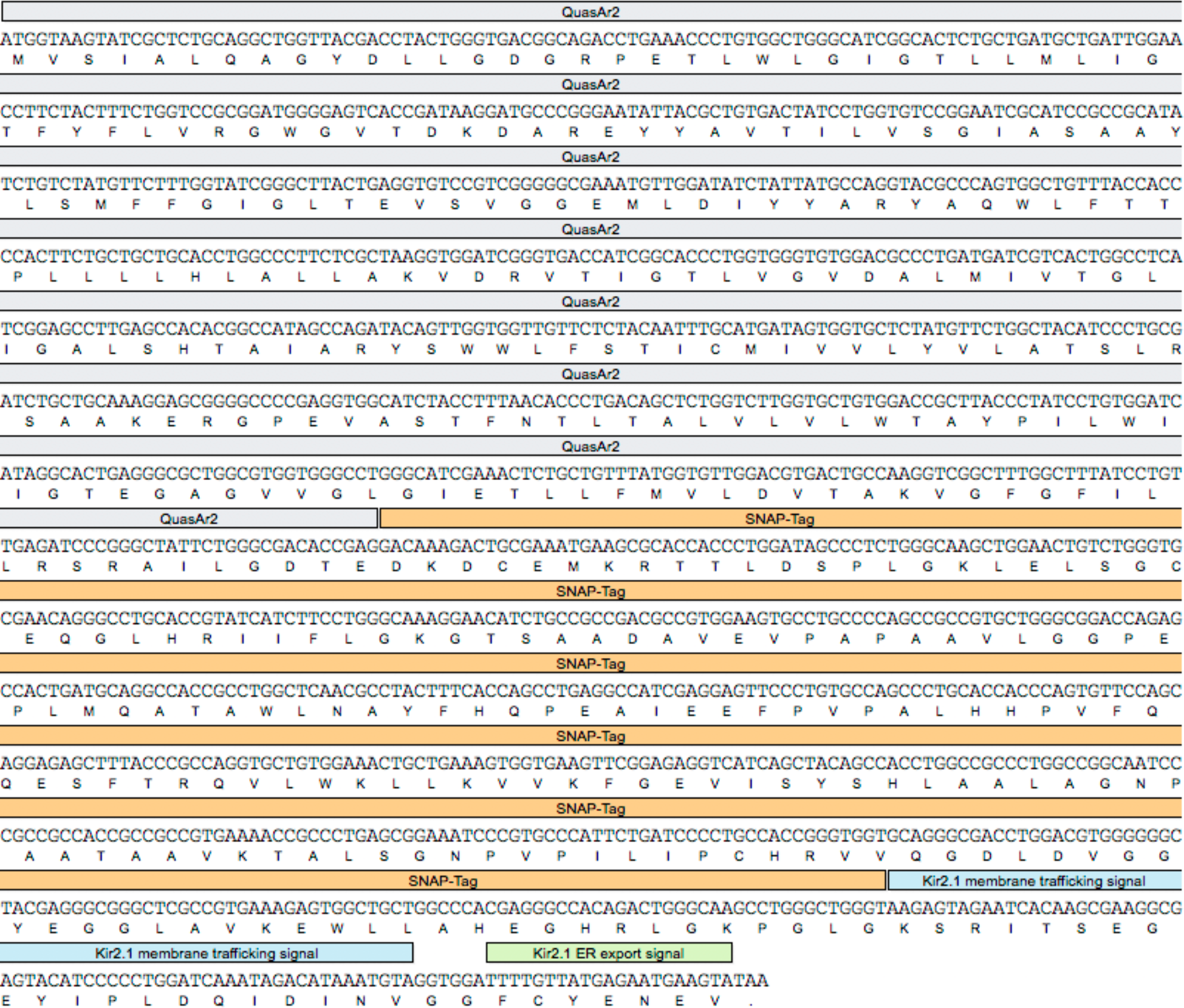
DNA and amino acid sequence of QuasAr2-SNAP-Tag, with sequence features annotated.

**Supplementary Figure 5.**
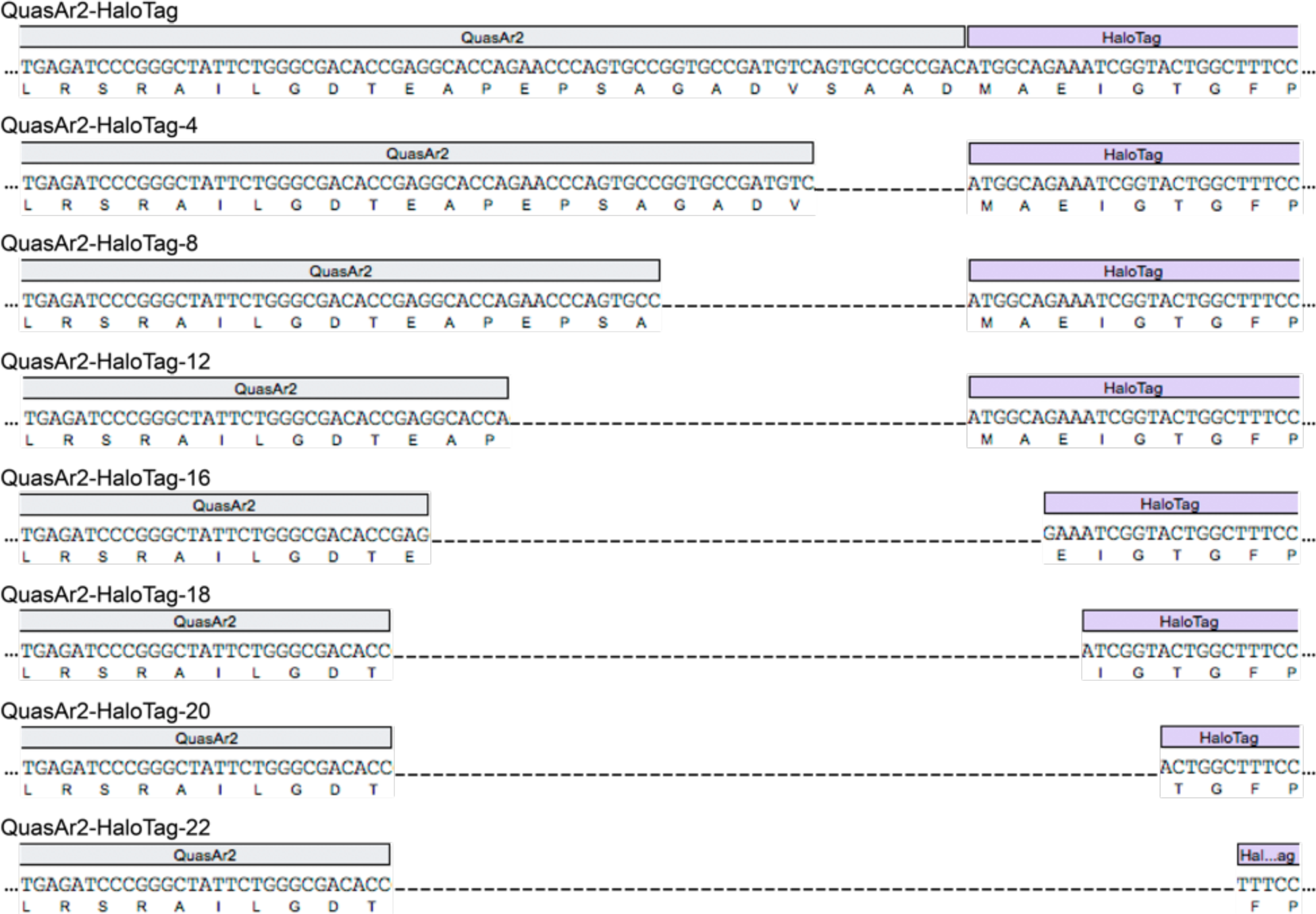
DNA and amino acid sequence of QuasAr2-HaloTag linker length truncations, with
sequence features annotated.

**Supplementary Figure 6.**
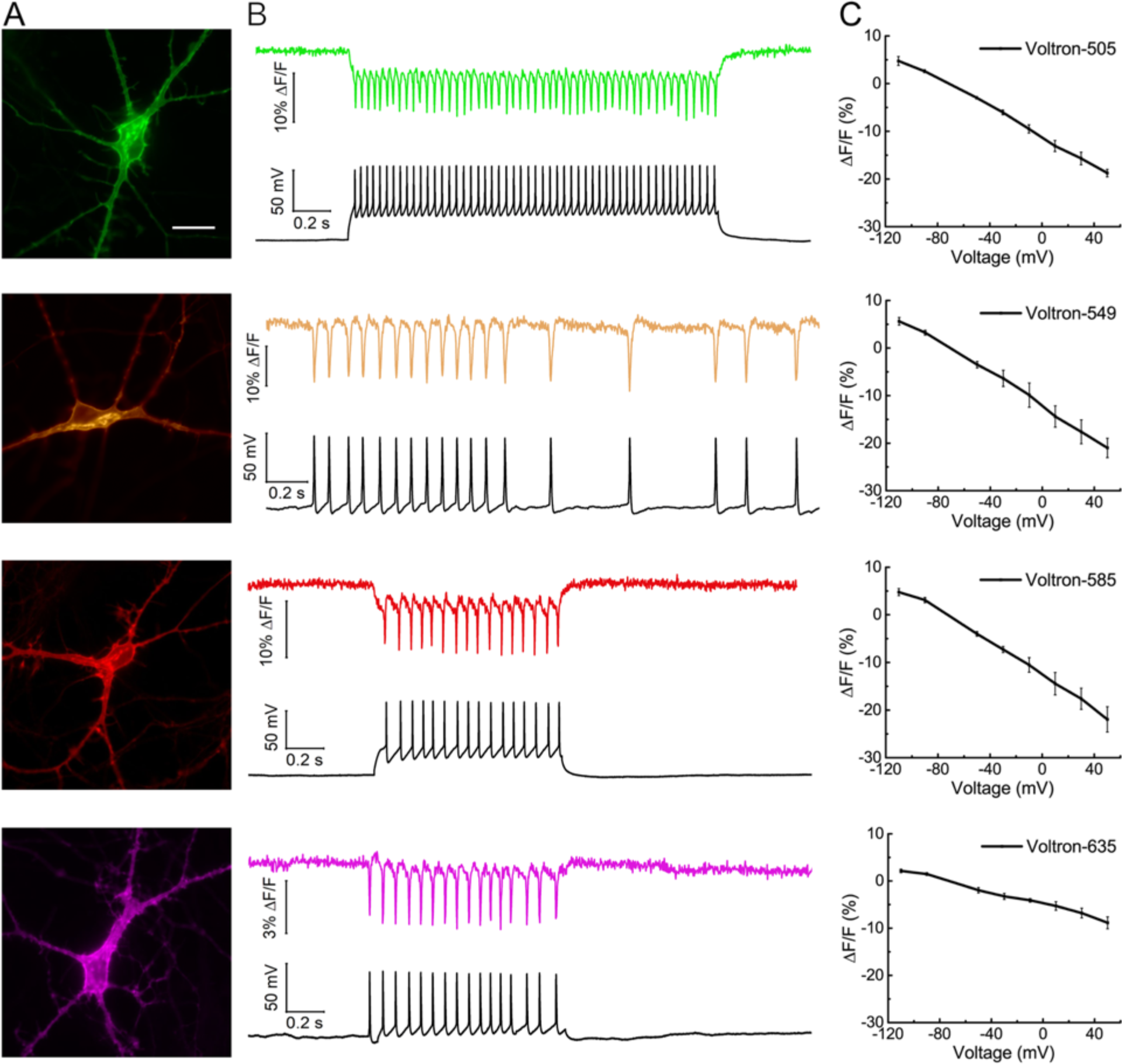
(A) Neurons expressing Voltron and labeled with different JF-HaloTag dye conjugates (Top to bottom: JF505, JF549, JF585, JF635) Scale bar: 20 μm (B) Single-trial recording of action potentials and subthreshold voltage signals from current injections. Raw fluorescence traces, colored by dye emission, are shown on the top of each panel. Membrane potential, recorded with a patch pipette, is shown at the bottom of each panel in black. (C) Voltron fluorescence change as a function of membrane voltage with different JF dye conjugates.

**Supplementary Figure 7.**
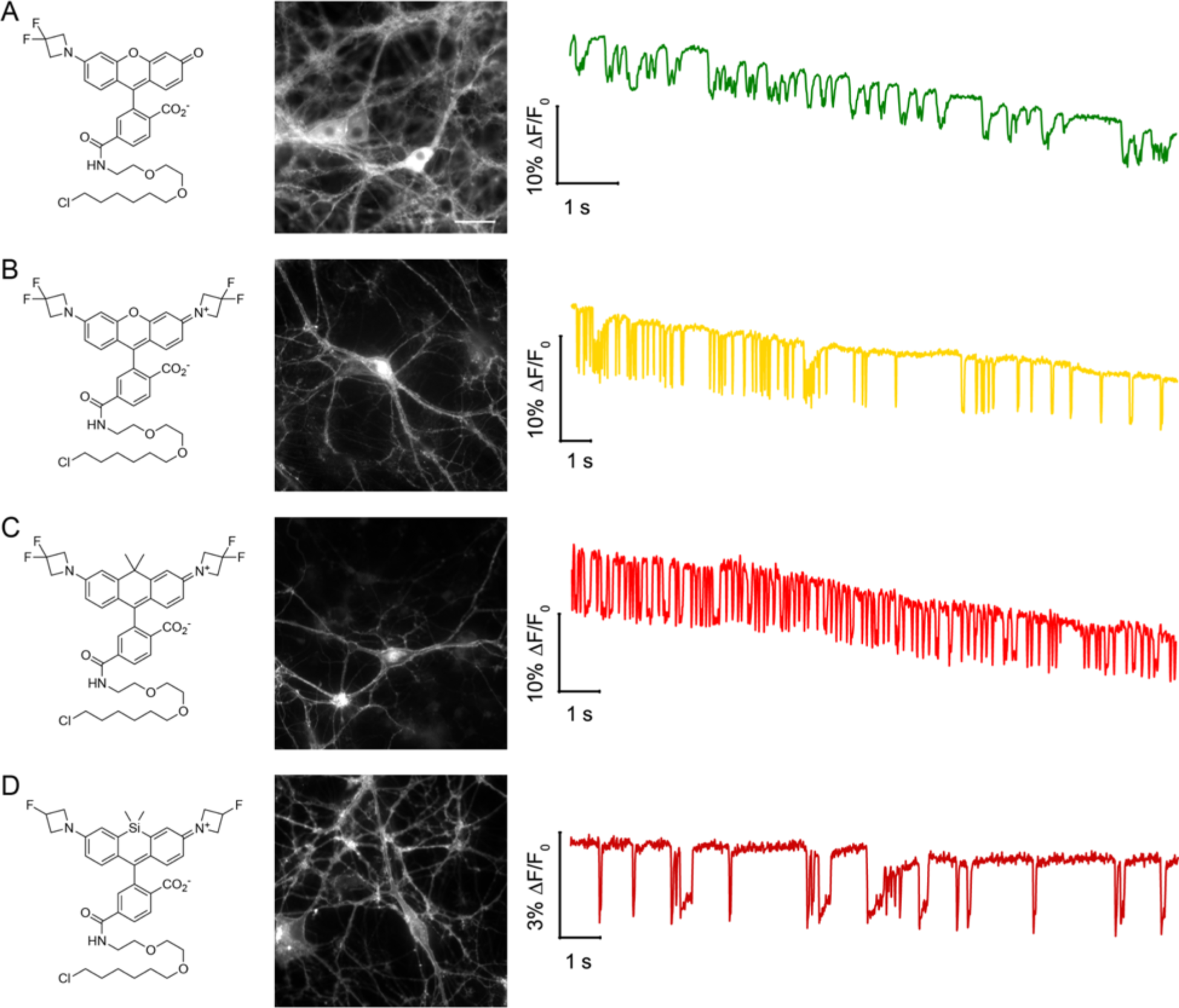
(A) Left: Chemical structure of JF505-HaloTag ligand, Middle: fluorescence image of hippocampal neurons in culture expressing QuasAr2-HaloTag-16 labeled with JF503. Right: Fluorescence trace over time showing voltage-dependent fluorescence changes resulting from spontaneous action potentials of the neurons. (B) Left: Chemical structure of JF525-HaloTag ligand, Middle: fluorescence image of hippocampal neurons in culture expressing QuasAr2-HaloTag-16 labeled with JF525. Right: Fluorescence trace over time showing voltage-dependent fluorescence changes resulting from spontaneous action potentials of the neurons. (C) Left: Chemical structure of JF585-HaloTag ligand, Middle: fluorescence image of hippocampal neurons in culture expressing QuasAr2-HaloTag-16 labeled with JF585. Right: Fluorescence trace over time showing voltage-dependent fluorescence changes resulting from spontaneous action potentials of the neurons. (D) Left: Chemical structure of JF635-HaloTag ligand, Middle: fluorescence image of hippocampal neurons in culture expressing QuasAr2-HaloTag-16 labeled with JF635. Right: Fluorescence trace over time showing voltage-dependent fluorescence changes resulting from spontaneous action potentials of the neurons. Scale bar: 20μm

**Supplementary Figure 8.**
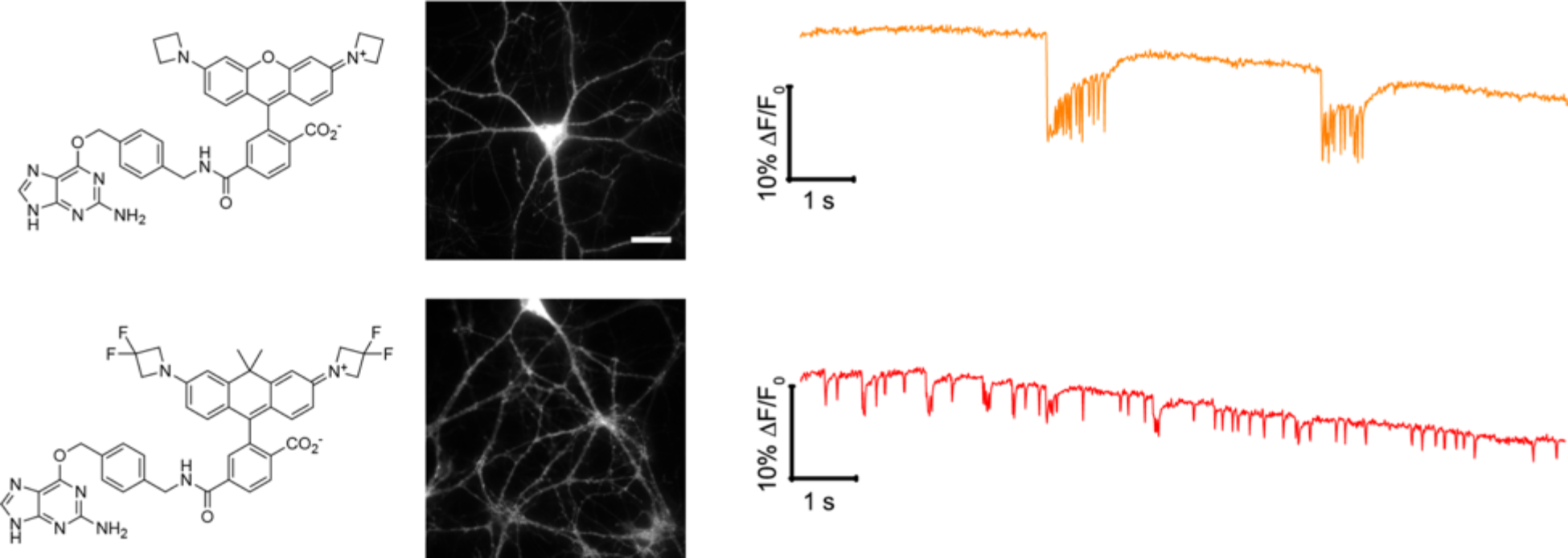
(A) Left: Chemical structure of JF_549_-SNAP-tag ligand, Middle: fluorescence image of hippocampal neurons in culture expressing QuasAr2-SNAP-tag labeled with JF_549_. Right: Fluorescence trace over time showing voltage-dependent fluorescence changes resulting from spontaneous action potentials of the neurons. (B) Left: Chemical structure of JF_585_-SNAP-tag ligand, Middle: fluorescence image of hippocampal neurons in culture expressing QuasAr2-SNAP-tag labeled with JF585. Right: Fluorescence trace over time showing voltage-dependent fluorescence changes resulting from spontaneous action potentials of the neurons. Scale bar: 20μm

**Supplementary Figure 9.**
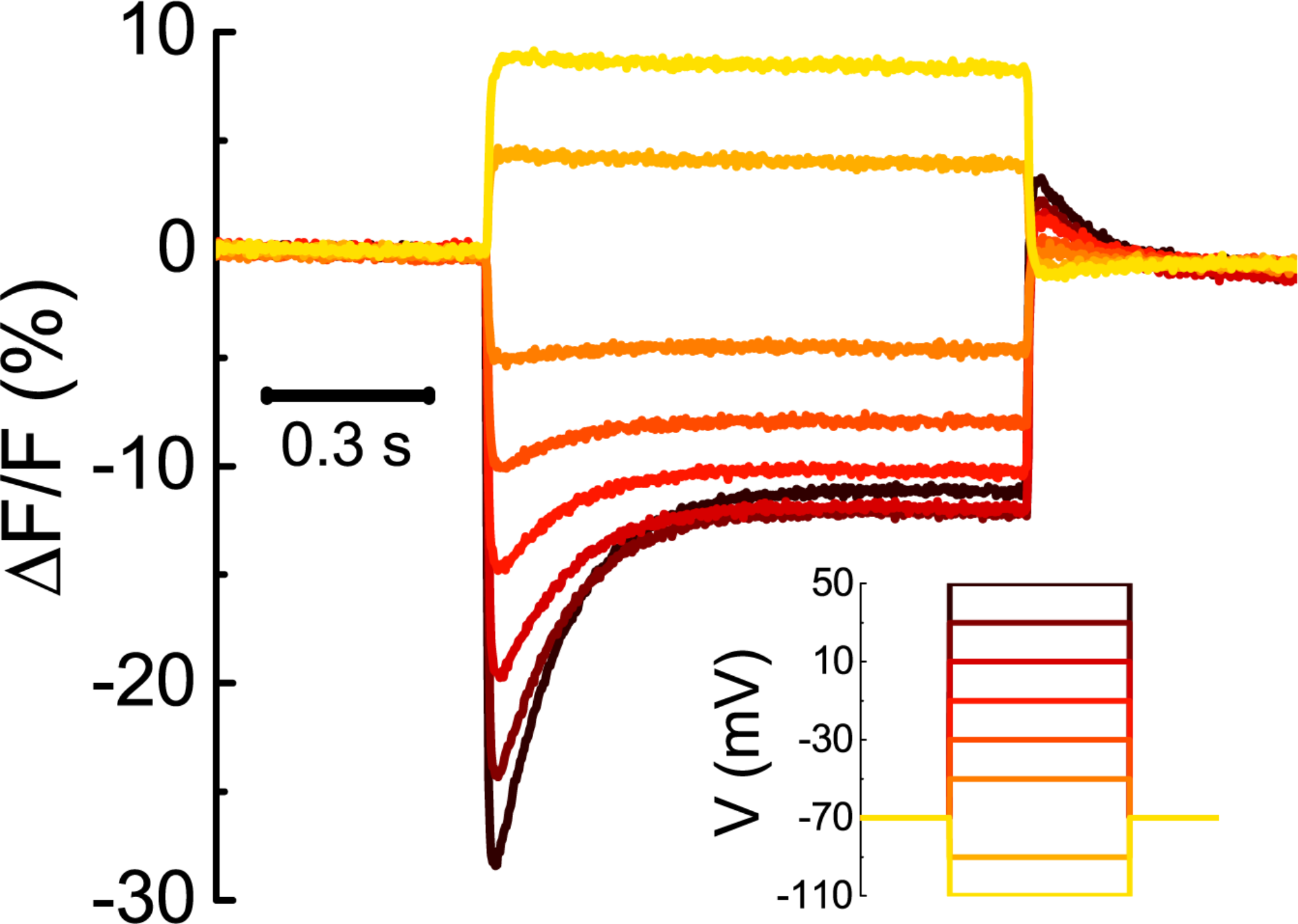
Representative fluorescence traces of Voltron_525_ in response to a series of voltage steps (from −110 mV to +50 mV in 20 mV increments). Image acquisition rate = 400 Hz. For complete fluorescence vs voltage plots for different JF dyes, see Fig.1E.

**Supplementary Figure 10.**
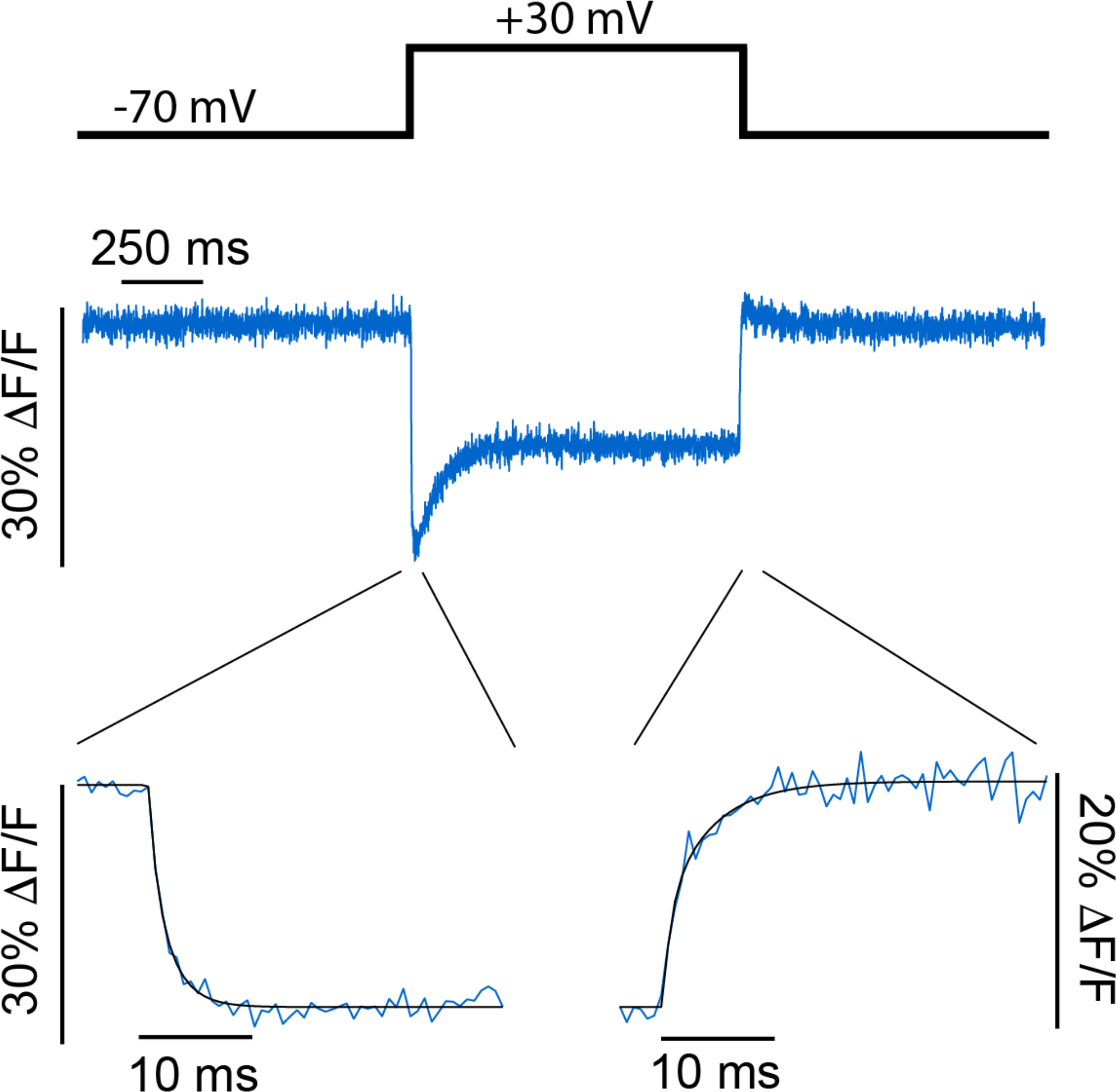
Representative fluorescence response of Voltron_525_ in a cultured neuron to a 100 mV potential step delivered in voltage clamp. Insets: Zoom in on change of Voltron fluorescence to depolarization and hyperpolarization. Solid black line is double exponential fit according to ΔF/F(t) = Ae^−t/τfast^ + Be^−t/τslow^. Image acquisition rate 3.2 kHz. For full kinetics data, see Table 1.

**Supplementary Figure 11.**
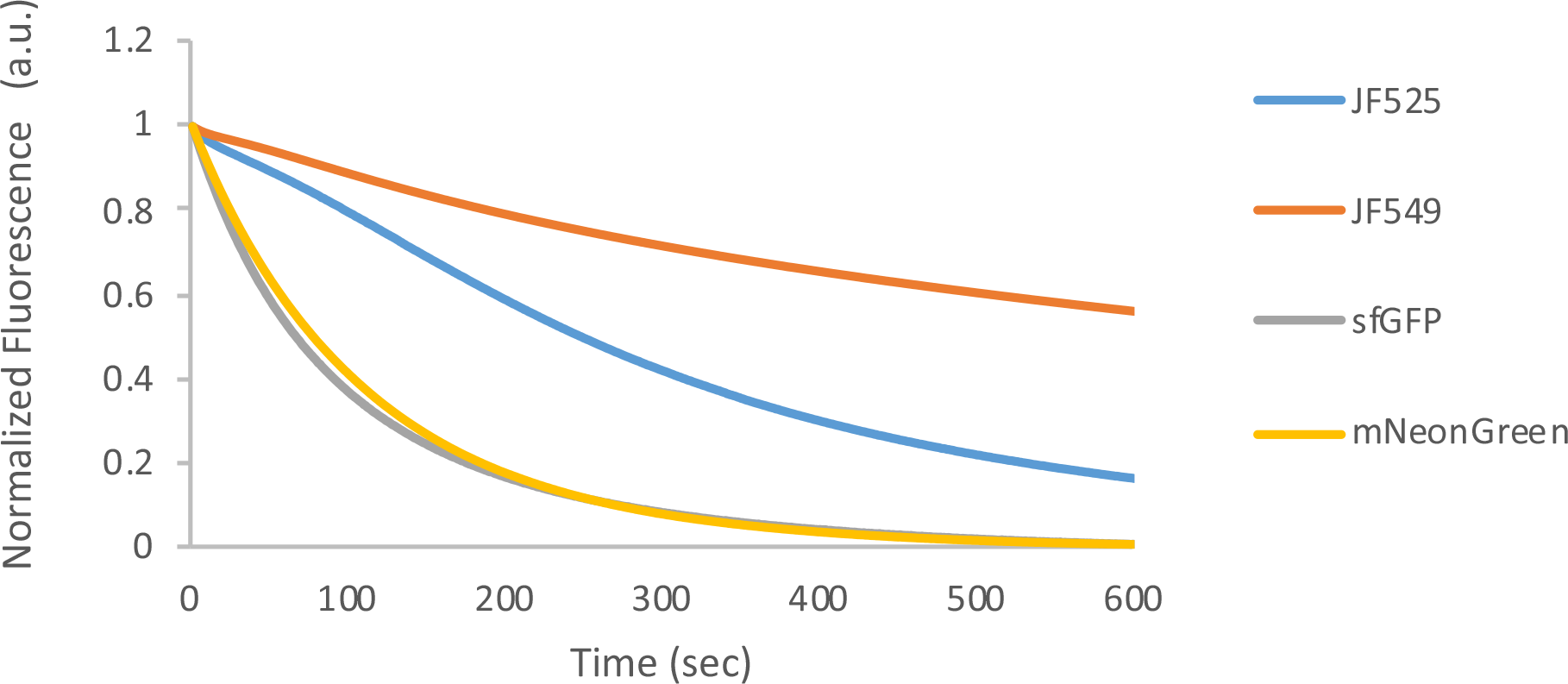
Photobleaching profile of HaloTag-bound JF525, JF549 and fluorescent proteins sfGFP and mNeonGreen, measured in aqueous droplets with widefield microscopy. Data taken at excitation rates W for JF525 (W = 1549 s^−1^), JF549 (1540 s^−1^), sfGFP (1597 s^−1^) and mNeonGreen (1772 s^−1^).

**Supplementary Figure 12.**
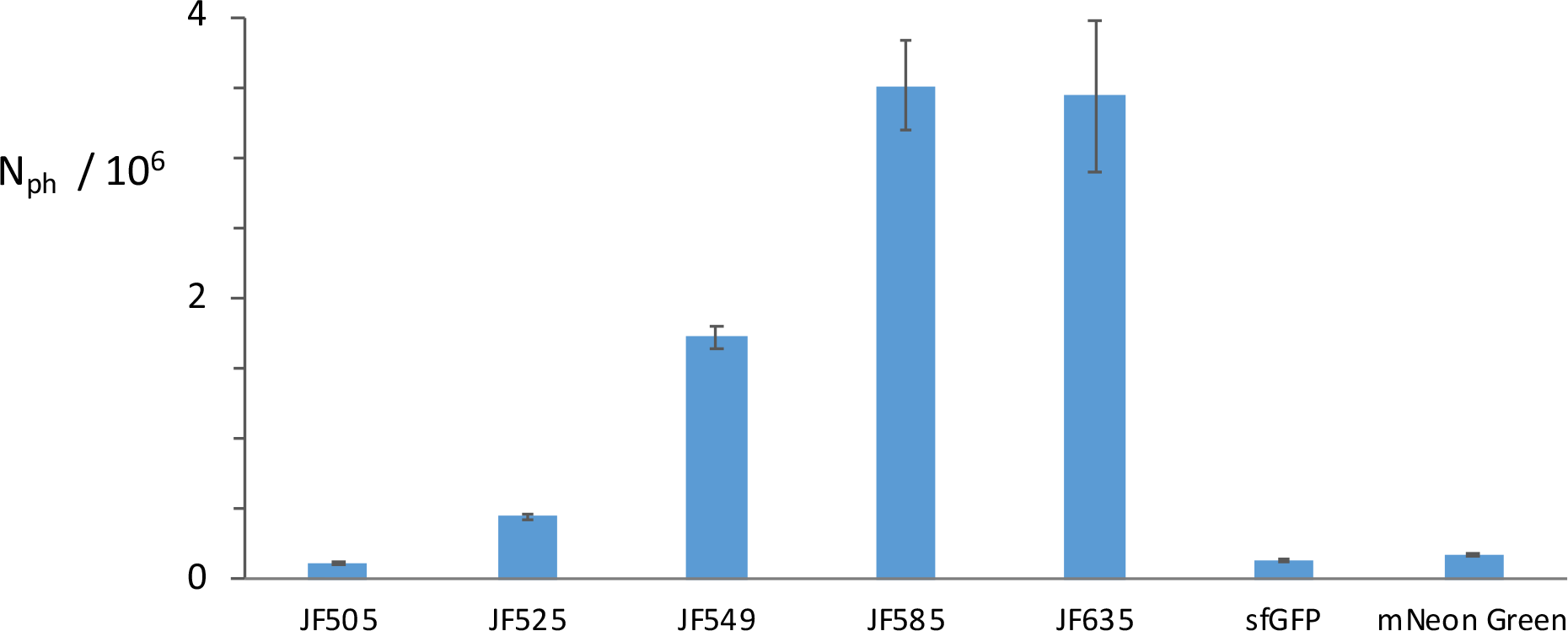
Mean number of photons per molecule emitted before photobleaching N_ph_ for JFdyes-HT conjugates and fluorescent proteins obtained in aqueous droplets using widefield microscopy. Error bars are standard deviation (n = 9).

**Supplementary Figure 13.**
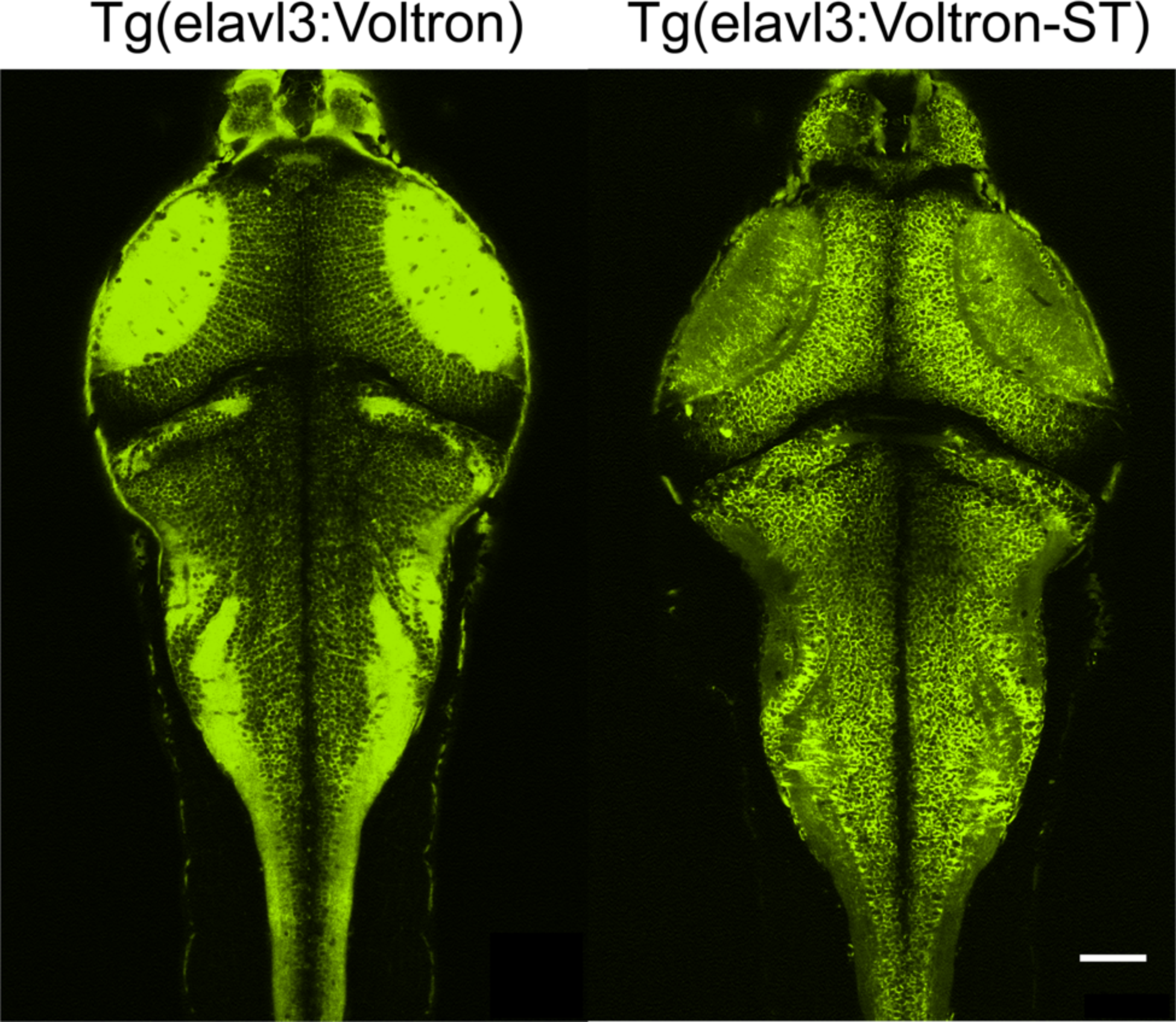
Voltron expression in zebrafish. A. Confocal image of Tg[elavl3:Voltron] (left) and Tg[elavl3:Voltron-ST] (right) zebrafish (4 dpf) labeled with JF525 dye. Scale bar: 50 μm.

**Supplementary Figure 14.**
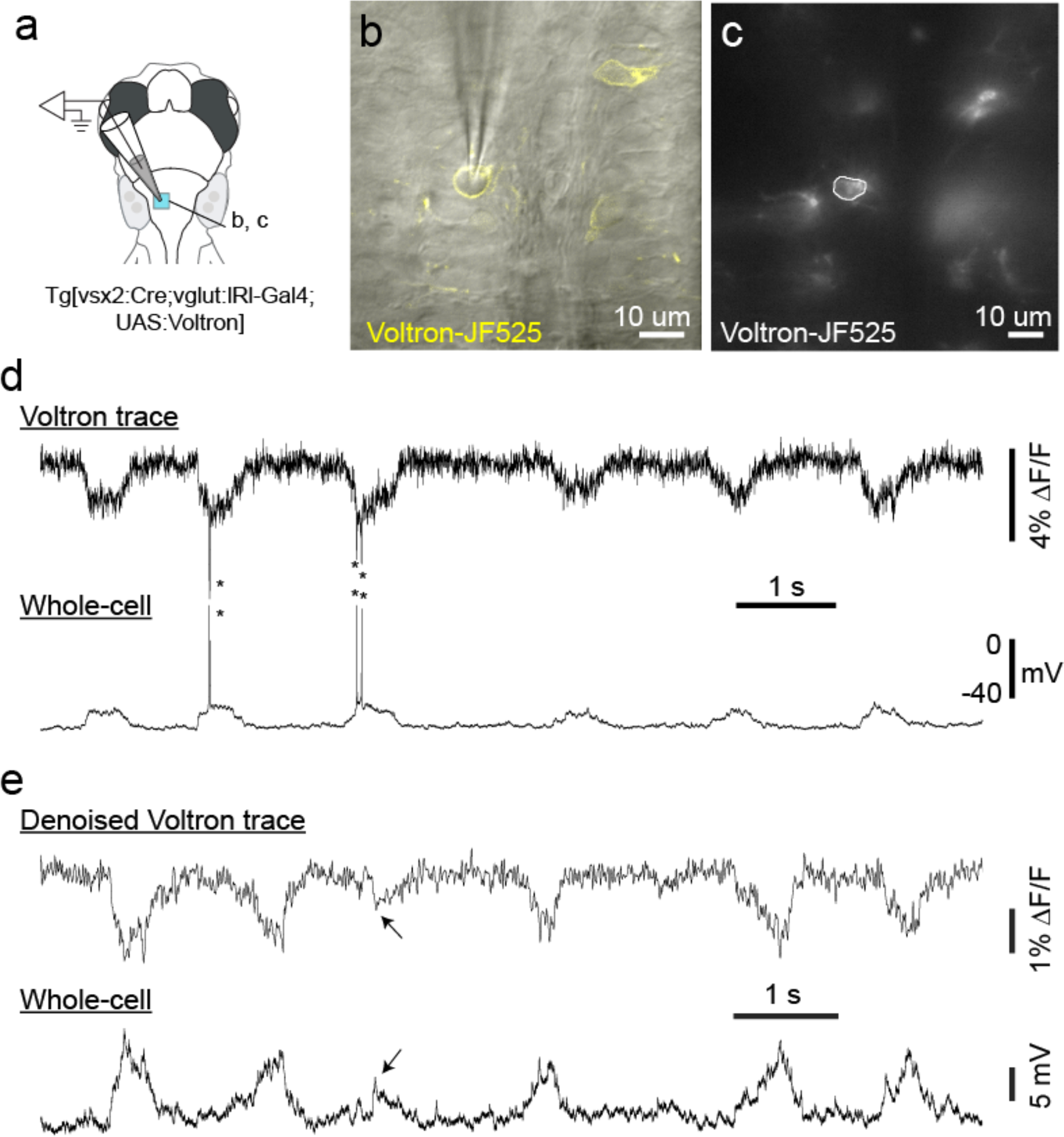
Simultaneous whole-cell recording and Voltron imaging in zebrafish **(A)** A schematic drawing showing the configuration of the experiment and the location of the images in the subsequent panels. **(B)** A single plane of two-photon image of the patched cell expressing Voltron (yellow) in the background of scanned Dodt gradient contrast (grey). **(C)** A widefield image showing the location of the region of interest used to extract Voltron signal in the following panels. **(D)** Traces of Voltron fluorescence (above) and whole-cell recording (below). Spikes are indicated by asterisks. **(E)** Traces of denoised Voltron fluorescence (above) and whole-cell recording (below). A small subthreshold event is indicated by an arrow.

**Supplementary Figure 15.**
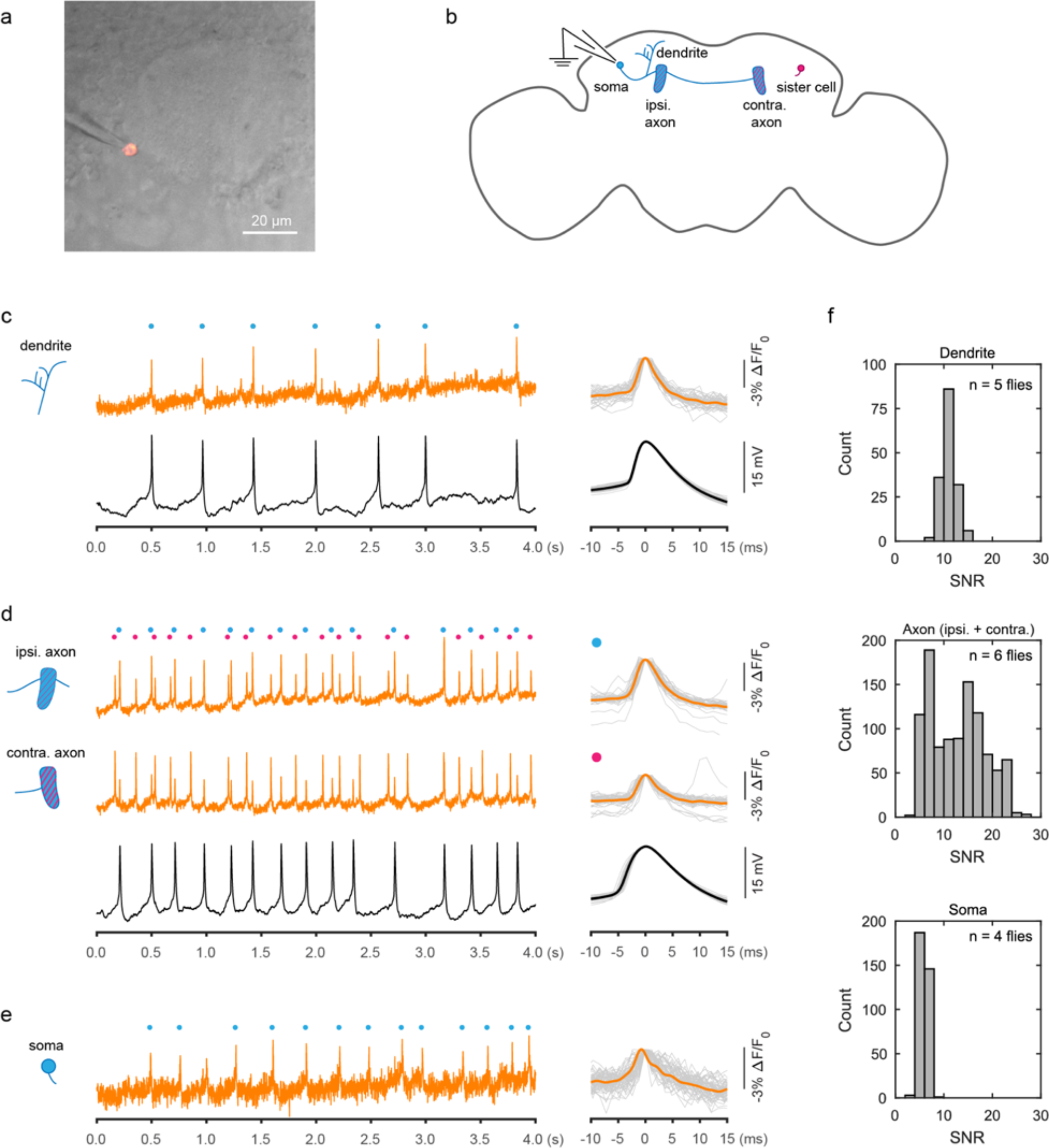
Recording spontaneous dopamine neuron activity in living adult flies using Voltron imaging and whole-cell patch clamp **(A)** Infrared image of fly brain overlaid with Voltron_549_ florescence in a dopamine neuron. Voltron expression was driven by a split-Gal4 driver (MB058B-Gal4) that labels a pair of PPL1-α’2/α2 dopamine neurons, one on each brain hemisphere. **(B)** PPL1-α’2/α2 neurons receive dendritic input from the ipsilateral hemisphere, and send axonal output bilaterally to form patch-like innervations in both the ipsilateral (ipsi.) and contralateral (contra.) mushroom body lobes. Consequently, each projection zone in the mushroom body contains axonal terminals of both cells, although each cell body contributes more extensive arbor to the projection zone in the same hemisphere. **(C-E)** Voltron imaging in different neuronal compartments. Left, single-trial recordings of fluorescence traces and cell membrane potential from concurrent wholecell recording. Circles mark action potential spikes detected in the Voltron traces. In dendrites, there was a perfect match between spikes detected from Voltron and from whole-cell recording. In axons, about half of spikes on the Voltron traces were contributed by the sister cell whose soma is in the opposite hemisphere. When these spikes were segregated (see methods), the remaining events aligned with whole-cell recording with marginal error (3 false positive from 447 Voltron spikes, 7 false negative from 451 wholecell recording spikes). Note that Voltron traces from the soma could not be imaged while recording in whole-cell mode. Right, spike waveforms aligned to their peaks. For axon, both Voltron spike waveforms were from the ipsilateral traces. **(F)** Signal-to-noise ratios (SNRs) in different neuronal compartments, calculated as Voltron spike peak amplitude over standard deviation of the spike-free zones of the trace.

**Supplementary Figure 8.**
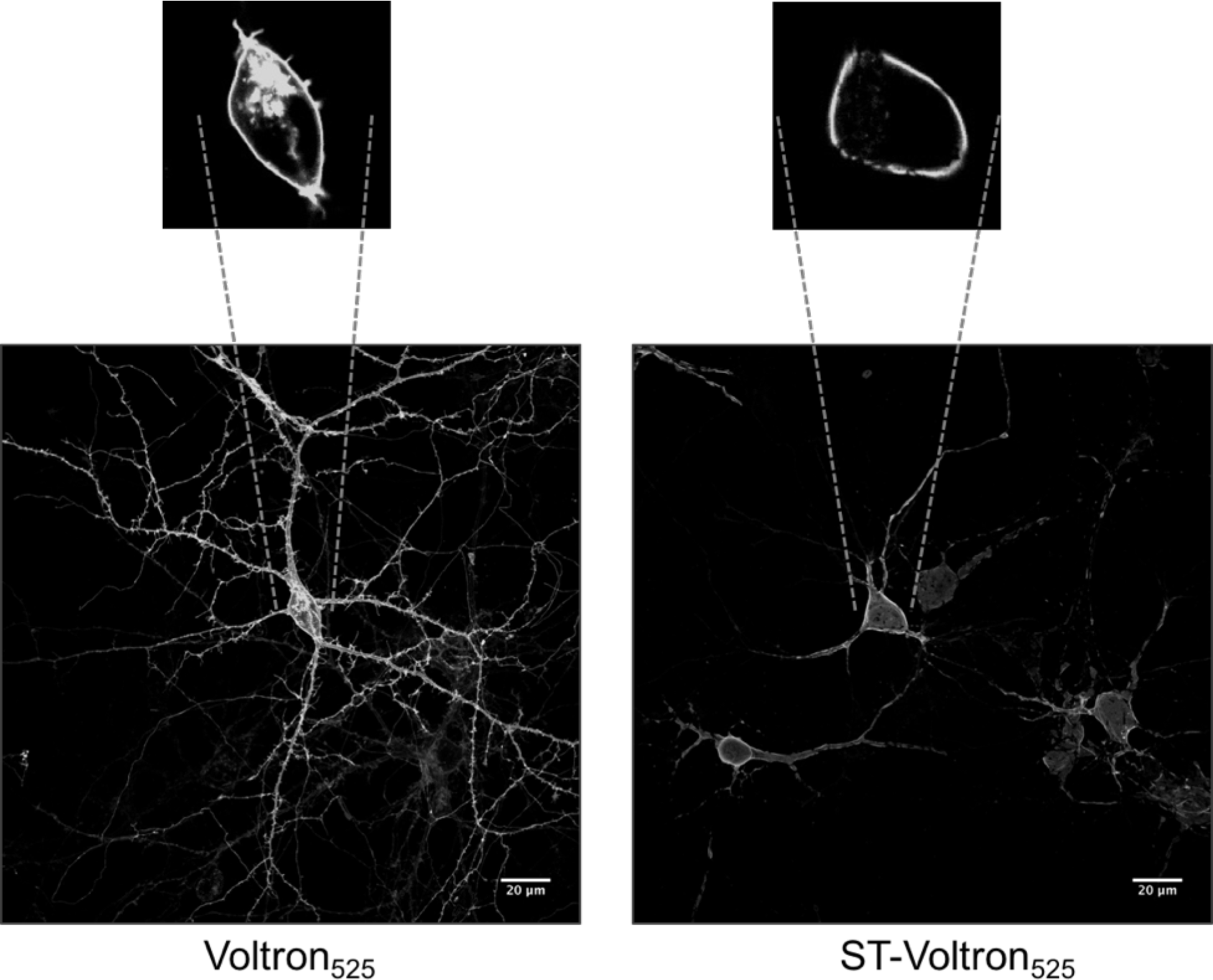
Bottom: Maximum intensity projections of confocal stacks of neurons in culture expressing Voltron (left) or soma-targeted Voltron (right) and labeled with JF_525_. Top: Zoom in on neuron soma showing cell membrane labeling and intracellular labelling, presumably endoplasmic reticulum. The soma localization tag limits labeling of processes and improves trafficking of Voltron to the cell membrane.

**Supplementary Figure 17.**
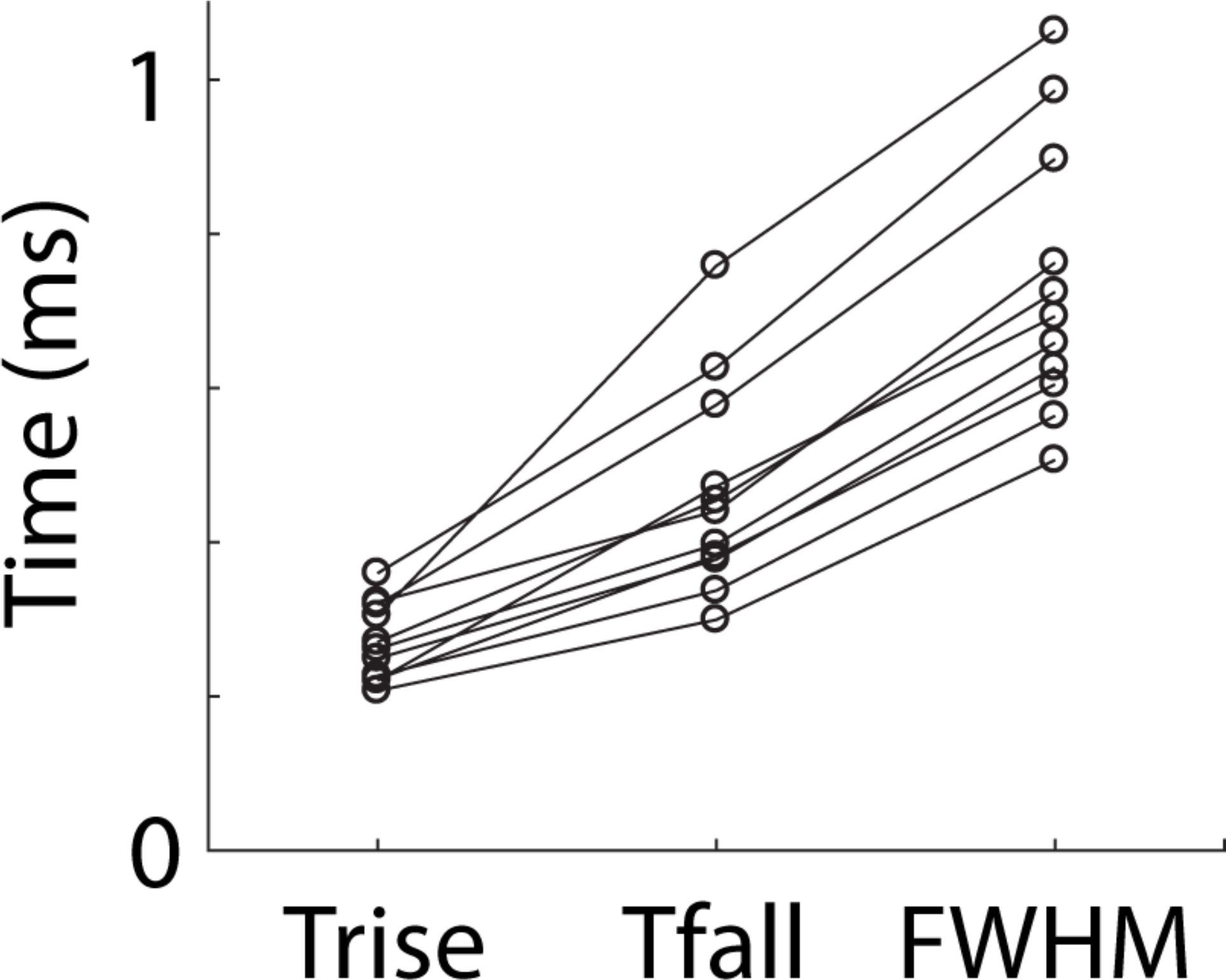
Analysis of membrane voltage dynamics in hippocampal parvalbumin (PV) neurons of awake mouse. The half rise time, half decay time, and full width half maximum of the spike waveforms shown in **Fig. 2G**.

**Supplementary Figure 18.**
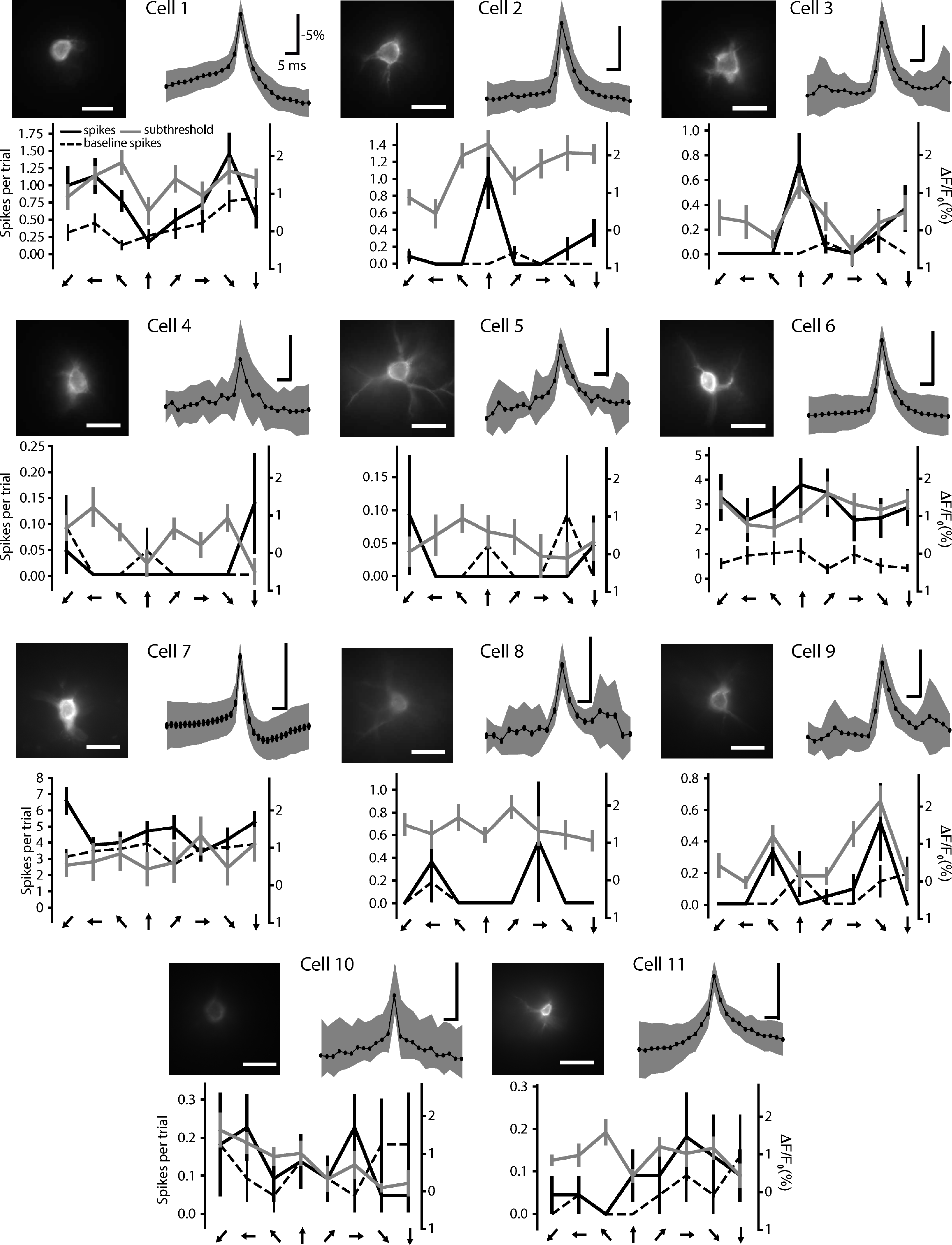
Orientation tuning of 11 pyramidal cells at depths of 100 - 250 μm in visual cortex of two C57B6 mice expressing Voltron_525_ under the control of CamKII-Cre. Each of the 11 cell panels includes fluorescence image of cell (top left, scalebar: 20 μM), average of all spikes in session (top right, scalebars: −5% ΔF/F, 5ms) and orientation tuning to full-frame drifting gratings of neurons (bottom), displayed from number of spikes during trials (solid black line), number of spikes during preceding inter-trial intervals (dashed black line), and subthreshold ΔF/F0 (right y-axis, solid gray line) after low-pass filtering traces using a 10-point median filter.

**Supplementary Figure 19.**
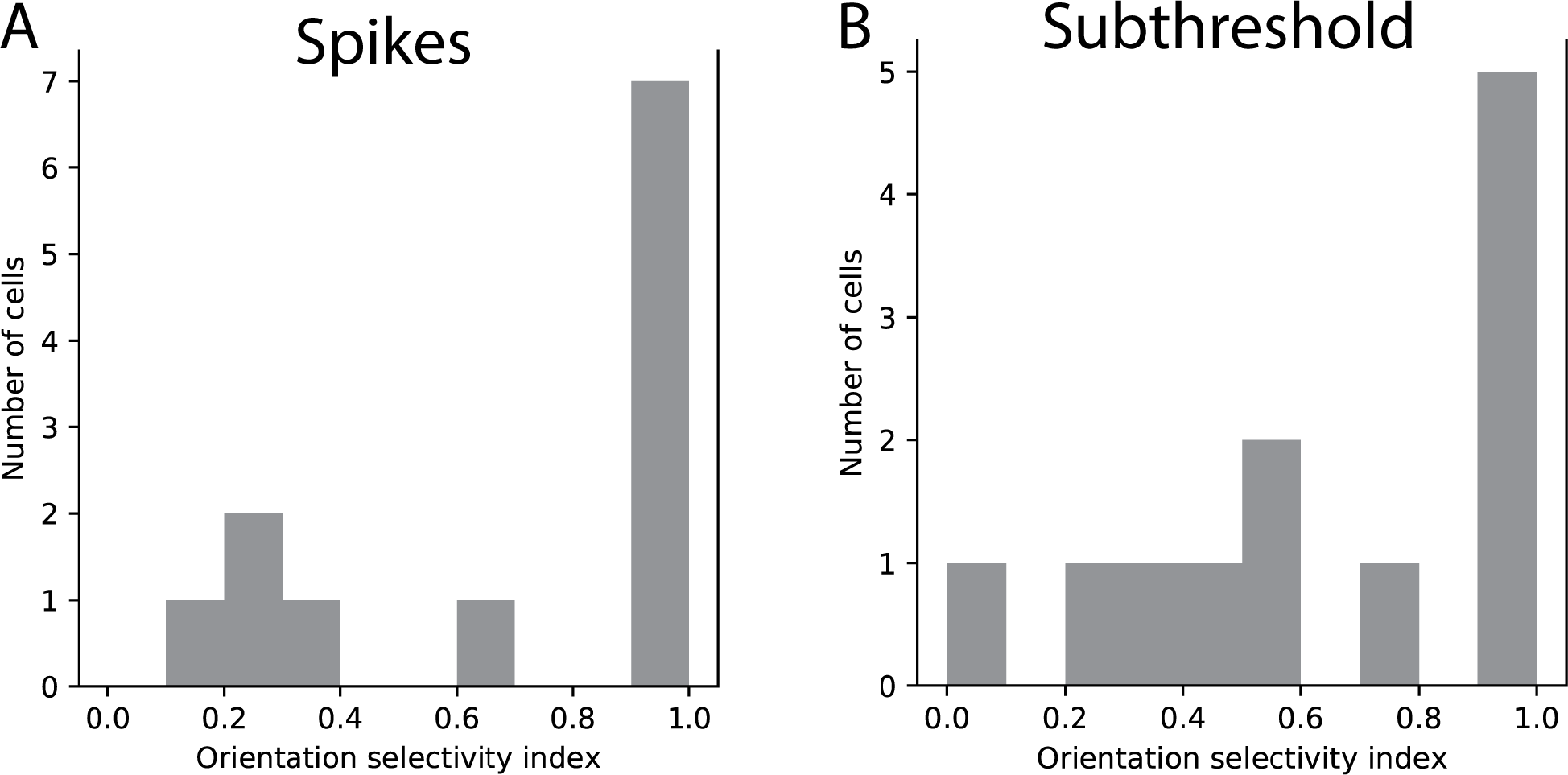
**(A)** Histogram of orientation selectivity index calculated from number of spikes in trial **(B)** Histogram of orientation selectivity index calculated from subthreshold membrane potential

**Supplementary Figure 20.**
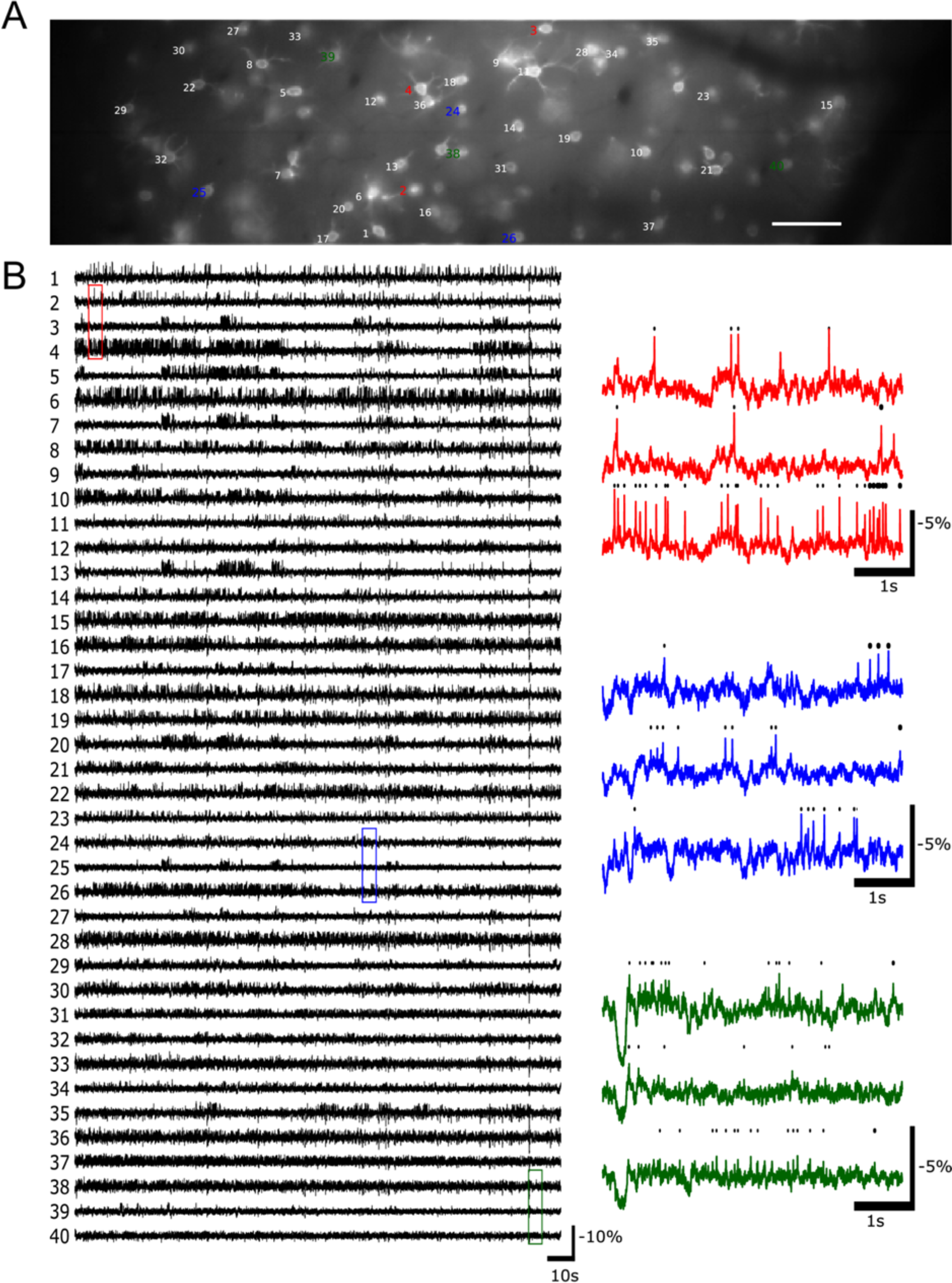
**(A)** Layer 1 interneurons expressing Voltron_525_ (same field of view as in Fig 3F). Scalebar: 100 μm. **(B)** Fluorescence traces from neurons labeled in (A), in decreasing order of signal to noise ratio. Signals processed as in Fig. 3G but without the last step of global background subtraction. **(C)** Zooms of fluorescence traces from color coded regions of with detected action potentials represented as black dots above, illustrating representative traces with high (top), medium (middle), and low (bottom) SNR.

**Supplementary Figure 21.**
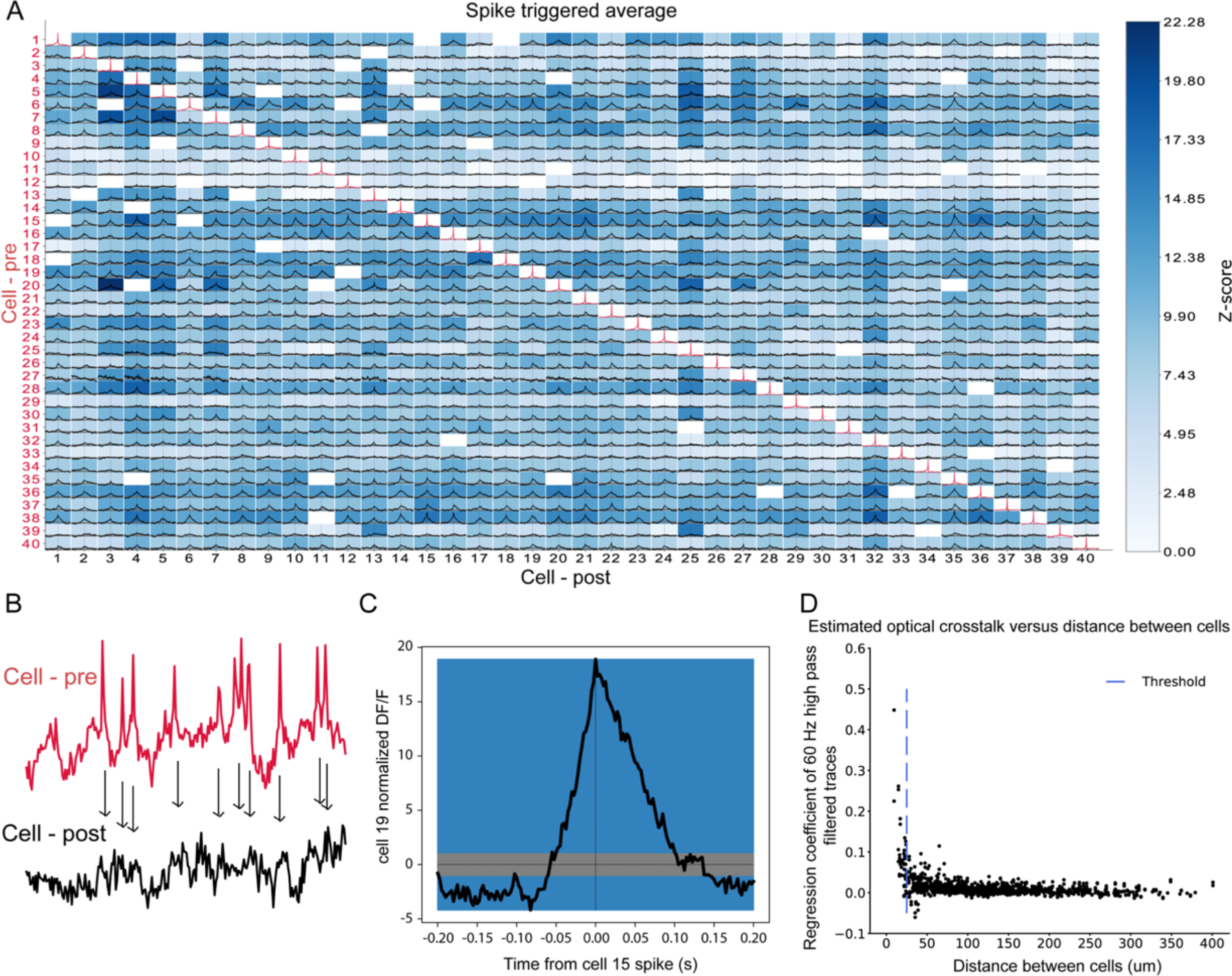
Spike triggered averages of neuron ensemble. **(A)** Spike triggered averages calculated from traces shown in Fig. S20. For each pair of neurons, estimated spike times of the first neuron (cell-pre, rows) were used to calculate the average membrane potential of the second neuron (cell-post, columns) in a window of 400 ms around the spike times. Diagonal line shown in red is the average spike shape of each neuron. For details, see supplementary methods. **(B)** Schematic to illustrate calculation of spike triggered average **(C)** Zoom-in of spike triggered average for cell 15 (pre) to cell 19 (post). Gray bar: standard deviation of shuffled spike triggered average (see supplementary methods). **(D)** Estimated optical cross talk between a pair of neurons. For neurons very close to each other, there was apparent optical cross-talk between the neurons which makes the spike triggered average calculated in this way unreliable. Blue line: distance threshold based on which pairs of neurons were excluded in (A). Excluded neuron pairs are shown as white squares in (A).

**Supplementary Figure 22.**
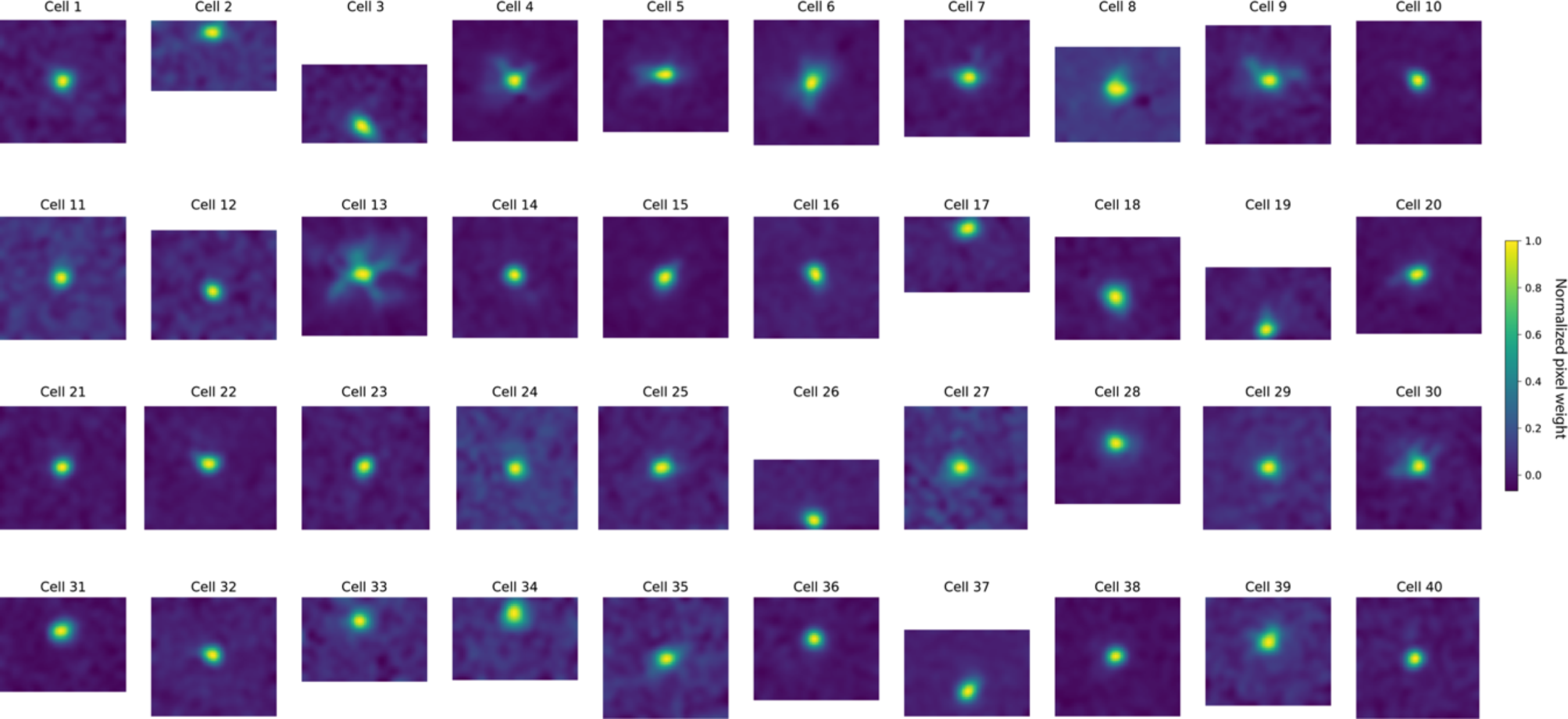
Spatial filters for context regions of 50×50 pixels centered on the ROI of each cell shown in Fig. 3F-G and Fig S20 (see supplementary methods for details). Spatial filters were estimated by Spike Pursuit (supplementary methods). Cells near the boundary of the field of view have different sizes of context region.

**Supplementary Figure 23.**
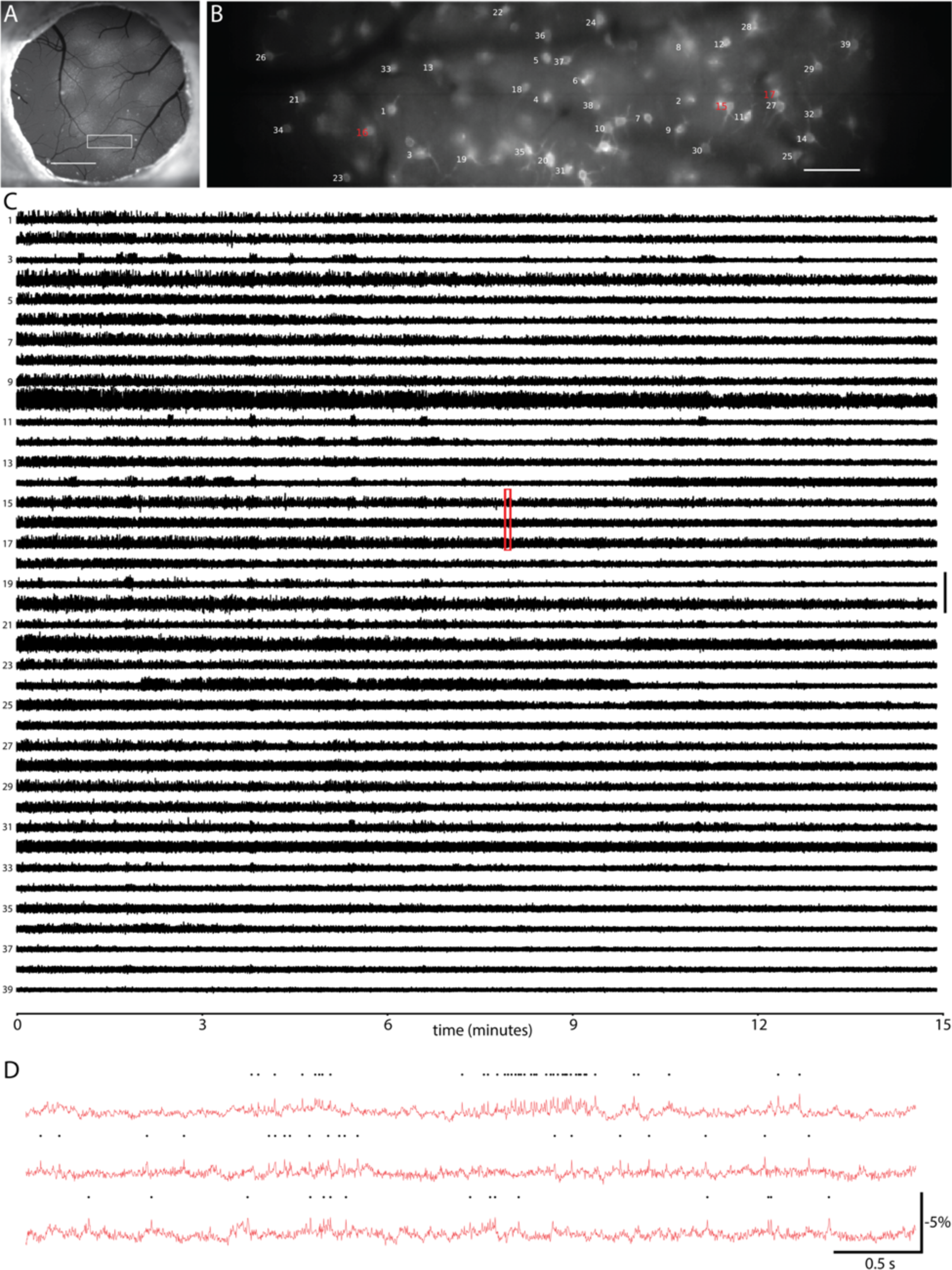
**(A)** Fluorescence image of a cranial window over primary visual cortex (V1) in an NDNF-Cre mouse showing Cre-dependent expression of soma targeted Voltron_525_. Scalebar, 1 mm. **(B)** Fluorescence image of area indicated by the white rectangle in (A), with neuron labels corresponding to fluorescence traces in (C). Scalebar, 100 μm. **(C)** Fluorescence traces during 10-15 minutes recordings from neurons indicated in (B), in decreasing order of signal to noise ratio. Signals processed as described in Supplementary Methods. (D) Zoom-in of fluorescence traces from area indicated by red rectangle in (C).

**Supplementary Figure 24.**
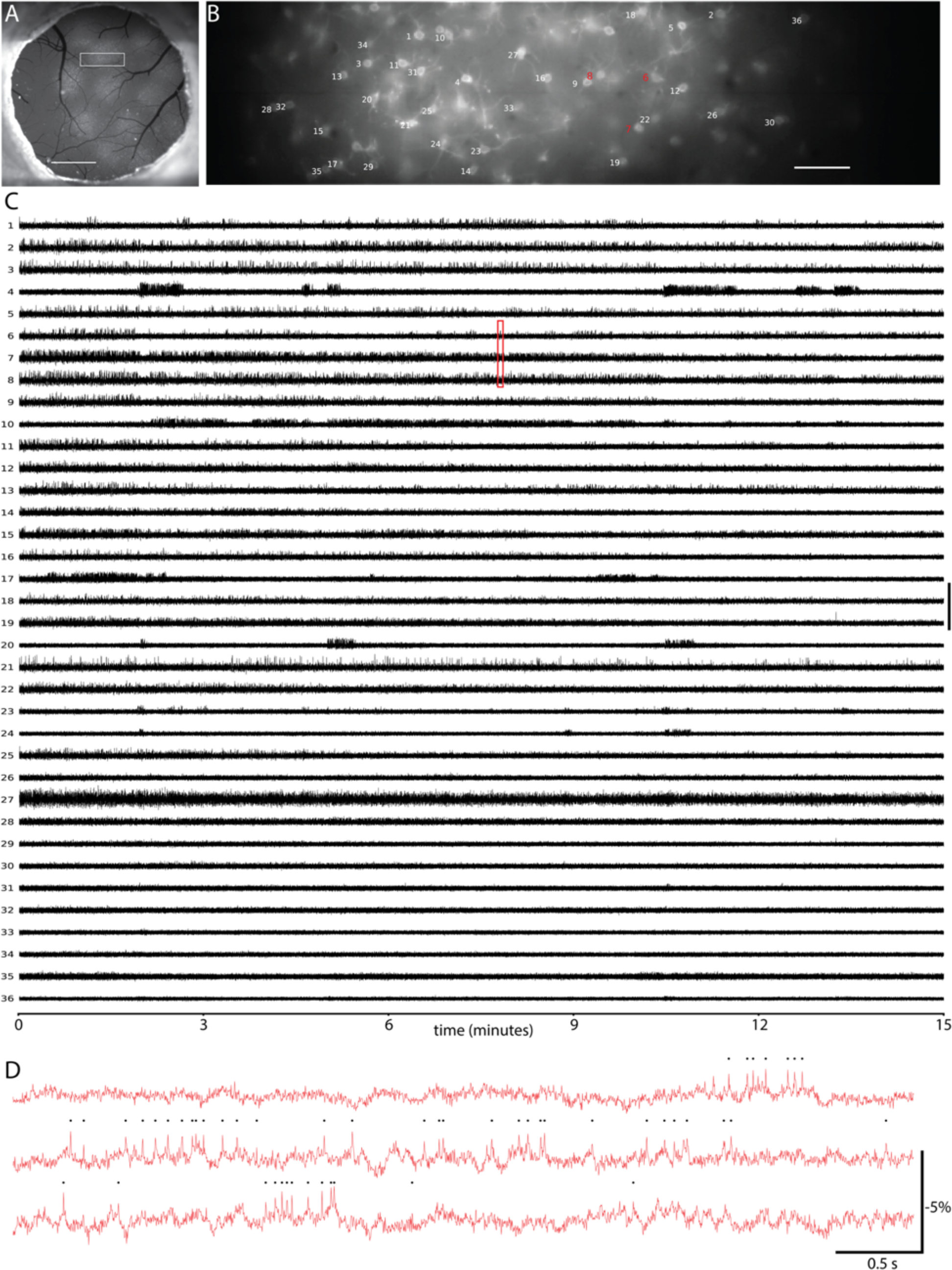
**(A)** Fluorescence image of a cranial window over primary visual cortex (V1) in an NDNF-Cre mouse showing Cre-dependent expression of soma targeted Voltron_525_. Scalebar, 1 mm. **(B)** Fluorescence image of area indicated by the white rectangle in (A), with neuron labels corresponding to fluorescence traces in (C). Scalebar, 100 μm. **(C)** Fluorescence traces during 10-15 minutes recordings from neurons indicated in (B), in decreasing order of signal to noise ratio. Signals processed as described in Supplementary Methods. (D) Zoom-in of fluorescence traces from area indicated by red rectangle in (C).

**Supplementary Figure 25.**
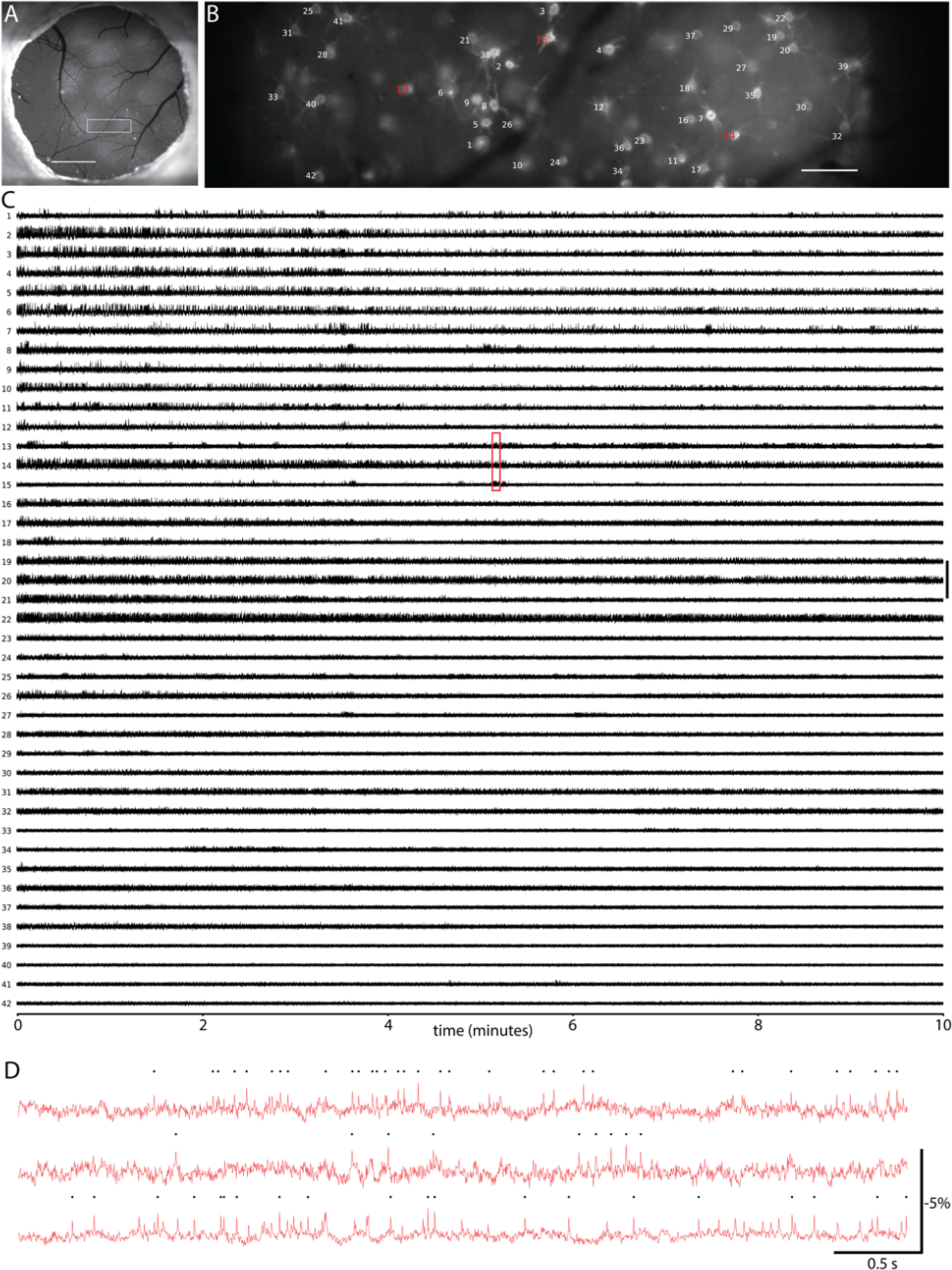
**(A)** Fluorescence image of a cranial window over primary visual cortex (V1) in an NDNF-Cre mouse showing Cre-dependent expression of soma targeted Voltron_525_. Scalebar, 1 mm. **(B)** Fluorescence image of area indicated by the white rectangle in (A), with neuron labels corresponding to fluorescence traces in (C). Scalebar, 100 μm. **(C)** Fluorescence traces during 10-15 minutes recordings from neurons indicated in (B), in decreasing order of signal to noise ratio. Signals processed as described in Supplementary Methods. (D) Zoom-in of fluorescence traces from area indicated by red rectangle in (C).

**Supplementary Figure 26.**
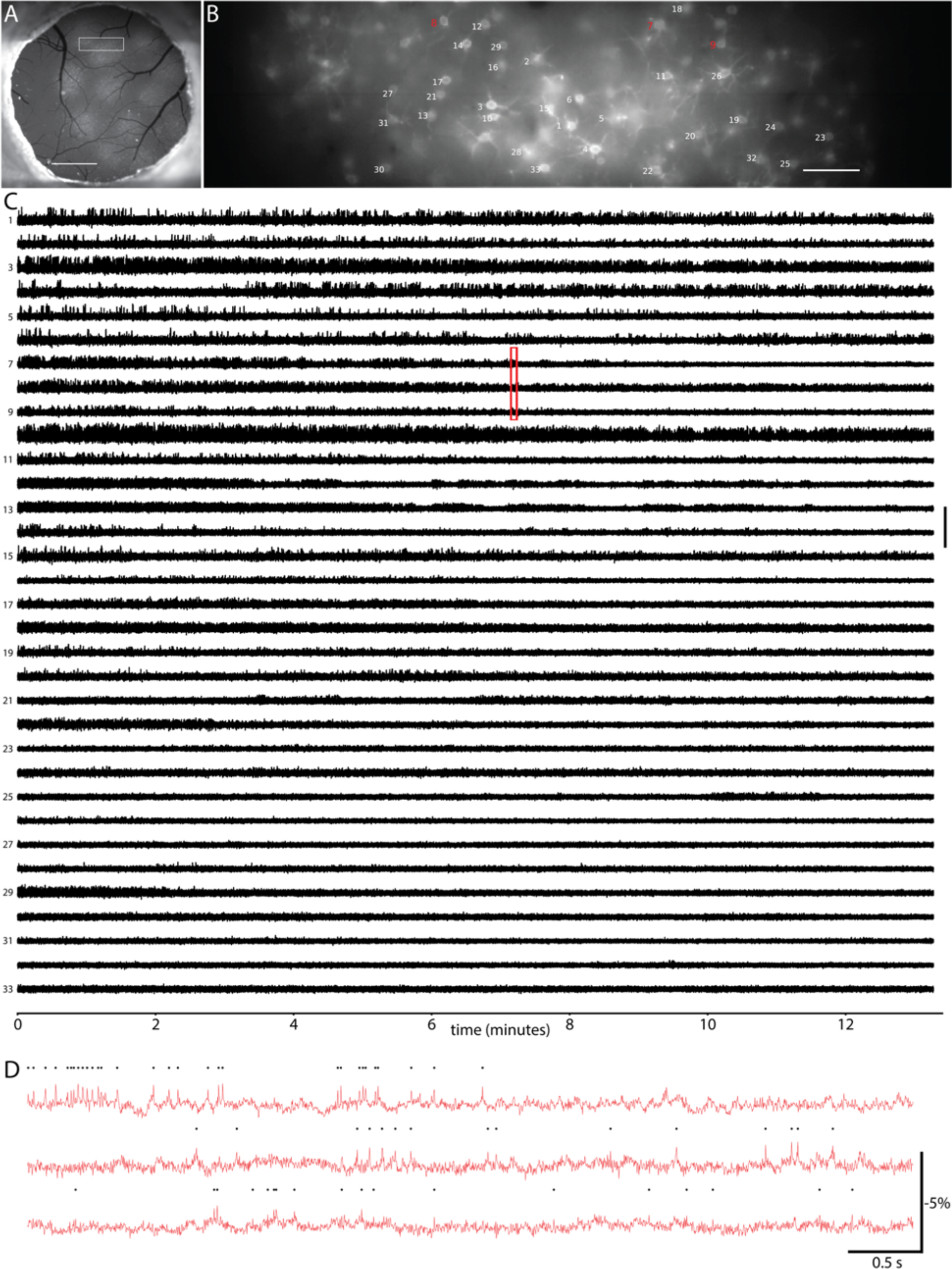
**(A)** Fluorescence image of a cranial window over primary visual cortex (V1) in an NDNF-Cre mouse showing Cre-dependent expression of soma targeted Voltron_525_. Scalebar, 1 mm. **(B)** Fluorescence image of area indicated by the white rectangle in (A), with neuron labels corresponding to fluorescence traces in (C). Scalebar, 100 μm. **(C)** Fluorescence traces during 10-15 minutes recordings from neurons indicated in (B), in decreasing order of signal to noise ratio. Signals processed as described in Supplementary Methods. (D) Zoom-in of fluorescence traces from area indicated by red rectangle in (C).

**Supplementary Figure 27.**
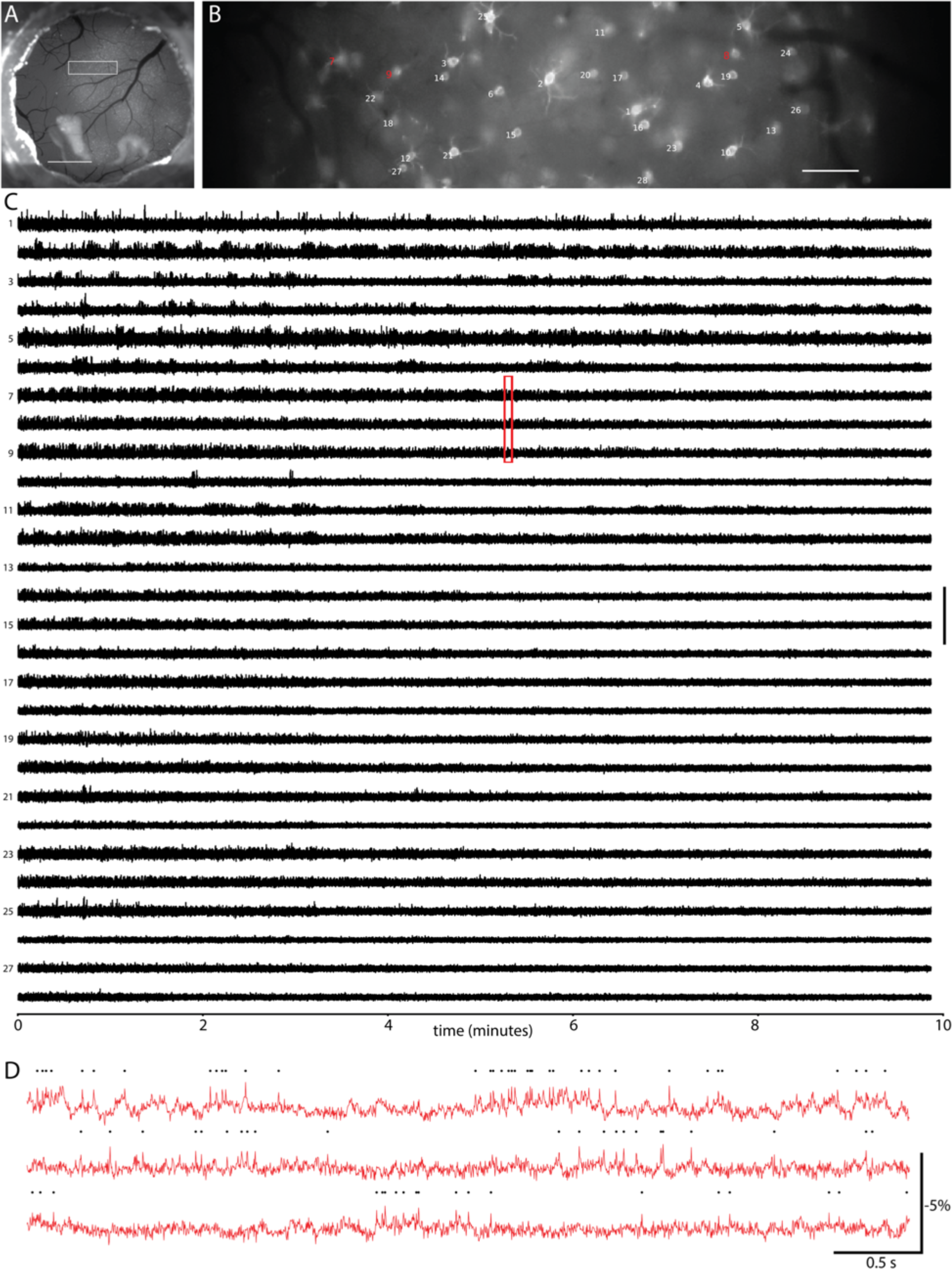
**(A)** Fluorescence image of a cranial window over primary visual cortex (V1) in an NDNF-Cre mouse showing Cre-dependent expression of soma targeted Voltron_525_. Scalebar, 1 mm. **(B)** Fluorescence image of area indicated by the white rectangle in (A), with neuron labels corresponding to fluorescence traces in (C). Scalebar, 100 μm. **(C)** Fluorescence traces during 10-15 minutes recordings from neurons indicated in (B), in decreasing order of signal to noise ratio. Signals processed as described in Supplementary Methods. (D) Zoom-in of fluorescence traces from area indicated by red rectangle in (C).

**Supplementary Figure 28.**
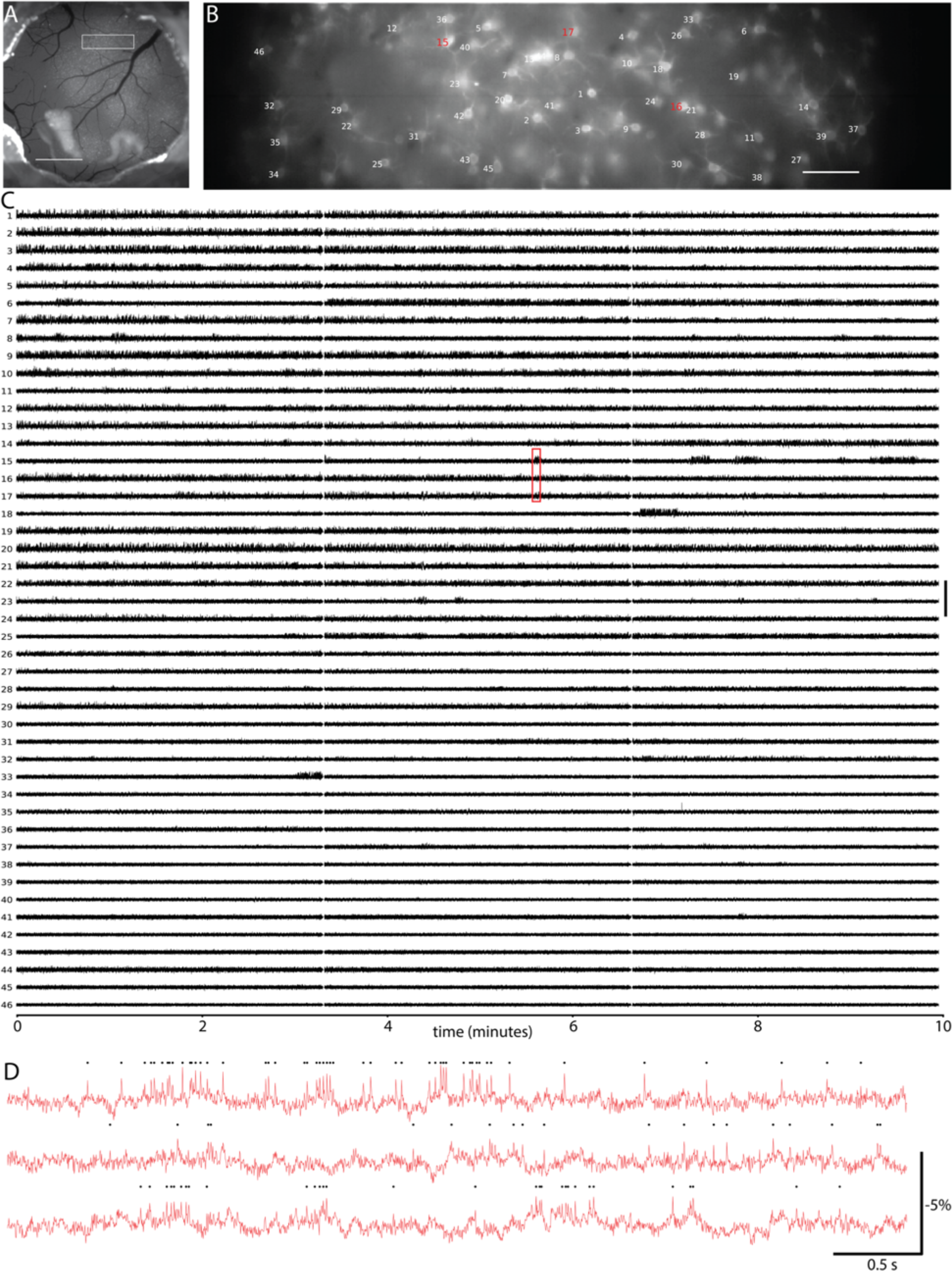
**(A)** Fluorescence image of a cranial window over primary visual cortex (V1) in an NDNF-Cre mouse showing Cre-dependent expression of soma targeted Voltron_525_. Scalebar, 1 mm. **(B)** Fluorescence image of area indicated by the white rectangle in (A), with neuron labels corresponding to fluorescence traces in (C). Scalebar, 100 μm. **(C)** Fluorescence traces during 10-15 minutes recordings from neurons indicated in (B), in decreasing order of signal to noise ratio. Signals processed as described in Supplementary Methods. (D) Zoom-in of fluorescence traces from area indicated by red rectangle in (C).

**Supplementary Figure 29.**
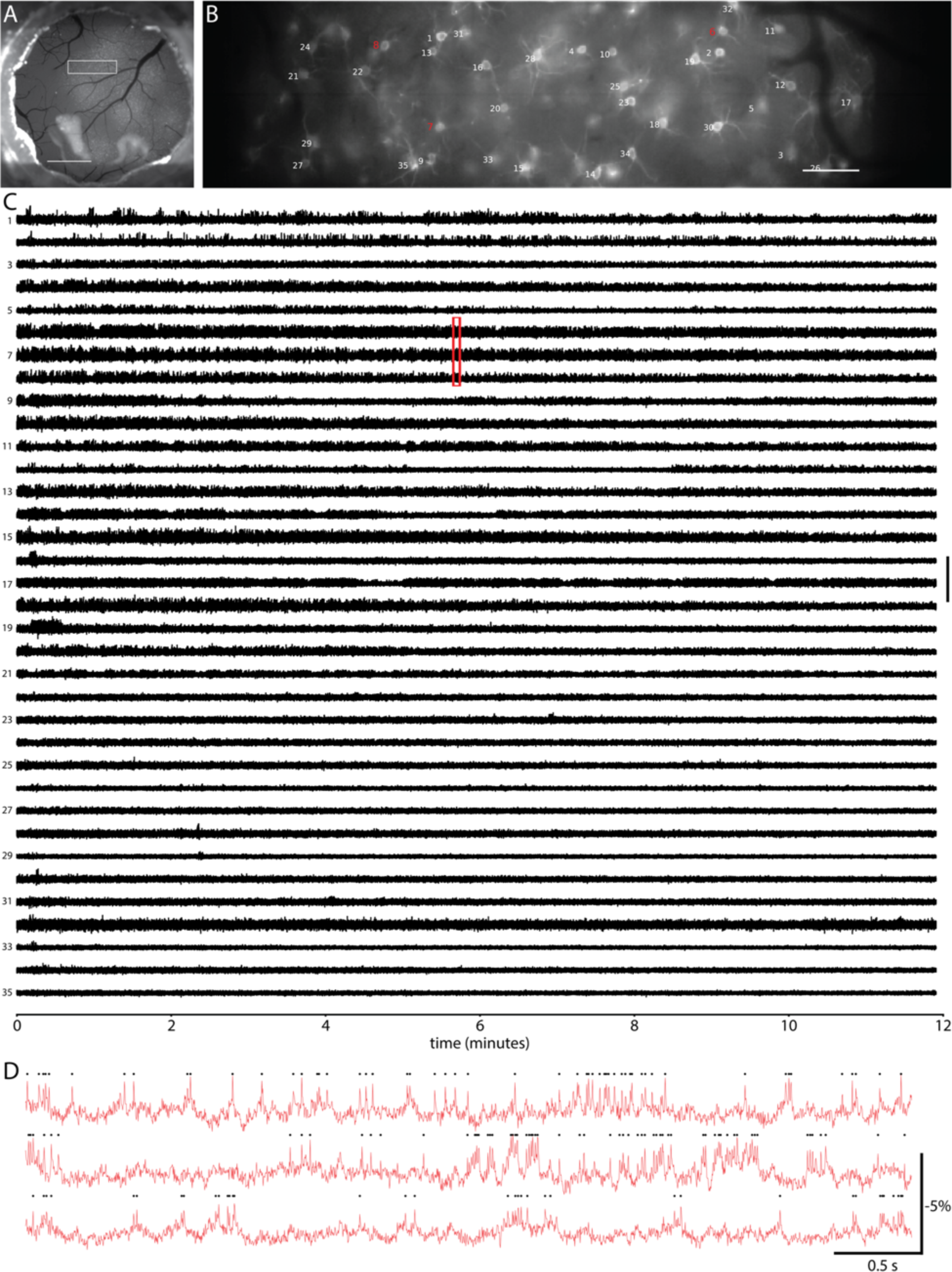
**(A)** Fluorescence image of a cranial window over primary visual cortex (V1) in an NDNF-Cre mouse showing Cre-dependent expression of soma targeted Voltron_525_. Scalebar, 1 mm. **(B)** Fluorescence image of area indicated by the white rectangle in (A), with neuron labels corresponding to fluorescence traces in (C). Scalebar, 100 μm. **(C)** Fluorescence traces during 10-15 minutes recordings from neurons indicated in (B), in decreasing order of signal to noise ratio. Signals processed as described in Supplementary Methods. (D) Zoom-in of fluorescence traces from area indicated by red rectangle in (C).

**Supplementary Figure 30.**
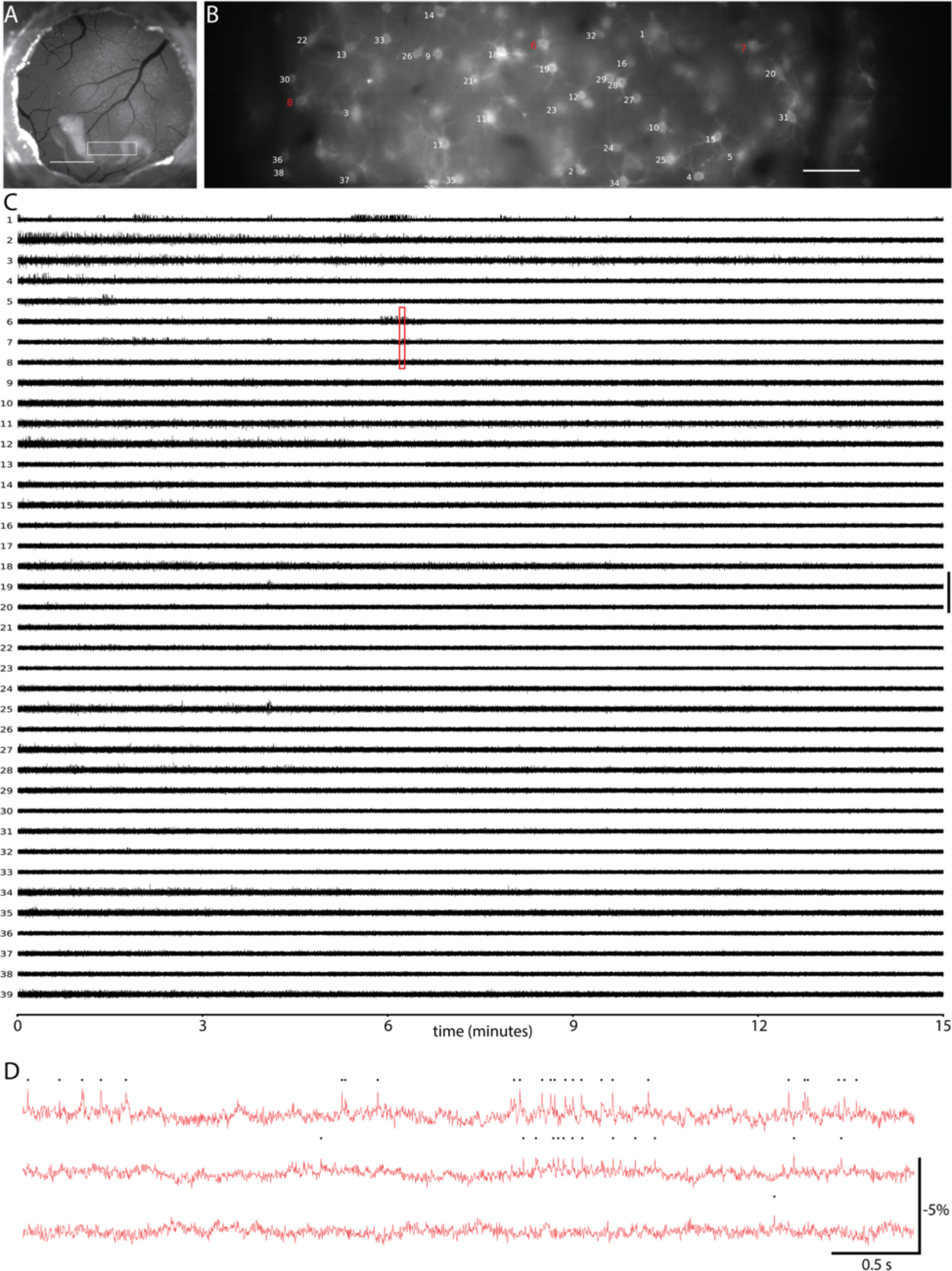
**(A)** Fluorescence image of a cranial window over primary visual cortex (V1) in an NDNF-Cre mouse showing Cre-dependent expression of soma targeted Voltron_525_. Scalebar, 1 mm. **(B)** Fluorescence image of area indicated by the white rectangle in (A), with neuron labels corresponding to fluorescence traces in (C). Scalebar, 100 μm. **(C)** Fluorescence traces during 10-15 minutes recordings from neurons indicated in (B), in decreasing order of signal to noise ratio. Signals processed as described in Supplementary Methods. (D) Zoom-in of fluorescence traces from area indicated by red rectangle in (C).

**Supplementary Figure 31.**
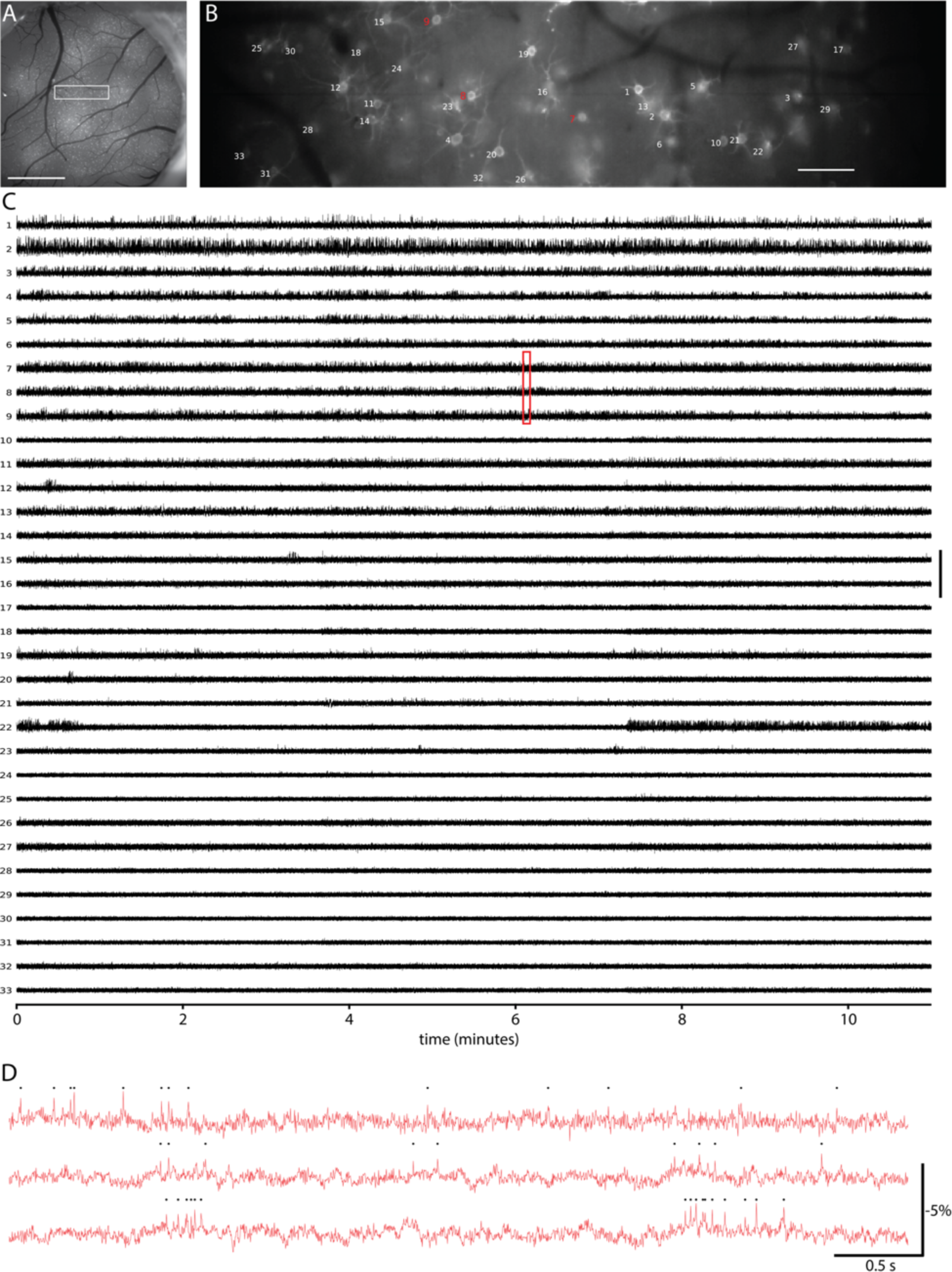
**(A)** Fluorescence image of a cranial window over primary visual cortex (V1) in an NDNF-Cre mouse showing Cre-dependent expression of soma targeted Voltron_525_. Scalebar, 1 mm. **(B)** Fluorescence image of area indicated by the white rectangle in (A), with neuron labels corresponding to fluorescence traces in (C). Scalebar, 100 μm. **(C)** Fluorescence traces during 10-15 minutes recordings from neurons indicated in (B), in decreasing order of signal to noise ratio. Signals processed as described in Supplementary Methods. (D) Zoom-in of fluorescence traces from area indicated by red rectangle in (C).

**Supplementary Figure 32.**
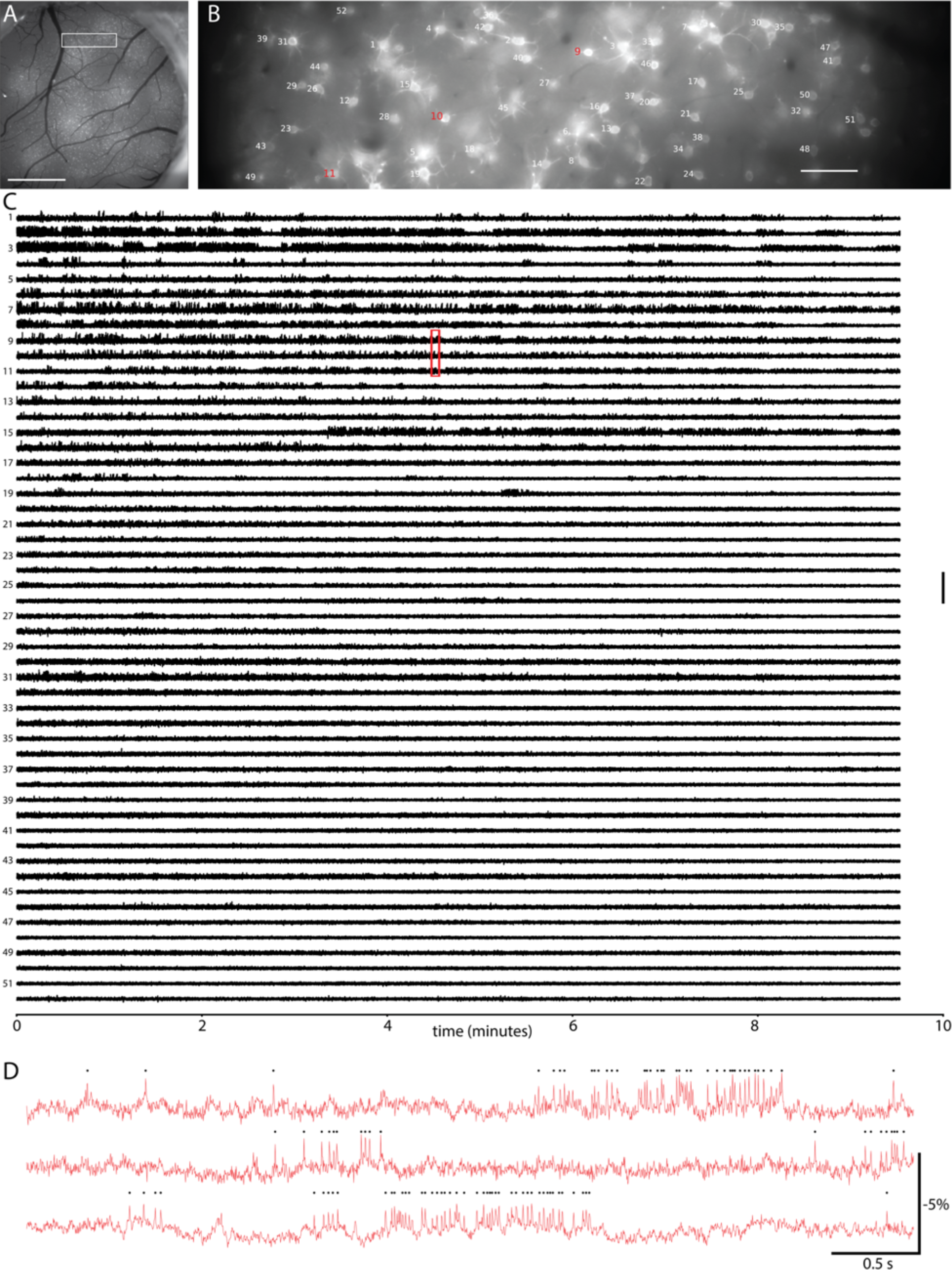
**(A)** Fluorescence image of a cranial window over primary visual cortex (V1) in an NDNF-Cre mouse showing Cre-dependent expression of soma targeted Voltron_525_. Scalebar, 1 mm. **(B)** Fluorescence image of area indicated by the white rectangle in (A), with neuron labels corresponding to fluorescence traces in (C). Scalebar, 100 μm. **(C)** Fluorescence traces during 10-15 minutes recordings from neurons indicated in (B), in decreasing order of signal to noise ratio. Signals processed as described in Supplementary Methods. (D) Zoom-in of fluorescence traces from area indicated by red rectangle in (C).

**Supplementary Figure 33.**
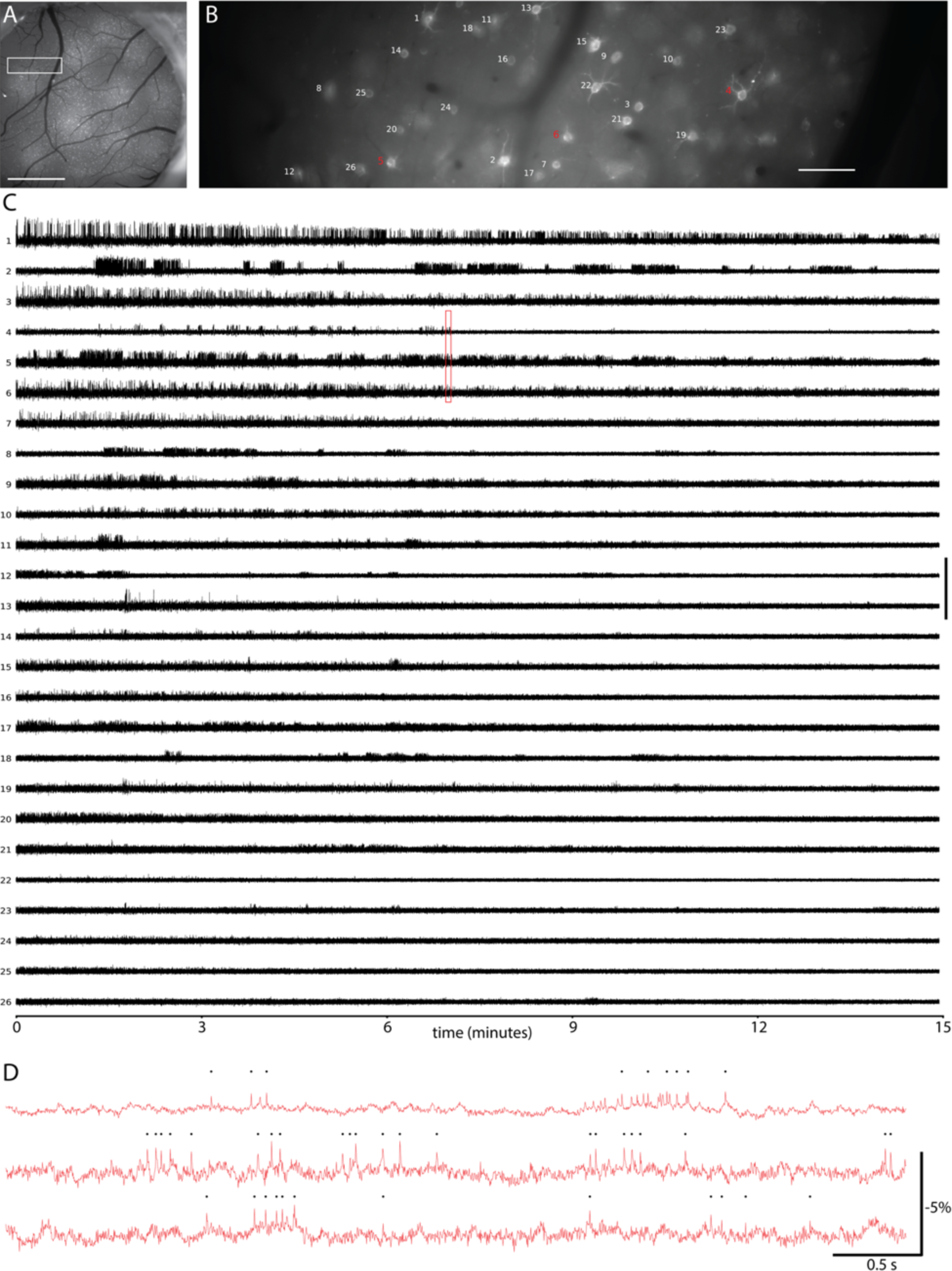
**(A)** Fluorescence image of a cranial window over primary visual cortex (V1) in an NDNF-Cre mouse showing Cre-dependent expression of soma targeted Voltron_525_. Scalebar, 1 mm. **(B)** Fluorescence image of area indicated by the white rectangle in (A), with neuron labels corresponding to fluorescence traces in (C). Scalebar, 100 μm. **(C)** Fluorescence traces during 10-15 minutes recordings from neurons indicated in (B), in decreasing order of signal to noise ratio. Signals processed as described in Supplementary Methods. (D) Zoom-in of fluorescence traces from area indicated by red rectangle in (C).

**Supplementary Figure 34.**
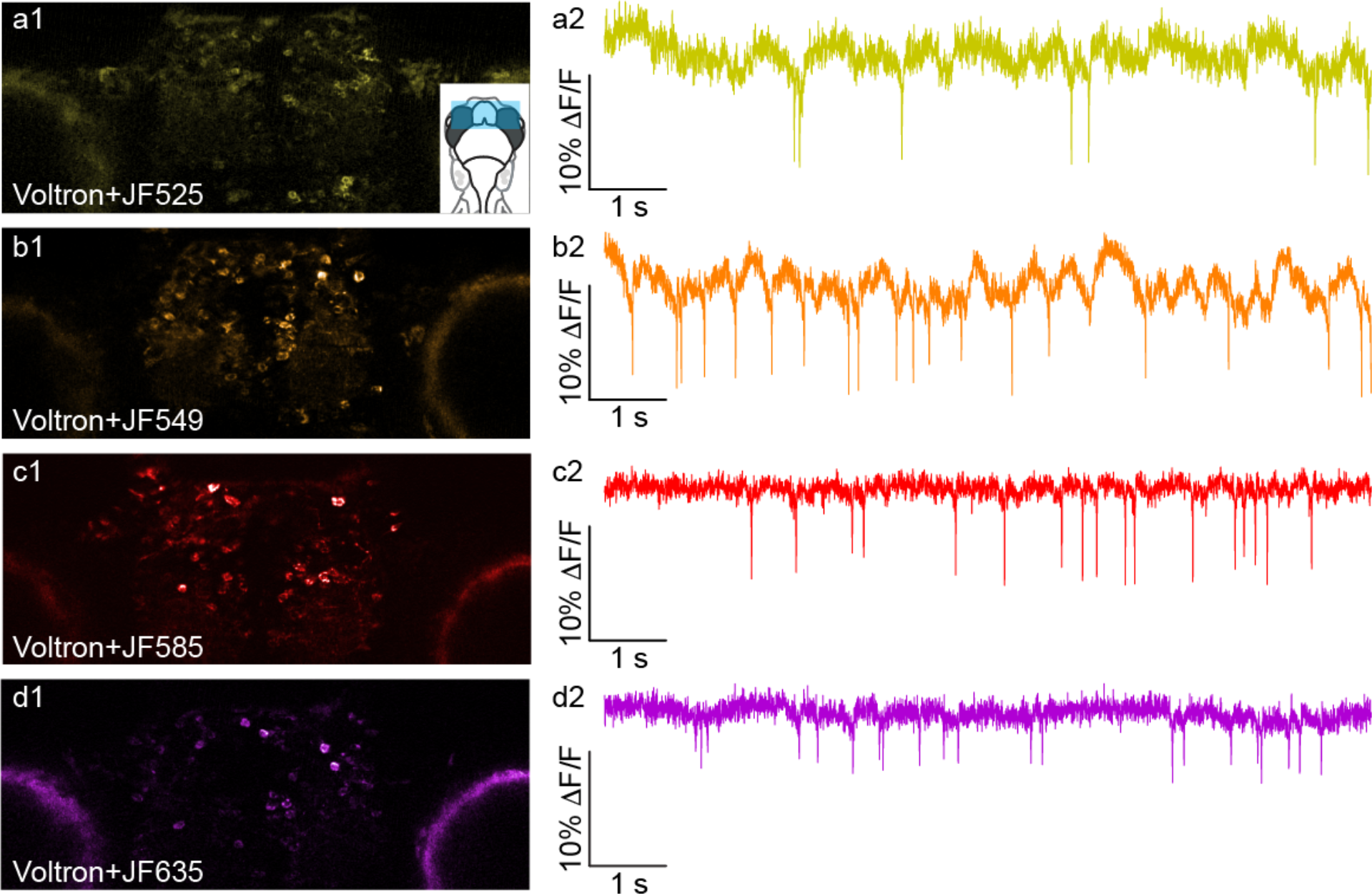
Multi-spectrum voltage imaging in zebrafish **(A1)** Forebrain neurons expressing Voltron-ST labeled with Janelia Fluor 525 (JF525). (Inset) A schematic drawing showing the location of the image. **(A2)** Fluorescence signal from a neuron labeled with Voltron-ST+JF525 showing spontaneous spiking activity. **(B1)** Forebrain neurons expressing Voltron-ST labeled with Janelia Flour 549 (JF549). **(B2)** Fluorescence signal from a neuron labeled with Voltron-ST+JF549 showing spontaneous spiking activity. **(C1)** Forebrain neurons expressing Voltron-ST labeled with Janelia Fluor 585 (JF585). **(C2)** Fluorescence signal from a neuron labeled with Voltron-ST+JF585 showing spontaneous spiking activity. (**D1)** Forebrain neurons expressing Voltron-ST labeled with Janelia Fluor 635 (JF635). **(D2)** Fluorescence signal from a neuron expressing Voltron-ST+JF635 showing spontaneous spiking activity.

**Supplementary Figure 35.**
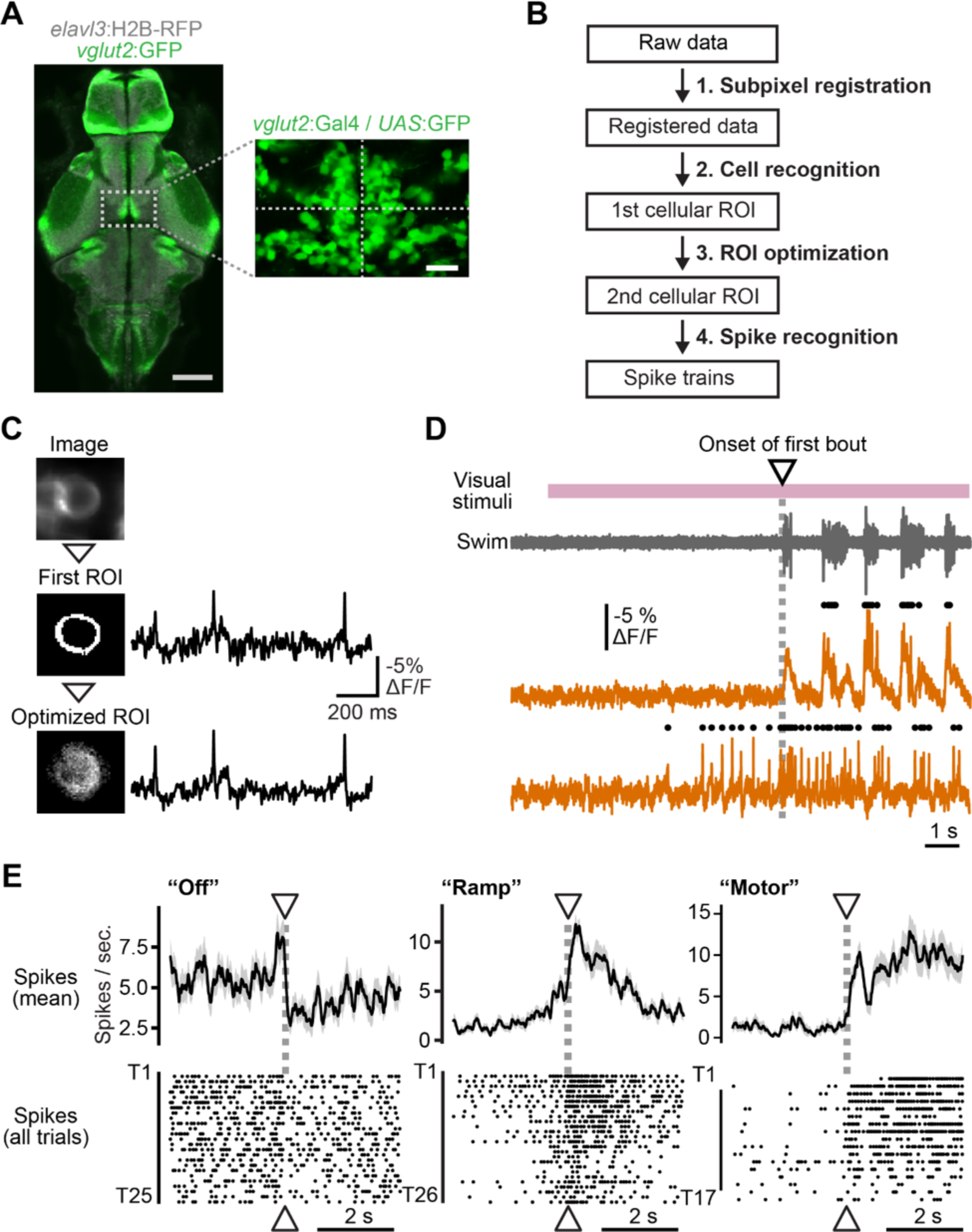
Recording and analyzing Voltron data in behaving zebrafish. **(A)** *Left*, anatomical location of the midbrain nucleus imaged in this study. The image was taken from plane 81 of image stacks from *Tg(eval3:H2B-RFP*) (gray) and *Tg(vglut2:GFP*) (green) transgenic zebrafish in the Z-brain atlas(*33*). Right, a representative image of the same nucleus in a *Tg(vglut2:Gal4); Tg(UAS:GFP*) transgenic zebrafish. **(B)** Flow chart of the data processing pipeline for the acquired data. **(C)** An example of pixel weight optimization for a representative neuron. Traces from the initial pixel weights (middle) and the final pixel weights (bottom) of the same neurons are plotted on the right. **(D)** Schematic of averaging procedure of neural activity at the onset of first bout of the swimming for each trial for the analysis in (E). **(E)** Average firing rates (top) and raster plots across trials (bottom) at the onset of the first bout for each trial plotted for 3 representative neurons on a long timescale (−3 seconds to 3 seconds). The example ‘Off’ neuron shows suppression of firing at the onset of swimming (following a brief increase in firing rate just before swimming). The ‘Ramp’ neuron shows a gradual increase in activity starting about 1.5 seconds before the onset of swimming, and a decay in activity after swim onset. ‘Motor’-type neurons (subdivided into ‘Onset’ and ‘Late’ neurons in Fig. 4) show increased firing at the onset of swimming. Shadows represent standard error of the mean (s.e.m.) across multiple trials.

**Supplementary Table1.** Voltron_525_ and Ace2N-mNeon kinetics in primary neuron culture cells

|  | Activation<br>(-70 mV to 30 mV) |  |  | Deactivation<br>(30 mV to -70 mV) |  |  |
| --- | --- | --- | --- | --- | --- | --- |
| | $\tau_{\text{fast}}$ (ms) | $\tau_{\text{slow}}$ (ms) | % fast | $\tau_{\text{fast}}$ (ms) | $\tau_{\text{slow}}$ (ms) | % fast |
| Ace2N-mNeon | $0.59 \pm 0.10$ | $6.9 \pm 0.5$ | $62 \pm 3$ | $0.63 \pm 0.09$ | $6.0 \pm 1.1$ | $55 \pm 4$ |
| Voltron-JF525 | $0.64 \pm 0.09$ | $4.1 \pm 0.6$ | $61 \pm 4$ | $0.78 \pm 0.12$ | $3.9 \pm 0.2$ | $55 \pm 7$ |

Neurons expressing Ace2N-mNeon and Voltron<sub>525</sub> were imaged at 3.2 kHz during whole cell voltage clamp as detailed in the methods section. Fluorescence traces were then fit using a double exponential model (fig. S4). Errors are s.e.m. n = 7 cells for Ace-mNeon and n = 8 cells for Voltron<sub>525</sub>.

**Supplementary Table 2.** Photophysical properties of parent fluorophore HaloTag-bound JFdyes and green fluorescent proteins (mean values, n = 9)

| fluorophore | $\lambda_{\text{abs}}$<br>(nm) | $\lambda_{\text{em}}$<br>(nm) | $\epsilon_{\text{max}}$<br>($\text{mM}^{-1}\text{cm}^{-1}$) | $\Phi_{\text{fl}}$ | $W^{(a)}$<br>($\text{sec}^{-1}$) | $\tau_1(A_1)^{(b)}$<br>(sec) | $\tau_2(A_2)$<br>(sec) | $\langle \tau_b \rangle^{(c)}$<br>(sec) | $P_b / 10^{-6}^{(d)}$ | $N_{\text{ph}} / 10^5^{(e)}$ | $\tau_{1/2}^{(f)}$<br>(sec) |
| --- | --- | --- | --- | --- | --- | --- | --- | --- | --- | --- | --- |
| JF505 | 509 | 531 | 56.8 | 0.83 | 941 | 59 (0.59) | 264 (0.41) | 144 | 7.36 | 1.13 | 51 |
| JF525 | 532 | 553 | 83.6 | 0.87 | 1,549 | 329 (1.0) |  | 330 | 1.96 | 4.43 | 336 |
| JF549 | 555 | 578 | 91.3 | 0.86 | 3,158 | 907 (0.65) | 124 (0.35) | 633 | 0.50 | 17.2 | 804 |
| JF585 | 593 | 611 | 109 | 0.83 | 4,443 | 931 (1.0) |  | 950 | 0.24 | 35.1 | 3,320 |
| JF635 | 641 | 656.5 | 64.0 | 0.7 | 2,486 | 1976 (1.0) |  | 1,980 | 0.20 | 34.4 | 3,130 |
| sfGFP | 488 | 511 | 51.5 | 0.66 | 1,597 | 90 (0.66) | 167 (0.34) | 117 | 5.32 | 1.24 | 71 |
| mNeon Green | 506 | 517.5 | 120 | 0.85 | 1,773 | 102 (0.72) | 144 (0.28) | 114 | 4.83 | 1.80 | 118 |

<sup>(a)</sup> Excitation rate (absorbed photons/sec) equal to the integral over wavelength of the product of extinction coefficient and spectral irradiance.
<sup>(b)</sup> Decay constant (normalized amplitude) of multi-exponential fit to the photobleaching decay curve.
<sup>(c)</sup> Amplitude-weighted lifetime equal to $A_1\tau_1 + A_2\tau_2$ .
<sup>(d)</sup> Calculated probability of photobleaching per absorbed photon.
<sup>(e)</sup> Calculated number of photons emitted per molecule before photobleaching.
<sup>(f)</sup> Calculated time to bleach from a rate of 1000 to 500 photons per sec per molecule.

**Supplementary Table 3.** Photobleaching properties of GEVI sensors in neuronal cell culture (mean values, n = 5)

| sensor | $W^{(a)}$<br>(sec<sup>-1</sup>) | $\tau_1(A_1)^{(b)}$<br>(sec) | $\tau_2(A_2)$<br>(sec) | $\langle \tau_b \rangle^{(c)}$<br>(sec) | $P_b / 10^{-6}^{(d)}$ | $N_{ph} / 10^5^{(e)}$ | $\tau_{1/2}^{(f)}$<br>(sec) |
| --- | --- | --- | --- | --- | --- | --- | --- |
| Voltron525 | 1,549 | 171 (0.84) | 1497 (0.15) | 373 | 1.74 | 5.03 | 206 |
| Voltron549 | 3,158 | 675 (0.67) | 61.5 (0.31) | 471 | 0.672 | 12.8 | 538 |
| ASAP-2f | 1,597 | 400 (0.68) | 24.9 (0.30) | 281 | 2.23 | 2.96 | 132 |
| Ace2N-mNeon | 1,773 | 118 (0.88) | 147 (0.11) | 120 | 4.70 | 1.81 | 119 |

<sup>(a)</sup> Excitation rate (absorbed photons/sec) equal to the integral over wavelength of the product of extinction coefficient and spectral irradiance.
<sup>(b)</sup> Decay constant (normalized amplitude) of multi-exponential fit to the photobleaching decay curve.
<sup>(c)</sup> Amplitude-weighted lifetime equal to $A_1\tau_1 + A_2\tau_2$ .
<sup>(d)</sup> Calculated probability of photobleaching per absorbed photon.
<sup>(e)</sup> Calculated number of photons emitted per molecule before photobleaching.
<sup>(f)</sup> Calculated time to bleach from a rate of 1000 to 500 photons per sec per molecule.

